# Synergistic Phase Separation of Two Pathways Promotes Integrin Clustering and Integrin Adhesion Complex Formation

**DOI:** 10.1101/2021.08.09.455653

**Authors:** L.B. Case, L. Henry, M.K. Rosen

## Abstract

Integrin adhesion complexes (IACs) are integrin-based plasma membrane-associated compartments where cells sense environmental cues. The physical mechanisms and molecular interactions that mediate nascent IAC formation are unclear. We found that both p130Cas (“Cas”) and Focal adhesion kinase (“FAK”) undergo liquid-liquid phase separation *in vitro* under physiologic conditions. Cas- and FAK- driven phase separation is sufficient to reconstitute kindlin-dependent integrin clustering *in vitro*. *In vitro* condensates and cellular IACs exhibit similar sensitivities to environmental perturbations including changes in temperature and pH. Furthermore, mutations that inhibit or enhance phase separation *in vitro* reduce or increase the number of IACs in cells, respectively. Finally, we find that the Cas and FAK pathways act synergistically to promote phase separation, integrin clustering and IAC formation *in vitro* and in cells. We propose that Cas- and FAK- driven phase separation provides an intracellular trigger for integrin clustering and nascent IAC formation.

## INTRODUCTION

Integrin-mediated adhesion complexes (IACs) are plasma membrane-associated compartments that provide specific adhesion between cells and their surroundings (Case and Waterman, 2015; Chastney et al., 2021). IACs serve as sites of force transmission between the actin cytoskeleton and the extracellular matrix (ECM) to drive tissue morphogenesis, cell movement, and ECM remodeling. Additionally, they serve as signaling hubs where cells sense biochemical and physical cues in their environment to regulate the cell cycle, differentiation and death. Thus, IACs mediate an array of functions involving biochemical and physical interactions between the cell and its environment.

Integrins are heterodimeric transmembrane receptors that specifically bind to ligands within the ECM or to receptors on adjacent cells (Hynes, 2002). Both chains of the α/β heterodimer have a large extracellular domain that imparts ligand specificity and a small cytoplasmic domain (20 – 50 amino acids) that mediates intracellular signaling. Since integrins lack catalytic activity and do not directly bind actin (Hynes, 2002), downstream signaling requires the macromolecular assembly of integrins with cytoplasmic adaptor proteins, signaling molecules, and actin binding proteins to form IACs. Thus, molecular interactions that regulate integrin clustering and initial IAC formation control integrin-dependent signaling (Hotchin and Hall, 1995; Miyamoto et al., 1995; Robertson et al., 2015; Theodosiou et al., 2016).

IACs initially form as puncta (∼120 nm diameter) within the lamellipodia, termed nascent adhesions. Nascent adhesion formation is dependent on ligand binding, integrin conformational activation, and mechanical forces imparted by retrograde actin flow (Changede et al., 2015; Choi et al., 2008). Although nascent adhesion formation and cell spreading are impaired on soft substrates (where mechanical forces are reduced), this can be rescued by inducing the integrin conformational change with Mn^2+^, suggesting that during the initial steps of nascent adhesion assembly force is only required to activate integrins (Oakes et al., 2018). Kindlin and talin are adaptor proteins that directly bind the β integrin cytoplasmic domain (Sun et al., 2019). Kindlin binding is required for integrin clustering and nascent adhesion assembly, although it is not clear how kindlin orchestrates this higher-order assembly (Theodosiou et al., 2016; Ye et al., 2013). Many proteins appear to simultaneously assemble into nascent adhesions, including α5β1 integrin, αVβ3 integrin, talin, kindlin, focal adhesion kinase (FAK), paxillin, and p130Cas (Bachir et al., 2014; Changede et al., 2015; Choi et al., 2008; Donato et al., 2010; Lawson et al., 2012; Yu et al., 2011). After initial formation, nascent adhesions undergo dramatic growth and compositional maturation dependent on talin, increased forces, and bundled actin filaments (Choi et al., 2008; Kuo et al., 2011; Oakes et al., 2012; Schwarz and Gardel, 2012). While IAC maturation is reasonably well understood, the specific molecular interactions that regulate initial nascent adhesion formation are unclear. In this study, we investigate how specific protein-protein interactions contribute to nascent adhesion formation.

Liquid-liquid phase separation driven by weak interactions between multivalent molecules has emerged as an important mechanism that can drive the formation of micron-sized, membraneless cellular compartments, termed “biomolecular condensates” (Banani et al., 2017; Case et al., 2019a; Shin and Brangwynne, 2017). At the plasma membrane, phase separation can promote the assembly of transmembrane proteins and their cytoplasmic binding partners into dynamic, micron-sized clusters (Banjade and Rosen, 2014; Beutel et al., 2019; Case et al., 2019a; Su et al., 2016; Zeng et al., 2016). Phase separation is highly dependent on molecular valence (Li et al., 2012), and increasing the valence of interactions reduces the concentration threshold required for phase separation. Condensates often possess liquid-like material properties, can contain hundreds of molecular constituents, and form over a wide range of molecular stoichiometries (Banani et al., 2017; Case et al., 2019b). Condensate composition can vary dramatically and change rapidly in response to signals (Banani et al., 2016; Markmiller et al., 2018; Youn et al., 2018).

IACs are discrete, micron-sized structures that exhibit many characteristics of phase separated compartments. They are highly enriched in multivalent adaptor proteins whose valence can be modulated by phosphorylation and mechanical force (Pellicena and Miller, 2001; Schiller and Fassler, 2013; Yao et al., 2016), concomitant with changes in IAC size. IACs often exhibit liquid-like material properties such as rapid constituent exchange, (Table S1) (Lavelin et al., 2013; Pasapera et al., 2010; Stutchbury et al., 2017) and the ability to fuse (Berginski et al., 2011; Changede et al., 2015). IACs contain hundreds of different proteins (Schiller and Fassler, 2013), are stoichiometrically undefined (Bachir et al., 2014), and their composition can change dramatically under different cellular conditions (Horton et al., 2015; Kuo et al., 2011; Schiller and Fassler, 2013). Because of their properties and molecular composition, we hypothesized that multivalent oligomerization and phase separation of IAC-associated proteins might contribute to integrin clustering and nascent adhesion assembly.

Here we investigated biochemical interactions among the earliest proteins to arrive at nascent adhesions. We found that several of these undergo liquid-liquid phase separation when combined *in vitro* at physiologic concentrations and under physiologic buffer conditions. Multivalent interactions between phosphorylated p130Cas (“pCas”), Nck and N-WASP are sufficient for phase separation. Focal Adhesion Kinase (“FAK”) also phase separates, and multivalent interactions between FAK and paxillin enhance this behavior. pCas and FAK undergo phase separation synergistically to form droplets that more strongly concentrate kindlin, and consequently integrin. Using a novel experimental platform to reconstitute β1 integrin clustering on supported phospholipid bilayers, we found that kindlin likely plays a central role in nascent adhesion assembly by coupling the pCas and FAK pathways to integrins. *In vitro* condensates and cellular IACs exhibit similar sensitivity to temperature and pH. Further, mutations in Cas or FAK that inhibit phase separation *in vitro* reduce the number IACs in cells, while paxillin mutations that enhance phase separation *in vitro* increase the number of IACs in cells. Finally, we find that the Cas and FAK pathways act synergistically to promote phase separation, integrin clustering and nascent adhesion formation *in vitro* and in cells. We propose that pCas- and FAK- driven phase separation provides an intracellular trigger for integrin clustering and nascent adhesion formation.

## RESULTS

### Multivalent interactions promote phase separation of p130Cas under physiologic conditions

The adaptor protein Cas (p130Cas/BCAR1) is present in nascent adhesions (Donato et al., 2010) and has been implicated in promoting IAC formation (Meenderink et al., 2010). Cas contains an N-terminal SH3 domain, a central disordered “substrate” domain, a Src-binding domain, and a C-terminal Cas-family homology domain (Figure 1 – figure supplement 1) (Meenderink et al., 2010). Within the substrate domain, there are 15 YXXP motifs that, when phosphorylated, can bind to the SH2 domain-containing adaptor proteins, Crk-II and Nck (Iwahara et al., 2004; Pellicena and Miller, 2001; Schlaepfer et al., 1997). In cells, Cas is robustly phosphorylated on 10 of its YXXP motifs, and proper cell migration requires a minimum of four phosphorylation sites (Shin et al., 2004). Crk-II contains two SH3 domains which can bind adaptor proteins such as SOS and C3G that contain multiple proline rich motifs (PRMs) (Birge et al., 2009). Similarly, Nck contains three SH3 domains which can bind adaptor proteins such as N-WASP, SOS and Abl that contain multiple PRMs (Li et al., 2001). In other signaling pathways, phosphorylation of proteins on multiple tyrosine residues and subsequent binding by multivalent adaptor proteins can promote phase separation (Banjade and Rosen, 2014; Kim et al., 2019; Su et al., 2016). For example, multivalent interactions between phosphorylated Nephrin, Nck, and N-WASP or phosphorylated Lat, Grb2 and SOS drive phase separation to form droplets in solution and clusters at membranes (Banjade and Rosen, 2014; Banjade et al., 2015; Kim et al., 2019; Li et al., 2012; Su et al., 2016). Thus, we sought to determine if the multiply-phosphorylated Cas substrate domain could promote phase separation, similar to phosphorylated Nephrin and Lat. Nck has been localized within IACs (Goicoechea et al., 2002; Horton et al., 2015), N-WASP colocalizes with Cas within the lamellipodia where nascent adhesions form (Zhang et al., 2014), and both Nck and N-WASP have been implicated in regulating cell adhesion to fibronectin (Misra et al., 2007; Ruusala et al., 2008). Thus, we chose to use Nck as the SH2/SH3 domain containing adaptor protein and N-WASP as the PRM containing adaptor protein for our studies (Fig. 1a, Table S2). To assess phase separation *in vitro*, we purified recombinant proteins (Figure 1 – figure supplement 2a), phosphorylated Cas on tyrosine to an average of ∼19 sites/molecule (pCas, Figure 1 – figure supplement 2b), and assessed phase separation by both measuring solution turbidity (Absorbance at 350 nm) and visualizing droplets with fluorescence microscopy. Although purified pCas is more highly phosphorylated than in cells, the valency for Nck is likely similar (five of the six Nck-binding motifs are robustly phosphorylated in cells) (Shin et al., 2004). For all *in vitro* experiments, we used buffer containing 50 mM HEPES (pH 7.3), 50 mM KCl, 1 mM TCEP and 0.1% BSA. No crowding agents were used in any experiments. In mammalian cells, the concentration of Nck is ∼200 nM, that of N-WASP (and its homolog, WASP) ranges from 150 nM – 10 µM, and that of Cas is ∼70 nM (Table S3, (Hein et al., 2015; Higgs and Pollard, 2000; Isaac et al., 2010; Roybal et al., 2016)). We combined Nck and N-WASP at physiologic concentrations (200 nM Nck/1 µM N-WASP and 500 nM Nck/1 µM N-WASP) and titrated increasing concentrations of unphosphorylated or phosphorylated Cas. We specifically observed an increase in solution turbidity at physiologic concentrations (10 – 200 nM) of pCas (Fig. 1b). Spinning disk confocal fluorescence microscopy confirmed that the increase in solution turbidity was due to formation of condensed foci at physiologic concentrations (Figure 1 – figure supplement 3). We increased protein concentrations to 1 µM to ensure that the foci were larger than the point spread function of the microscope, revealing that they are spherical (Fig. 1c) and suggesting that they behave as liquid droplets. We measured the partition coefficient (PC) of molecules into the droplets ([Intensity inside droplet/[Intensity in bulk solution]; Fig. 1d, Figure 1 – figure supplement 4, see methods) and performed fluorescence recovery after photobleaching (FRAP) analysis (Figure 1 – figure supplement 5). We found that pCas, Nck and N-WASP rapidly exchanged between droplets and bulk solution (Fig. 1e; Table S4; pCas t_1/2 fast_ = 2 s; pCas t_1/2 slow_ = 39s; Nck t_1/2_ = 8 s; N-WASP t_1/2 fast_ = 1 s; N-WASP t_1/2 slow_ = 37 s), although a fraction of Nck and pCas molecules did not recover in the 100 s timeframe of the experiment, suggesting a slower phase of recovery and/or an immobile fraction (Table S4). In cells, IAC-associated proteins, including Cas, often contain a population of fast exchanging molecules (t_1/2_ < 30 s) as well as an immobile fraction (up to 50% immobile for some proteins; Table S1). Thus, similar to cellular IACs, droplets exhibit liquid-like material properties with some solid-like elements as well. We conclude that interactions between multiply-phosphorylated Cas, Nck, and N-WASP are sufficient to promote liquid-liquid phase separation at physiologic protein concentrations.

**Figure 1.**
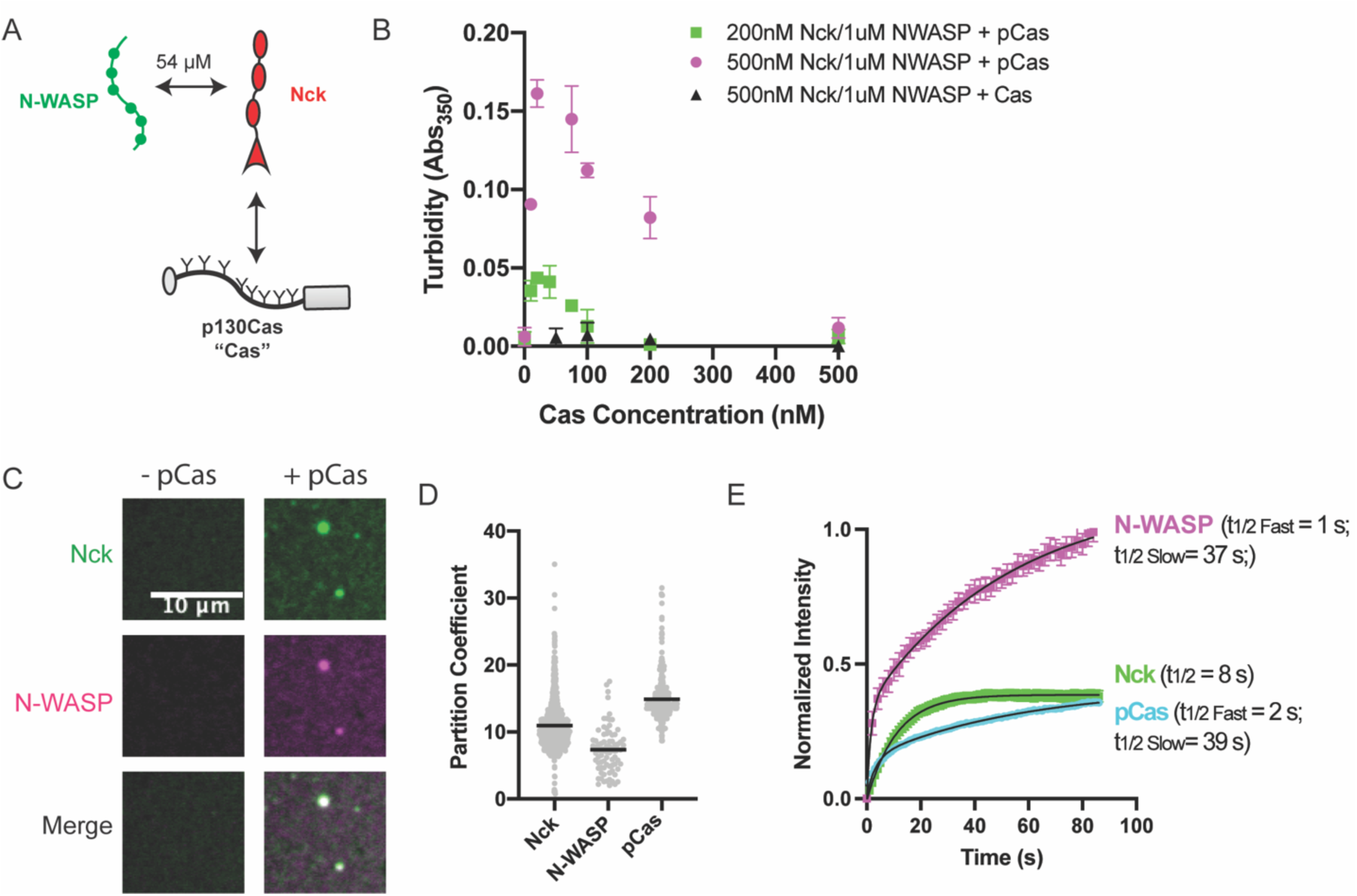
p130Cas, Nck, and N-WASP undergo liquid-liquid phase separation **(A)** Molecular interactions of IAC proteins, K_D_ values indicated where known. Details and references in Table S1. **(B)** Solution turbidity measurements. Nck (200 nM, green; or 500 nM, magenta + black) and N-WASP (1 µM) were combined with increasing concentrations of phosphorylated Cas (pCas, green + magenta) or unphosphorylated Cas (Cas, black). Each point represents the mean ± SEM of three independent measurements. **(C)** Spinning disk confocal fluorescence microscopy images of droplets. Nck (1 μM, 15% Alexa568 labeled) and N-WASP (1 μM, 15% Alexa647 labeled) were combined ± pCas (1 μM, unlabeled). (**D**) Quantification of constituent partitioning into droplets. Each grey point represents an individual measurement, and the mean is written and indicated by black line. Each condition contains at least 75 measurements from 2 or more independent experiments. **(E)** Fluorescence Recovery After Photobleaching (FRAP) measurements of droplets. Droplets formed from 1 μM each of Nck (15% Alexa568 labeled), N-WASP (15% Alexa647 labeled) and pCas (5%-647 labeled). Each point represents the mean ± SEM of at least six independent measurements. Recovery curves were fit with a single exponential (Nck) or biexponential (pCas, N-WASP) model and the fits are overlayed on the graph (black line). Detailed fit information in Table S4. All scalebars = 10 μm.

### FAK and paxillin phase separate under physiologic conditions

Like many condensates, IAC assembly is robust to changes in composition. Although Cas-/- fibroblasts still form IACs, they exhibit decreased adhesion assembly rates (Meenderink et al., 2010). These observations suggest a potential role for Cas in nascent adhesion assembly that is at least partially compensated for by additional proteins (Honda et al., 1998). Thus, we sought to identify additional interactions that may promote integrin clustering at IACs, perhaps through phase separation (Fig. 2a, Table S2). Talin, kindlin, paxillin, and Focal Adhesion Kinase (FAK) regulate nascent adhesion assembly, are present in nascent adhesions, and rapidly exchange between IACs and the cytoplasm (Table S1)(Bachir et al., 2014; Changede et al., 2015; Choi et al., 2008; Meenderink et al., 2010; Swaminathan et al., 2016; Theodosiou et al., 2016). Full length talin is autoinhibited (Dedden et al., 2019), but the talin head domain (talinH) is sufficient to rescue nascent adhesion formation in talin depleted cells (Changede et al., 2015). We expressed and purified recombinant talinH, kindlin, paxillin, and FAK (Figure 1 – figure supplement 2) and screened for phase separation *in vitro* by measuring solution turbidity. We observed an increase in solution turbidity with increasing concentrations of FAK, starting at ∼100 nM, but not with increasing concentrations of talinH, kindlin, or paxillin (Fig 2b, Figure 2 – figure supplement 1a). Note that we purify FAK in buffer containing 300 mM NaCl, preventing self-assembly prior to dilution into the experimental buffer. Using spinning disk confocal fluorescence microscopy, we observed small FAK foci at 40 nM concentration (Figure 2 – figure supplement 2a-b), the upper end of FAK concentrations in mammalian cells (5 - 40 nM) (Brami-Cherrier et al., 2014; Hein et al., 2015) (Table S3). With 1 µM FAK (1% Alexa-647-labeled), we observe larger, micron-scale spherical droplets (Fig. 2c) that exchange molecules with bulk solution as assessed by FRAP (Fig. 2g. Fig. 2 – figure supplement 3a; Table S4; t_1/2 fast_ = 6 s; t_1/2 slow_ = 56 s), although a fraction of molecules (38%) did not recover. Thus, droplets exhibit liquid-like material properties, albeit with some solid-like elements as well. We conclude that FAK undergoes liquid-liquid phase separation *in vitro* under physiologic conditions.

**Figure 2.**
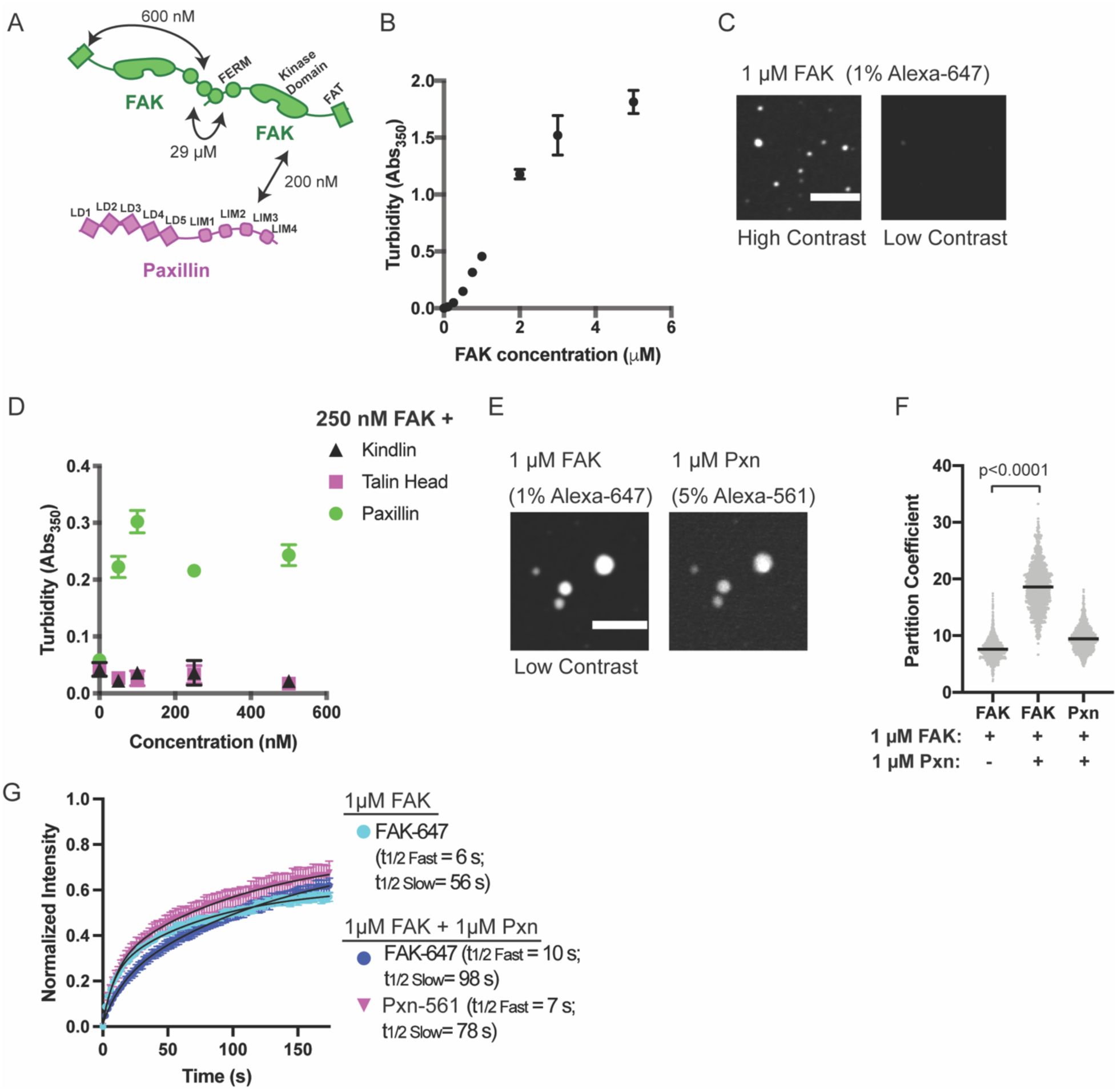
FAK and paxillin undergo liquid-liquid phase separation **(A)** Molecular interactions of IAC proteins, K_D_ values indicated where known. Details and references in Table S1. **(B)** Solution turbidity measurements with increasing concentrations of FAK. **(C)** Spinning disk confocal fluorescence microscopy images of droplets formed from 1 μM FAK (1% Alexa647 labeled). The same image is displayed with two different contrast settings for comparison to panel E. **(D)** Solution turbidity measurements. 250 nM FAK was combined with increasing concentrations of talinH (magenta), kindlin (black), or paxillin (green). **(E)** Spinning disk confocal fluorescence microscopy images of droplets formed from 1 μM FAK (1% Alexa647 labeled) and 1 μM FAK paxillin (5% Alexa546 labeled). For comparison, contrast settings of FAK image are identical to those in 2C, low contrast. **(F)** Quantification of constituent partitioning into droplets. Each condition contains at least 750 measurements from 4 or more independent experiments. Significance tested with student’s t-test. **(G)** Fluorescence Recovery After Photobleaching (FRAP) measurements of droplets. Droplets formed from 1 μM of FAK (1% Alexa647 labeled, cyan) or 1 μM each of FAK (1% Alexa647 labeled, dark blue) and paxillin (5% Alexa546 labeled, magenta). Each point represents the mean ± SEM of at least 15 independent measurements. Recovery curves were fit with a biexponential model and the fits are overlayed on the graph (black line). Detailed fit information in Table S4. In (B) and (D), each point represents the mean ± SEM of three independent measurements. In (F), each grey point represents an individual measurement, and the mean is written and indicated by black line. All scalebars = 10 μm.

#### Paxillin enhances FAK phase separation

Next, we sought to identify additional interactions that might enhance FAK phase separation. FAK can interact directly with paxillin and talin (Lawson et al., 2012; Thomas et al., 1999). Thus, we added increasing concentrations of paxillin, talin, or kindlin to 250 nM FAK. We found that paxillin, but not talinH or kindlin, increased solution turbidity (Fig. 2d). Turbidity of a solution containing 250 nM FAK and paxillin was not increased by addition of talinH or kindlin (Figure 2 – figure supplement 1b). When paxillin (5% Alexa546) and FAK (1% Alexa647) were combined at 1 µM concentration, micron-sized spherical droplets containing both proteins were formed (Fig. 2e). In the presence of equimolar paxillin, FAK enrichment in the droplets increased two-fold (Fig. 2f). Both FAK and paxillin exchange between droplets and bulk solution in FRAP analyses (Fig. 2g, Fig. 2 – figure supplement 3b-c; Table S4); FAK t_1/2 fast_ = 10 s; FAK t_1/2 slow_ = 98 s ; paxillin t_1/2 fast_ = 7 s, paxillin t_1/2 slow_ = 78 s), although FAK recovery is slowed relative to droplets containing FAK alone. Thus, similar to cellular IACs (Table S1), droplets formed with FAK and paxillin exhibit predominantly liquid-like material properties with some solid-like elements. Finally, we found that paxillin enhances droplet formation even at low nanomolar concentrations of FAK and paxillin (Figure 2 – figure supplement 2c-d). We conclude that interactions with paxillin enhance FAK phase separation.

### The pCas and FAK pathways phase separate synergistically

Our results suggest that pCas-dependent and FAK-dependent phase separation can promote higher-order assembly of IAC-associated proteins. Furthermore, pCas directly interacts with both paxillin and FAK (Wisniewska et al., 2005; Zhang et al., 2017), suggesting these two pathways could function together to regulate IAC formation (Fig. 3a, Figure 1 – figure supplement 1, Table S2). Thus, we sought to better understand the relationship between pCas- and FAK- driven phase separation *in vitro*. When we combined 1 µM each of pCas, Nck, N-WASP, FAK and paxillin (“pCas+FAK mix”), we observed a single class of droplets containing all molecules (Figure 3b). Moreover, partitioning of all molecules into droplets increased 2- to 5-fold with the pCas+FAK mix (Fig. 3c vs Figs. 1d and 2f). Next, we combined lower concentrations of proteins (either 250 nM of each or the cytoplasmic concentrations (Table S3)) and measured solution turbidity in buffers with increasing concentrations of salt. Turbidity was observed at salt concentrations up to 100 mM (cytoplasmic protein concentrations) or 150 mM (uniform 250 nM protein concentration), and generally decreased with increasing salt, consistent with phase separation driven by electrostatic interactions (Figure 3 – figure supplement 1). We conclude that the pCas and FAK pathways undergo phase separation synergistically to form droplets that more strongly concentrate cytoplasmic adaptor proteins.

**Figure 3.**
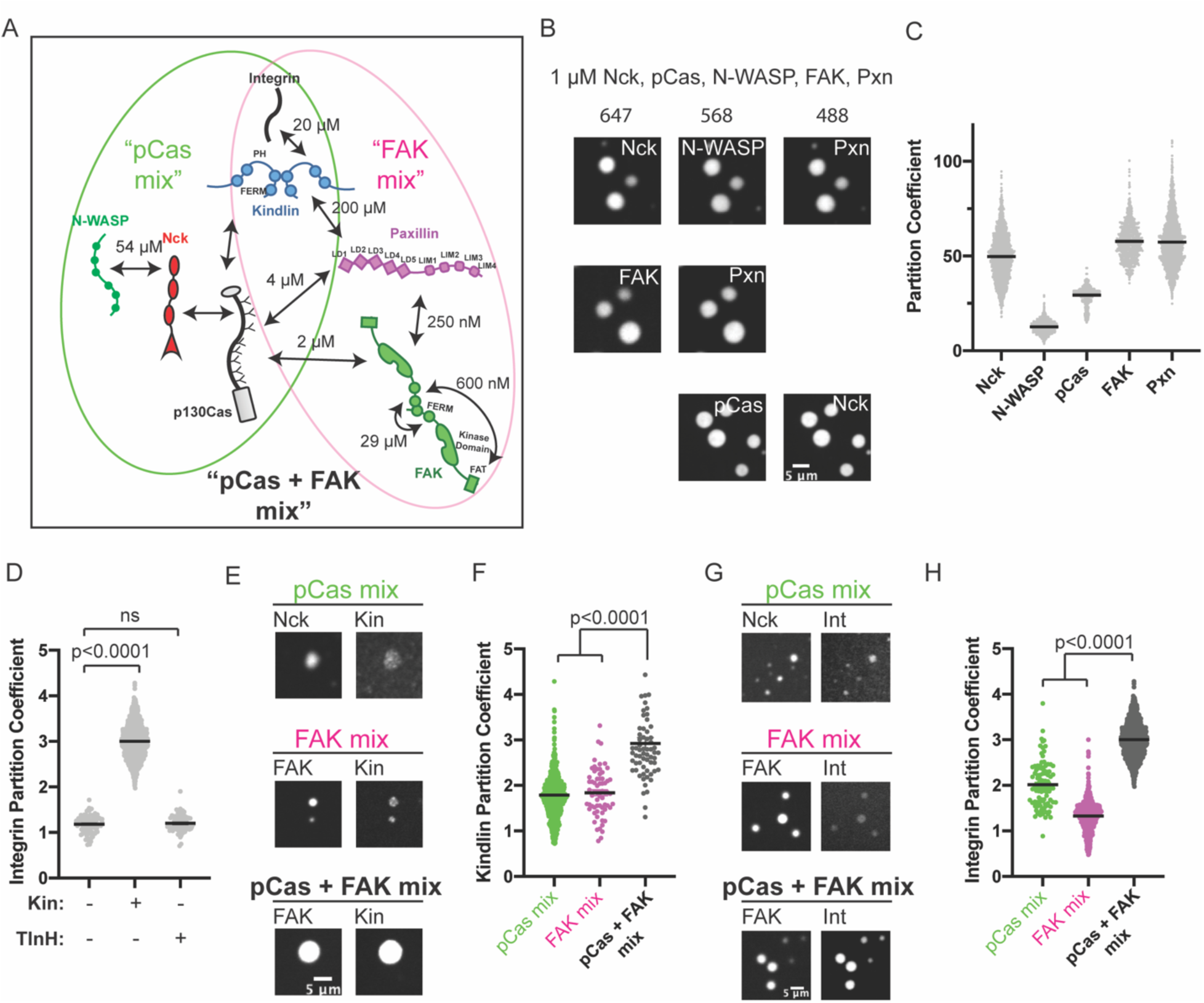
The pCas and FAK pathways phase separate synergistically **(A)** Molecular interactions and known K_D_ values, including between the two pathways. Details and references in Table S1**. (B)** Spinning disk confocal fluorescence microscopy images of droplets formed with the ***pCas + FAK mix***. TOP: 1 μM each of Nck (15% alexa647), N-WASP (15% Alexa568), pCas, FAK, paxillin (15% Alexa488); MIDDLE: 1 μM each of Nck, N-WASP, pCas, FAK (1% Alexa647 labeled), paxillin (1% Alexa488 labeled); BOTTOM: 1 μM each of Nck (15% Alexa488 labeled), N-WASP, pCas (1% Alexa647 labeled), FAK, paxillin; **(C)** Quantification of constituent partitioning into droplets. Each condition contains at least 750 measurements from 4 or more independent experiments. **(D)** Quantification of integrin-GFP partitioning into droplets formed with the ***pCas + FAK mix***. 1 μM each of Nck, N-WASP, pCas, FAK, paxillin and B1 Integrin (15% GFP labeled) with either 1 μM Kin or 1 μM TlnH. Each condition contains at least 70 measurements from 2 or more independent experiments. **(E)** Spinning disk confocal fluorescence microscopy images of droplets. Top: droplets formed with the ***pCas mix*** (1 μM each of Nck (15% Alexa647 labeled), N-WASP, pCas, and kindlin (5% Alexa568 labeled)). Middle: Droplets formed with the ***FAK mix*** (1 μM each of FAK (1% Alexa647 labeled), paxillin, and kindlin (5% Alexa568 labeled)). Bottom: Droplets formed with the ***pCas + FAK mix*** (1 μM each of Nck, N-WASP, pCas, FAK (1% Alexa647 labeled), paxillin, and kindlin (5% Alexa568 labeled)). Contrast settings of kindlin images are matched in all images. **(F)** Quantification of kindlin partitioning. Each condition contains at least 60 measurements from 2 or more independent experiments. **(G)** Spinning disk confocal fluorescence microscopy images of droplets. Top: Droplets formed with the ***pCas mix*** (1 μM each of Nck (15% Alexa647 labeled), N-WASP, pCas, kindlin, and integrin (15% GFP labeled)). Middle: Droplets formed with the ***FAK mix*** (1 μM each of FAK (1% Alexa647 labeled), paxillin, kindlin and integrin (15% GFP labeled)). Bottom: Droplets formed with the ***pCas + FAK mix*** (1 μM each of Nck, N-WASP, pCas, FAK (1% Alexa647 labeled), paxillin, kindlin and integrin (15% GFP labeled)). Contrast settings of integrin images are matched in all images. **(H)** Quantification of integrin partitioning. Each condition contains at least 70 measurements from 2 or more independent experiments.

### Kindlin recruits integrin into pCas+FAK droplets

Next, we sought to determine whether phase separated droplets could recruit the β1 integrin cytoplasmic tail. Droplets formed from 1 µM pCas, Nck, N-WASP, FAK and paxillin did not recruit β1 integrin (PC = 1.2), and this was not changed by addition of 1 µM talinH (PC = 1.2). However, adding 1 µM kindlin, significantly increased integrin partitioning in droplets ((PC = 3.0, Fig. 3d). Thus, kindlin specifically couples phase separation of pCas and FAK to the integrin cytoplasmic tail.

To better understand how kindlin recruits integrin into droplets, we measured kindlin partitioning into droplets formed from the pCas mix, FAK mix, and pCas+FAK mix (Fig. 3 e-f). We found that kindlin weakly partitioned into droplets formed with either the pCas mix or the FAK mix, but its partitioning was significantly increased in droplets formed with the pCas+FAK mix (Fig. 3f). Although kindlin can directly bind paxillin (albeit with low affinity, K_D_ = 200 µM) (Bottcher et al., 2017; Zhu et al., 2019), the protein has not been reported to interact with Cas, Nck, or N-WASP (Dong et al., 2016). Thus, we next sought to determine how kindlin partitions into droplets containing only pCas, Nck and N-WASP (the pCas mix). First, we measured kindlin partitioning into previously described droplets formed by phosphorylated Nephrin, Nck and N-WASP but lacking pCas (Li et al., 2012). Kindlin did not enrich in these droplets (PC = 1), suggesting that kindlin does not interact strongly with pNephrin, Nck, or N-WASP (Figure 3 – figure supplement 2a-b). Thus, we hypothesized that kindlin partitions into droplets containing pCas, Nck and N-WASP by binding pCas. To determine if Kindlin could directly interact with Cas, we performed a pull-down experiment. We found that His-tagged kindlin can pull-down both unphosphorylated and phosphorylated Cas (Figure 3 – figure supplement 2c-d). We conclude that kindlin enriches in droplets formed with the pCas mix through direct interactions with Cas that do not require Cas phosphorylation (Fig. 3a). A low affinity interaction with both Paxillin and Cas is consistent with a smaller partition coefficient.

Next, we measured β1 integrin tail partitioning into droplets formed with the pCas mix, FAK mix, and pCas+FAK mix (all containing kindlin). Similar to kindlin, β1 integrin weakly partitions into droplets formed with either the pCas mix or the FAK mix, but partitioning significantly increases when both pathways are combined to form droplets with the pCas+FAK mix (Fig. 3g-h). We conclude that pCas and FAK undergo phase separation synergistically to form droplets that more strongly concentrate kindlin, and consequently integrin.

#### pCas- and FAK- dependent phase separation synergistically promote integrin clustering on membranes

Since IACs are membrane-associated condensates, we next tested whether pCas- and FAK- dependent phase separation were sufficient to promote integrin clustering on supported phospholipid bilayers. The α and β integrin cytoplasmic tails separate upon activation (Kim et al., 2003; Wegener and Campbell, 2008), and talin and kindlin bind to the latter (Li et al., 2017). Thus, the β integrin cytoplasmic domain is the minimal fragment required to examine talin- or kindlin-dependent integrin clustering. Although the cell surface expression of integrin receptors varies under different conditions, integrin densities between 300 – 1500 molecules/μm^2^ have been observed in cells (Rossier et al., 2012; Wiseman et al., 2004). We attached His_10_-tagged β1 integrin cytoplasmic domain (His-β1) at a density of ∼1000 molecules/μm^2^ on phospholipid bilayers composed of 98% phosphatidylcholine (POPC) plus 2% Ni-NTA lipids (Fig. 4a)(Su et al., 2017). Using Total Internal Reflection Fluorescence (TIRF) microscopy, we confirmed our methodology consistently generated fluid phospholipid bilayers (Figure 4 – figure supplement 1) and that His-β1 was uniformly distributed and rapidly diffusing on the bilayer (Fig. 4b-c “control”).

**Figure 4.**
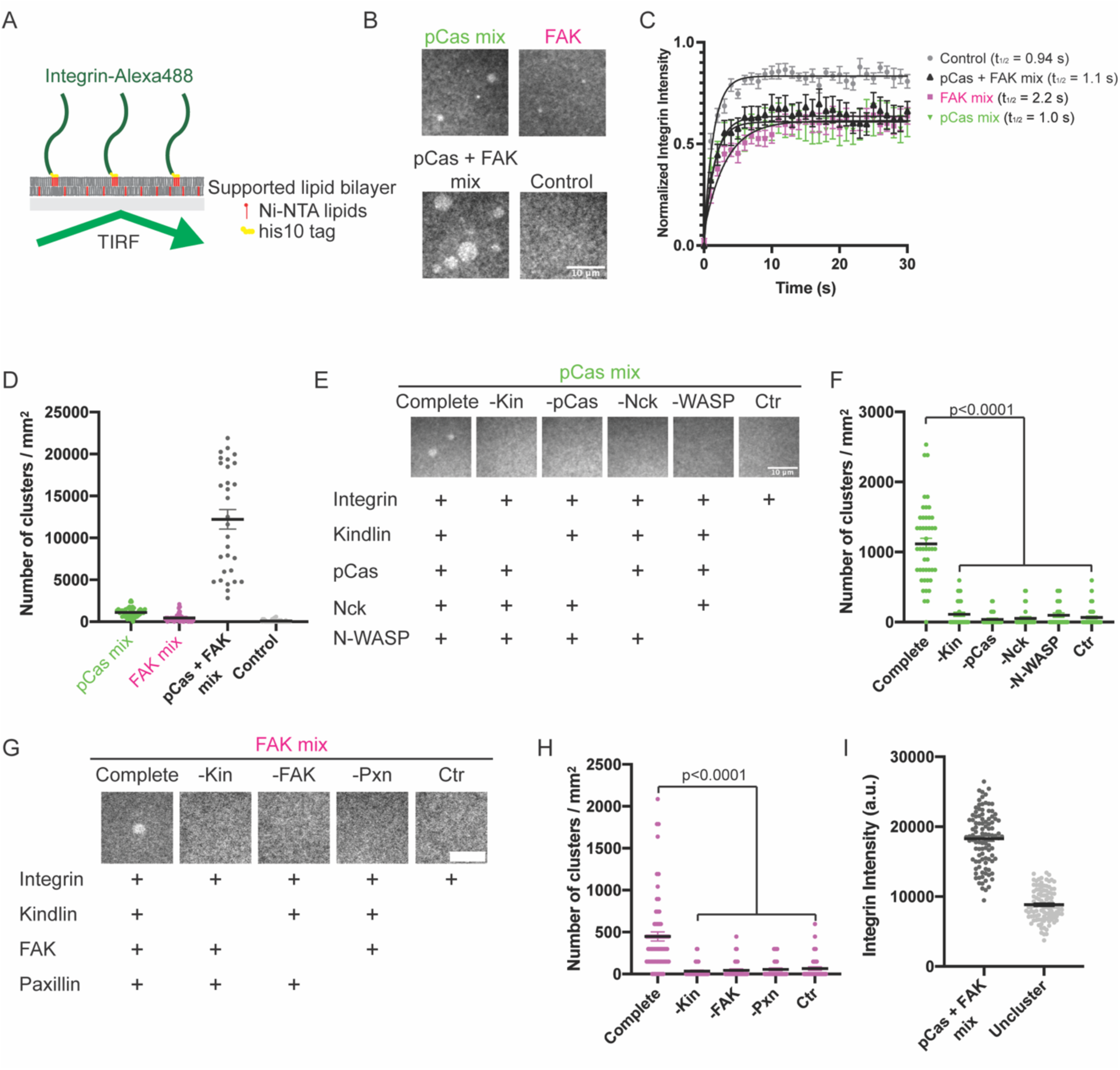
Phase separation is sufficient to reconstitute kindlin-dependent integrin clustering on supported phospholipid bilayers **(A)** Cartoon describing phospholipid bilayer reconstitution. **(B)** Total Internal Reflection Fluorescence (TIRF) microscopy images. Different combinations of proteins (p**Cas mix**: 1 μ M each of Nck, N-WASP, Cas and kindlin; **FAK mix**: 200 nM FAK, and 1 μM each of paxillin and kindlin; p**Cas+FAK mix**: 200 nM FAK and 1 μM each of Nck, N-WASP, Cas, paxillin and kindlin; **Control**: Buffer only) were added to membrane-bound integrin (15% Alexa488 labeled, ∼1000 molecules/μm^2^). (**C**) Fluorescence Recovery After Photobleaching (FRAP) measurements of his-Integrin in clusters. Each point represents the mean ± SEM of at least 12 independent measurements. The t_1/2_ was calculated from a single exponential fit; fit overlayed on graph (black line). **(D)** Quantification of integrin clusters. Each grey point represents a single field of view, black lines represent mean ± SEM. **(E, G)** Total Internal Reflection Fluorescence (TIRF) microscopy images. 1 μM each of the indicated proteins were added to membrane-bound integrin (15% Alexa488, ∼1000 molecules/μm^2^). **(F, H)** Quantification of integrin clusters. Each point represents a single field of view, black lines represent the mean ± SEM. Significance tested by one-way ANOVA followed by a Tukey multiple comparison test. **(I)** Quantification of integrin intensity within clusters formed with the ***pCas + FAK mix***) or in the surrounding unclustered regions (“***Uncluster***”). Each point represents a single measurement, black lines represent mean ± SEM. Each condition contains at least 150 measurements from 2 independent experiments. All scalebars = 10 μm.

After addition of 1 µM each of pCas, Nck, N-WASP and kindlin (“pCas mix”), we observe the formation of micron-sized integrin clusters (∼1,100 clusters/mm^2^, Fig. 4b-d). Similarly, after addition of 200 nM FAK plus 1 μM each of paxillin and kindlin (“FAK mix”), we observe the formation of micron-sized integrin clusters (∼500 clusters/mm^2^, Fig. 4b-d). However, when we combined 200 nM FAK and 1 µM each of pCas, Nck, N-WASP, paxillin, and kindlin (“pCas+FAK mix”), we observe more than a 10-fold increase in the number of integrin clusters (12,200 clusters/mm^2^) compared with either the pCas mix or FAK mix alone (Fig. 4b-d). We conclude that pCas- and FAK- driven phase separation synergistically promote integrin clustering on phospholipid bilayers.

FRAP analysis demonstrates that integrin rapidly exchanges between clusters and the surrounding membrane (Figure 4c), indicating that the clusters are dynamic assemblies. Furthermore, without the complete mixture of proteins in either the pCas mix (Fig. 4e-f) or FAK mix (Fig 4g-h), clusters fail to form. Thus, both kindlin and molecules that promote phase separation are required for integrin clustering on bilayers.

Next, we compared fluorescence intensity of integrin inside clusters formed with the pCas+FAK mix with that in the surrounding bilayer. We found that integrin intensity within pCas+FAK clusters increased an average of 2-fold compared with unclustered integrins (Fig. 4I). In both MEFs and CHO cells plated on fibronectin, α5β1 integrin is 1.3–2 times more concentrated in nascent adhesions compared with the surrounding regions of the membrane (Wiseman et al., 2004). Thus, kindlin coupled to pCas- and FAK- driven phase separation is sufficient to reconstitute physiologically-relevant enrichment of integrin within clusters.

### In vitro droplets and cellular IACs similarly respond to environmental perturbations

Since phase separation is sensitive to solvent conditions, we sought to compare how *in vitro* droplets and cellular IACs responded to changes in environment. First, we determined the effect of temperature on *in vitro* phase separation. We combined 250 nM pCas, Nck, N-WASP, FAK and paxillin, incubated at different temperatures for 30 min, and measured solution turbidity. We observed phase separation at 4°C, 22°C and 37°C, although there was a small but significant decrease in solution turbidity at 4°C (Fig. 5a). The decrease in phase separation at lower temperatures is somewhat unusual, but not unheard of (Jiang et al., 2015; Vrhovski et al., 1997), as for most protein systems phase separation is enhanced as temperature decreases (Nott et al., 2015; Yoshizawa et al., 2018). To determine how IACs respond to a transient change in temperature, cells were plated on fibronectin for 3 hr and then incubated at 4°C, 22°C or 37°C for 10 min, followed by fixation and immunostaining for endogenous paxillin. IACs were observed at all temperatures, but there was a decrease in total adhesion area at 4°C (Fig. 5b-c), mirroring the temperature dependence of the *in vitro* droplets.

**Figure 5.**
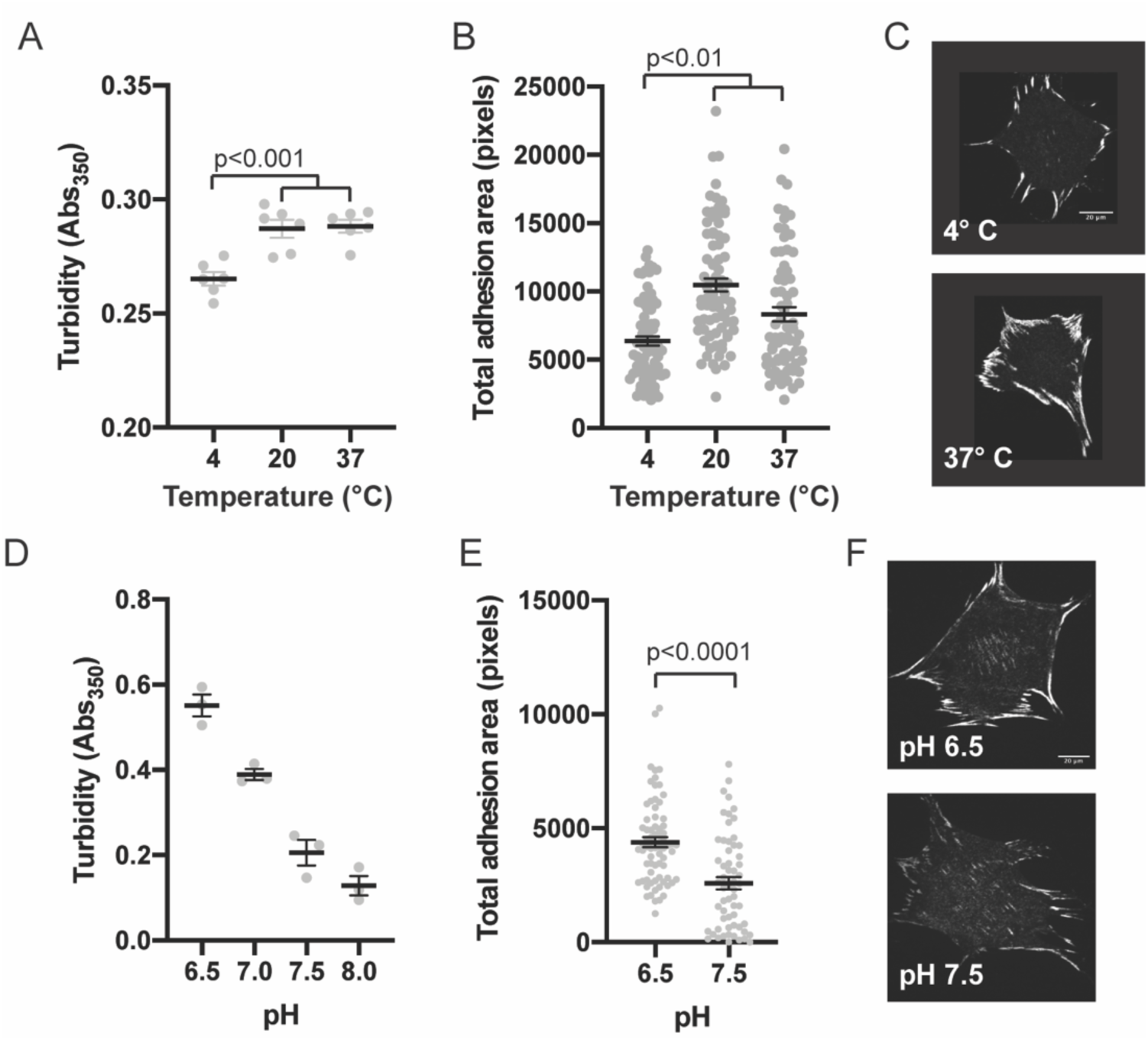
Integrin adhesion complexes are sensitive to solvent perturbations that alter phase separation **(A)** Solution turbidity measurements of solution containing 250 nM each of Nck, N-WASP, pCas, FAK, and Paxillin in buffer containing 50 mM Hepes pH 7.3, 50 mM KCl and 0.1% BSA. Solution incubated at indicated temperatures of 30 min prior to measurements. **(B)** Total adhesion area per cell quantified from spinning disk images of endogenous paxillin. Cells were incubated at indicated temperatures for 10 min prior to fixation and paxillin immunostaining. **(C)** Representative spinning disk confocal microscopy images. **(D)** Turbidity measurements of solution containing 250 nM each of Nck, N-WASP, pCas, FAK, and Paxillin in buffer containing 50 mM KCl, 0.1% BSA and either 50 mM of Hepes or Mes at the indicated pH. **(E)** Total adhesion area per cell quantified from spinning disk confocal microscopy images of endogenous paxillin. Cells were incubated for 10 min prior to fixation and paxillin immunostaining in buffer containing 10 µM nigericin, 10 µM valinomycin, 150 mM NaCl, 50 mM KCl, 1 mM CaCl_2_, 1 mM MgCl_2_, 2 mM Glutamax, and either 50 mM Mes pH 6.5 or 50 mM Hepes pH 7.5. Significance tested with unpaired t-test. **(F)** Representative spinning disk confocal microscopy images. All scale bars = 20 microns.

Next, we determined the effect of pH on phase separation. In cells, IACs are sensitive to intracellular pH (Choi et al., 2013). Mutations in Nhe1 that decrease intracellular pH from a typical resting value of ∼7.5 to 7.0 cause an increase in IAC size and number (Denker and Barber, 2002; Srivastava et al., 2008). We tested the effect of buffer pH on phase separation *in vitro* and found that turbidity of solutions containing 250 nM pCas, Nck, N-WASP, FAK and paxillin increased with increasing acidity. We measured a 2-fold decrease in turbidity between pH 6.5 and pH 7.5 (Fig. 5c). To transiently alter intracellular pH, we treated cells with buffer containing 10 uM Nigericin + valinomycin, ionophores that equilibrate the extracellular and intracellular pH (Triandafillou et al., 2020). We incubated cells with buffers for 10 min followed by fixation and immunostaining. We observed a 2-fold decrease in total adhesion area in cells treated with pH 7.5 compared with pH 6.5 (Fig. 5d-e). Thus, solution pH has parallel effects *in vitro* and in cells, with increasing pH causing decreased pCas+FAK phase separation and decreased total adhesion area. We conclude that solvent perturbations similarly alter phase separation *in vitro* and IACs in cells.

### In vitro droplets and cellular IACs respond similarly to genetic perturbations

#### Preventing Cas phosphorylation reduces the number of IACs in cells

We have identified two distinct sets of molecular interactions, one pCas-dependent and one FAK- dependent, that are sufficient to promote phase separation and drive kindlin-dependent clustering of integrins *in vitro*. Next, we tested whether mutations that perturb Cas or FAK phase separation *in vitro* similarly alter IACs in cells. Cas and FAK are important for both IAC formation and disassembly (Donato et al., 2010; Swaminathan et al., 2016). To distinguish between potentially confounding functions during these opposing processes, we quantified the number of IACs during initial cell spreading when formation dominates. We plated cells on fibronectin-coated glass and allowed them to spread and form IACs for 20 minutes. After fixation and immunostaining for endogenous paxillin, we counted the number of IACs (“number of adhesions,” Figure 6 – figure supplement 1) (Horzum et al., 2014).

We first examined the role of Cas in regulating the number of IACs. We found that Cas-/- MEFs (Figure 6 – figure supplement 2a) formed significantly fewer IACs than WT MEFs, consistent with a potential defect in nascent adhesion formation (Fig. 6a-b). As noted above, unphosphorylated Cas does not phase separate in the presence of Nck and N-WASP (Fig 1b). To parallel this material in cells, all 15 tyrosines within the Cas substrate domain were mutated to phenylalanine (Y15F), which prevents phosphorylation of Cas in cells (Donato et al., 2010). We found that expressing WT Cas in Cas-/- MEFs restored WT numbers of IACs, while expressing Cas Y15F failed to rescue the number of IACs (Fig. 6a-b). Thus, phosphorylation of Cas is required for Cas-dependent regulation of IACs during cell spreading.

**Figure 6.**
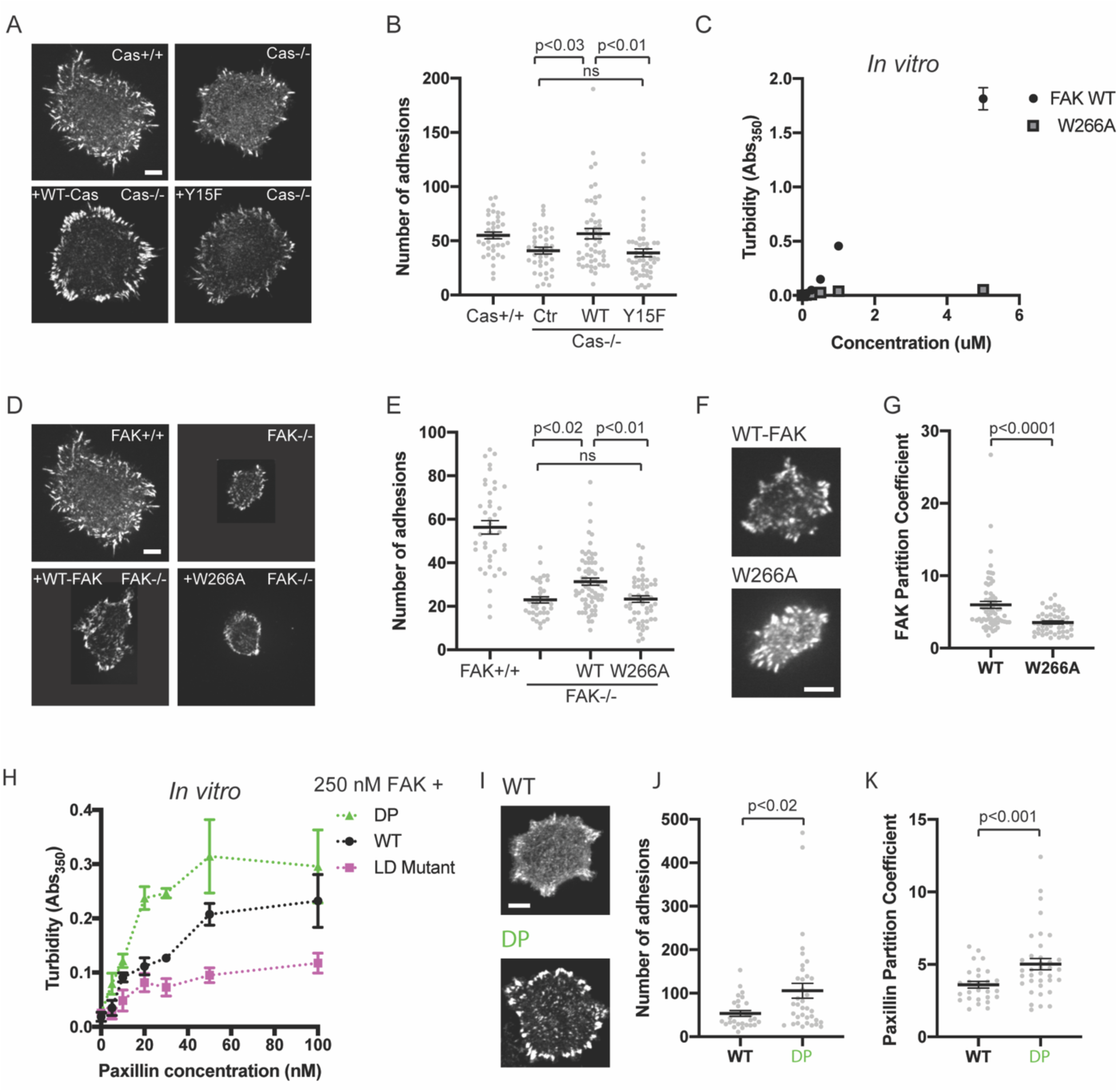
Integrin adhesion complexes are sensitive to genetic perturbations that alter phase separation **(A)** Spinning disk fluorescence microscopy images of MEFs with immunostaining for endogenous paxillin. **(B)** Quantification of number of adhesions **(C)** *In vitro* solution turbidity measurements with increasing concentrations of recombinant WT FAK or W266A FAK (to inhibit dimerization). WT FAK data is duplicated from Fig. 1B for comparison. Each point represents the mean ± SEM of three independent measurements. **(D)** Spinning disk fluorescence microscopy images of MEFs transiently expressing GFP-FAK variants with immunostaining for endogenous paxillin. **(E)** Quantification of number of adhesions. **(F)** Spinning disk fluorescence microscopy images of MEFs transiently expressing GFP-FAK variants. **(G)** Quantification of GFP-FAK partitioning into adhesions (partition coefficient). **(H)** *In vitro* solution turbidity measurements of paxillin variants. 250 nM recombinant FAK was combined with increasing concentrations of recombinant paxillin variants. **(I)** Spinning disk confocal fluorescence microscopy images of MEF cells transiently expressing GFP-paxillin variants. **(J)** Quantification of number of adhesions. **(K)** Quantification of GFP-paxillin partitioning into adhesions (partition coefficient). In (B), (E), (G), (J) and (K) each grey point represents a measurement from one cell, and the mean ± SEM mean is indicated by black lines. Data from at least 35 cells from two or more independent experiments. In (B) and (E) significance tested by one-way ANOVA followed by a Tukey multiple comparison test. In (G), (J) and (K) significance tested by an unpaired t-test. All scalebars = 10 μm.

#### Preventing FAK oligomerization inhibits phase separation and reduces the number of IACs in cells

FAK contains several molecular features that could underlie its ability to phase separate. There are at least two distinct sets of FAK-FAK intermolecular interactions that could promote higher-order oligomerization (Brami-Cherrier et al., 2014). FAK can dimerize through association of two FAK FERM domains, and the FERM:FERM interaction is then further stabilized by an additional interaction between a basic patch in the FERM domain and the C-terminal FAT domain (Fig. 2a). Importantly, these self-interactions can occur in the presence of paxillin, suggesting that higher-order FAK oligomerization is compatible with paxillin binding (Brami-Cherrier et al., 2014). FAK also contains an intrinsically disordered region (IDR) between the kinase domain and FAT domain that has sequence features common to IDRs that phase separate (Figure 6 – figure supplement 3a) (Vernon et al., 2018).

We sought to determine whether perturbing any of these molecular features would reduce FAK phase separation *in vitro*. Pyk2 is closely related to FAK, but the Pyk2 IDR is predicted to have a lower phase separation propensity than the FAK IDR (Figure 6 – figure supplement 3a). Fus is an RNA-binding protein with an IDR that is sufficient for phase separation *in vitro* (Lin et al., 2015). Swapping the FAK IDR for either the Pyk2 IDR or the FUS IDR did not dramatically alter FAK phase separation *in vitro* (Figure 6 – figure supplement 3a-b). Furthermore, a C-terminal fragment containing the IDR and FAT domain did not undergo phase separation (Figure 6 – figure supplement 3c-d). Thus, the IDR is not a dominant driver of FAK phase separation. In contrast, a single point mutation in the FERM domain (W226A) that weakens the FERM-FERM interaction (Brami-Cherrier et al., 2014) was sufficient to dramatically reduce FAK phase separation (Fig. 6c). Furthermore, titrating the FAK C-terminus inhibits phase separation of the full-length protein, suggesting that the FERM-FAT interaction is also important for phase separation (Figure 6 – figure supplement 3e). We conclude that FAK phase separation is primarily driven by higher-order oligomerization (Figure 6 – figure supplement 3f).

We next examined the role of FAK in regulating the number of IACs during cell spreading. We found that FAK-/- MEFs (Figure 6 – figure supplement 2b) formed significantly fewer IACs and were noticeably smaller than WT MEFs, consistent with the role of FAK in nascent adhesion assembly (Fig. 6d-e)(Swaminathan et al., 2016). While FAK-/- MEFs expressing GFP-WT FAK remained noticeably smaller than WT MEFs, GFP-WT FAK still partially rescued the number of IACs. In contrast, FAK-/- MEFS expressing GFP-FAK W266A had no change in the number of IACs compared with FAK-/- MEFs. Thus, FAK oligomerization is required for FAK-dependent regulation of IACs during cell spreading. We next measured FAK partitioning into IACs ([Intensity inside adhesions]/[Intensity in cytoplasm]). We found that the W266A mutation causes a 50% reduction in FAK partitioning (Fig. 6f-g). While W266A FAK cannot drive phase separation, it may still partition into IACs through additional protein-protein interactions, for example binding of the FAK proline-rich motifs to the Cas SH3 domain (Figure 1 – figure supplement 1). In the “scaffold/client” description of condensate composition, the mutation converts FAK from behaving more scaffold-like to more client-like (Banani et al., 2016; Ditlev et al., 2018; Xing et al., 2020). Thus, a decrease in FAK partitioning is consistent with loss of FAK- dependent phase separation. Together, these data demonstrate that impairing the phase separation of either Cas or FAK corelates with a partial reduction in the number of IACs observed after 20 minutes of cell spreading.

#### Increasing paxillin valence increases the number of IACs in cells

Next, we sought to identify mutations that might increase phase separation at IACs. Since interactions between paxillin and FAK enhance FAK-dependent phase separation *in vitro* (Fig. 2d-f), we sought to understand the molecular basis of this enhancement. The FAK C-terminal FAT domain contains two distinct binding sites for at least two paxillin LD motifs (LD2 and LD4)(Gao et al., 2004; Scheswohl et al., 2008; Thomas et al., 1999). To test if multivalent interaction between paxillin and FAK were necessary to enhance phase separation, we engineered paxillin mutants (Figure 6 – figure supplement 4). To weaken paxillin-FAK binding, we mutated a key Asp residue in each of the five LD motifs to reduce the affinity for FAK (“Paxillin LD mutant”)(Thomas et al., 1999). To increase paxillin valence, we duplicated the N-terminus (Residues 1-321) to double the number of LD motifs (“Double Paxillin”, DP). We added increasing concentrations of each paxillin mutant to 250 nM FAK and measured solution turbidity (Fig. 6h). We found that the paxillin LD mutant enhances FAK phase separation less than WT, while the DP mutant acts more strongly than WT. We conclude that multivalent interactions between paxillin LD motifs and the FAK C-term FAT domain enhance FAK phase separation.

Next, we tested whether increasing paxillin valence could alter the number of IACs formed during cell spreading. We expressed GFP-WT paxillin or GFP-DP paxillin in WT MEFs. Cells expressing DP paxillin formed twice as many IACs as cells expressing equal levels of WT paxillin (Fig. 6i-j). Thus, doubling the number of paxillin LD motifs is sufficient to increase the number of IACs. Furthermore, we found that DP paxillin was more strongly partitioned into IACs compared with WT paxillin (Fig. 6k), consistent with an increase in paxillin-dependent phase separation. Thus, enhancing paxillin-dependent phase separation corelates with an increase in the number of IACs. Together, these experiments demonstrate three distinct genetic perturbations that similarly alter phase separation behavior *in vitro* and IAC number in cells.

### pCas and FAK synergistically promote nascent adhesion assembly

Since our *in vitro* data demonstrate that pCas and FAK synergistically promote phase separation and kindlin-dependent integrin clustering, we sought to determine if FAK and Cas act synergistically to promote nascent adhesion formation in cells. To assess nascent adhesion assembly more specifically during cell spreading, we fixed cells after only 5 minutes of spreading on fibronection, immunostained for endogenous paxillin, and imaged cells with TIRF. At this 5- minute time point, cells were still predominantly undergoing isotropic cell spreading and IACs remained small and un-elongated. To simultaneously reduce Cas and FAK protein levels in cells, we first used siRNA to knock down Cas in WT or FAK-/- MEFs (Fig. 7- figure supplement 1a).

We found that the number of nascent adhesions formed in 5-minutes was significantly reduced in cells expressing Cas siRNA, while FAK-/- cells did not significantly differ from FAK+/+ cells (Fig. 7-figure supplement 1b-c). However, western blot analysis showed that Cas protein levels are elevated in FAK-/- MEFs (Fig. 7- figure supplement 1a) and can be returned to WT levels with the re-expression of WT-FAK-GFP (Figure 6 – figure supplement 2b). Additionally, FAK-/- MEFs were more resistant to Cas siRNA (Fig. 7-figure supplement 1a). While this genetic compensation supports our biochemical evidence suggesting that FAK and Cas may be functionally linked, this approach was not sufficient to simultaneously remove FAK and Cas protein from cells.

Next, we tried an alternative approach to remove FAK and Cas protein from cells by using siRNA to knock down FAK in WT or Cas-/- MEFs. Western blot analysis showed this strategy was more effective at achieving low protein levels of both Cas and FAK (Fig. 7a). Using this approach, we found that loss of either Cas or FAK partially reduces the number of nascent adhesions compared with WT MEFs (Fig. 7b-c). Simultaneous loss of both Cas and FAK causes a further reduction in the number of nascent adhesions compared with the loss of either protein alone (Fig. 7b-c). When both Cas and FAK are absent, cells form few nascent adhesions and fail to spread in 5 minutes. Fixation and immunostaining of cells after 20 minutes of spreading showed similar results (Fig. 7 – figure supplement 2), suggesting that decreased nascent adhesion assembly in the absence of Cas and FAK leads to significantly fewer IACs and reduced cell spreading even after 20 minutes.

**Figure 7.**
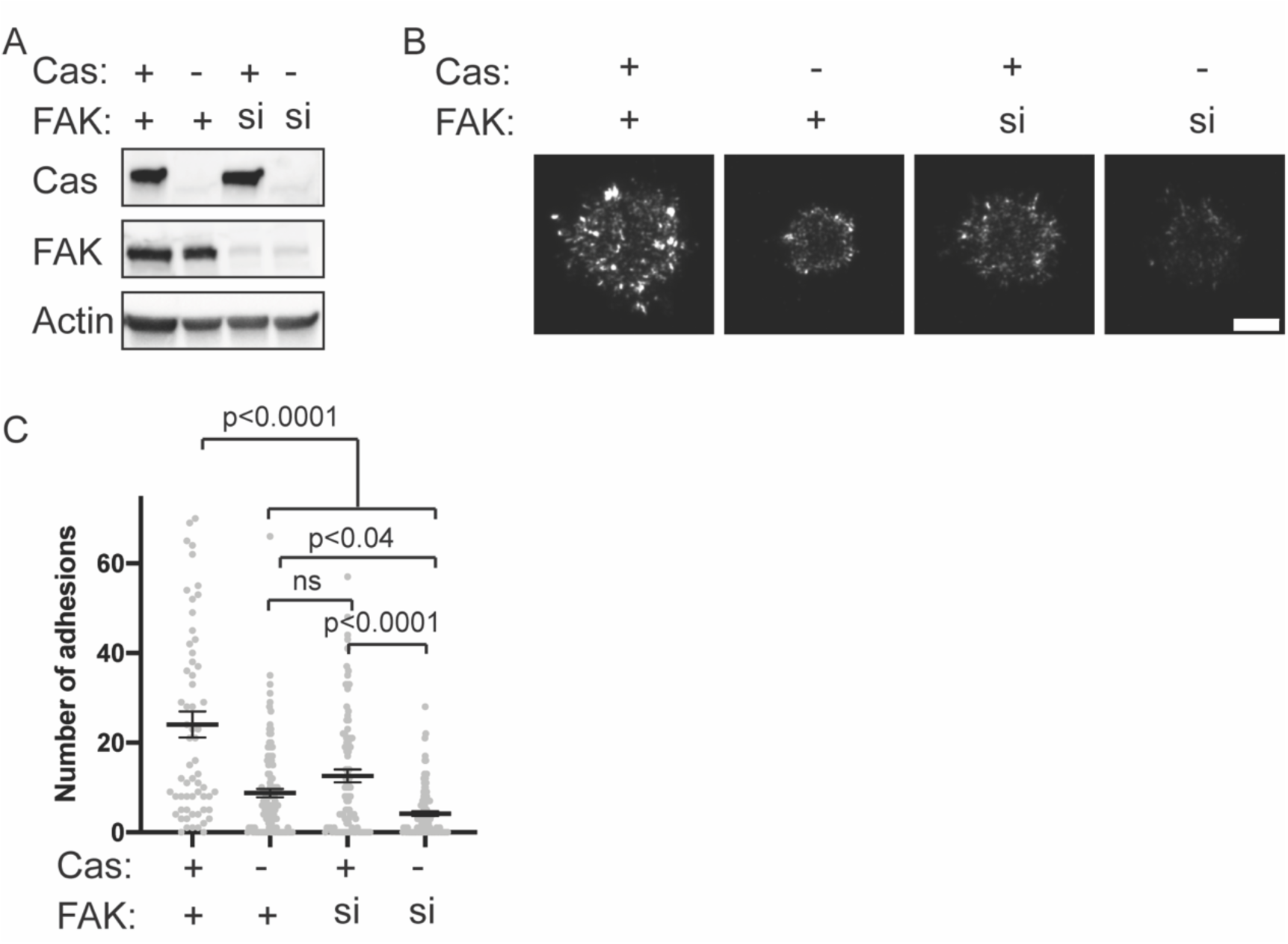
Cas and FAK synergistically promote nascent adhesion assembly during cell spreading **(A)** Western blot analysis of FAK knockdown. Cas^+/+^ or Cas ^-/-^ MEFs were treated with nontargeting (+) or FAK (si) siRNA for 48 hours and lysates were blotted with Cas, FAK or actin antibodies. Original raw files of blots in Figure7-source data. **(B)** Total Internal Reflection Fluorescence (TIRF) microscopy images of MEFs fixed after 5 minutes of spreading with immunostaining for endogenous paxillin. Scalebar = 5 micron. **(D)** Quantification of number of adhesions. Each grey point represents a measurement from one cell, and the mean ± SEM mean is indicated by black lines. Data from at least 60 cells from 2 or more independent experiments. Significance tested by one-way ANOVA followed by a Tukey multiple comparison test.

Together, these data suggest that Cas and FAK have functionally linked roles in promoting nascent adhesion assembly. Consistent with the *in vitro* reconstitution, maximal nascent adhesion assembly occurs in cells expressing both Cas and FAK. Furthermore, genetic studies have found that Cas and FAK are synthetic lethal in *Drosophila* (Tikhmyanova et al., 2010). Neither the FAK mutant nor the Cas mutant flies have a reported phenotype but combining the Cas and FAK mutations is 100% embryonic lethal with a phenotype similar to the β integrin mutant. Our data suggest that the synthetic lethality in flies may reflect the cooperativity in phase separation, integrin clustering, and nascent adhesion assembly that we have seen biochemically and in cultured cells.

## DISCUSSION

Combining purified recombinant proteins on phospholipid bilayers, we successfully developed an *in vitro* reconstitution of nascent adhesions. We found that a mixture of seven proteins is sufficient to reconstitute integrin clustering on phospholipid bilayers and recapitulate the integrin enrichment observed within nascent adhesions in cells. These proteins assemble through phase separation at physiologic concentrations and buffer conditions. This phase separation is necessary and sufficient for kindlin-dependent integrin clustering on phospholipid bilayers. Our data suggest that kindlin plays a central role in nascent adhesion assembly by coupling the Cas and FAK pathways to integrins. We find that both environmental perturbations and genetic perturbations can similarly alter phase separation *in vitro* and the number of IACs in cells. Finally, we find that pCas-dependent and FAK-dependent phase separation synergistically promote integrin clustering on phospholipid bilayers, and that while Cas and FAK can independently promote nascent adhesion assembly, maximum assembly occurs in cells expressing both proteins. Together, these data lead to a model wherein multivalency-driven phase separation of two connected pathways—one based on pCas and the other based on FAK, both coupled to integrins through kindlin—underlies formation of nascent adhesions in cells.

Although kindlin is required for robust IAC formation (Theodosiou et al., 2016), the specific role of kindlin in nascent adhesion formation has remained enigmatic. We find that kindlin is required to couple Cas- and FAK- dependent phase separation to integrins through direct interactions with Cas and paxillin. Observations from our biochemical reconstitution are consistent with previous observations that integrin associates with kindlin and not talin during nascent adhesion assembly (Bachir et al., 2014), that kindlin does not regulate integrin affinity but rather promotes integrin clustering (Ye et al., 2013), and that the kindlin-paxillin interaction is required for efficient nascent adhesion formation in cells (Theodosiou et al., 2016).

Although protein-driven phase separation is consistent with many features of cellular IACs, phase separation likely cooperates with additional factors to regulate nascent adhesion formation in cells. Ligand binding to the extracellular domain of integrin, talin binding to the cytoplasmic domain of integrin, and retrograde actin flow are all required for nascent adhesion formation in cells. Our data suggest that while the ECM-integrin-talin-actin linkage may be critical for integrin conformational activation, it may not be required for subsequent macromolecular assembly of adaptor proteins on the integrin cytoplasmic tail. Furthermore, many IAC-associated proteins can interact with negatively charged phosphoinositides. FAK and N-WASP preferentially bind PI(4,5)P_2_ via basic amino acids (Goni et al., 2014; Papayannopoulos et al., 2005), while kindlin preferentially binds PI(3,4,5)P_3_ via its PH domain (Liu et al., 2011). We find that phosphoinositides are not necessary to reconstitute clusters on phospholipid bilayers that resemble nascent adhesions. Consistent with our observations, reduction of PI(4,5)P_2_ synthesis at IACs in cells leads to the formation of small IACs that contain paxillin and kindlin, but lack talin and vinculin (Legate et al., 2011). Furthermore, talin has been shown to directly interact with and activate PIP5K1C (Di Paolo et al., 2002), and PIP4K2A was identified as a constituent of IACs whose local concentration potentially changes during adhesion assembly and growth (Horton et al., 2015). Thus, PI(4,5)P_2_ may be locally generated during IAC formation or maturation, and additional interactions between phosphoinositides and cytoplasmic adaptor proteins could be important to reduce the threshold concentration of phase separation during nascent adhesion assembly or subsequent adhesion maturation and growth (Mitchison, 2020). Relatedly, coupling between phase separation of lipids and phase separation of membrane-associated proteins was recently described *in vitro* for reconstituted clusters of T cell receptor signaling proteins (Chung et al., 2021). Extracellular ligand binding(Changede et al., 2015), steric exclusion(Paszek et al., 2009), differential lipid composition (Gaus et al., 2006; Sharma et al., 2004; Son et al., 2017), and the actomyosin cytoskeleton(Changede et al., 2015; Kalappurakkal et al., 2019) likely function together with pCas- and FAK- dependent phase separation to regulate different aspects of integrin activation, nascent adhesion formation and adhesion maturation.

We have focused our investigations here on a minimal set of proteins that are known to assemble at integrins coincident with the initiation of nascent adhesion assembly and which play important functional roles in generating the cellular structures. Yet phase separation at mammalian IACs, particularly during maturation where many more proteins arrive at the structures, likely involves even more proteins. Of the 60 proteins consistently found in proteomic analysis of IACs, 27 have three or more repeated modular binding domains (Table S5) (Horton et al., 2015). The multivalent nature of these proteins suggests that they could contribute to higher order assembly and phase separation within IACs. As IAC composition changes during maturation, different multivalent proteins become enriched (Horton et al., 2015; Kuo et al., 2011; Schiller and Fassler, 2013). Future work will be needed to understand how the changing composition might alter the physical properties of the condensed IAC phase and whether additional multivalent proteins become more important at different stages of IAC formation and maturation.

Mechanical forces play a critical role in regulating IAC composition and function. During nascent adhesion assembly, forces are simply required to promote integrin activation (Oakes et al., 2018). However, subsequent IAC growth and compositional maturation are dependent on forces transmitted across IACs between the extracellular matrix and the actin cytoskeleton (Choi et al., 2008; Kuo et al., 2011; Oakes et al., 2012; Schwarz and Gardel, 2012). Future work will be required to understand how the actin cytoskeleton interacts with and potentially reorganizes the condensed IAC phase. In other systems, actin can regulate condensates by acting as a physical barrier to prevent fusion (Feric et al., 2016), by providing mechanical force to promote movement across the membrane surface (Ditlev et al., 2019; Kim et al., 2019), or by acting as a scaffold along which condensates can wet or assemble (Case et al., 2019b; Ditlev et al., 2019; Su et al., 2016). Understanding how forces are transmitted across the condensed IAC phase will require an understanding of the emergent physical properties of the phase. *In vitro* reconstitution will be a useful tool to directly address these questions.

Mature IACs (called focal adhesions) are organized into vertical layers, with paxillin and FAK localized near the membrane and actin and actin binding proteins localized > 50 nm above the membrane (Kanchanawong et al., 2010). At mature focal adhesions, the plasma membrane and actin cytoskeleton provide an inherent polarity; the membrane is physically connected to the ECM through integrins on one side and actin generates forces on the other. These forces are transmitted across the components of the mature focal adhesion. While nascent adhesions may behave like an isotropic liquid condensate, the actin cytoskeleton could induce molecular reorganizations to produce order in the direction perpendicular to the membrane and generate a state more akin to a liquid crystal. While it is not known when this layered organization emerges during IAC assembly and maturation, talin plays a critical role in controlling the distance between actin and the plasma membrane (Liu et al., 2015). Additionally, talin undergoes force dependent conformational changes that extend the protein and expose additional vinculin binding sites (del Rio et al., 2009). Thus, force-dependent talin extension likely plays an important role in patterning the mature focal adhesion. Many other condensates interact with actin, and the layered organization of mature focal adhesions may be more common than previously appreciated (Beutel et al., 2019; Bresler et al., 2004; Zeng et al., 2016).

In conclusion, we have reconstituted minimal macromolecular integrin clusters that have similar composition and features to cellular nascent adhesions. Our biochemical and cellular data provide evidence that Cas- and FAK- dependent phase separation promotes integrin clustering and nascent adhesion assembly. A phase separation model is consistent with many well-documented characteristics of IACs. In other systems, phase separation of receptors enhances downstream signaling (Case et al., 2019b; Huang et al., 2019; Su et al., 2016). Thus, phase separation may provide an important framework for understanding the regulation of signaling and force transmission at IACs.

## ACKNOWLEDGEMENTS

We thank Steve Hanks (Vanderbilt) and Larisa Ryzhova (Maine Medical Center) for sharing Cas- /- cell lines, Robert Liddington (Sanford Burnham Institute) for sharing talin and FAK DNA plasmids, W. Todd Miller (Stony Brook University) for sharing p130Cas DNA plasmid, Jun Qin (Cleveland Clinic) for sharing kindlin DNA plasmid, and David Corey and Jiaxin Hu (UT Southwestern Medical Center) for use of their Cary 100 UV-Visible spectrophotometer and Peltier thermal controller. L.B.C. was a Robert Black Fellow of the Damon Runyon Cancer Research Foundation (DRG-2249-16). This work was also supported by the Howard Hughes Medical Institute and the Welch Foundation (grant I-1544).

## AUTHOR CONTRIBUTIONS

L.B.C. and L.H. prepared reagents. L.B.C. performed experiments and analyzed data. L.B.C. and M.K.R. wrote the manuscript.

---

**Figure 1 – figure supplement 1.**
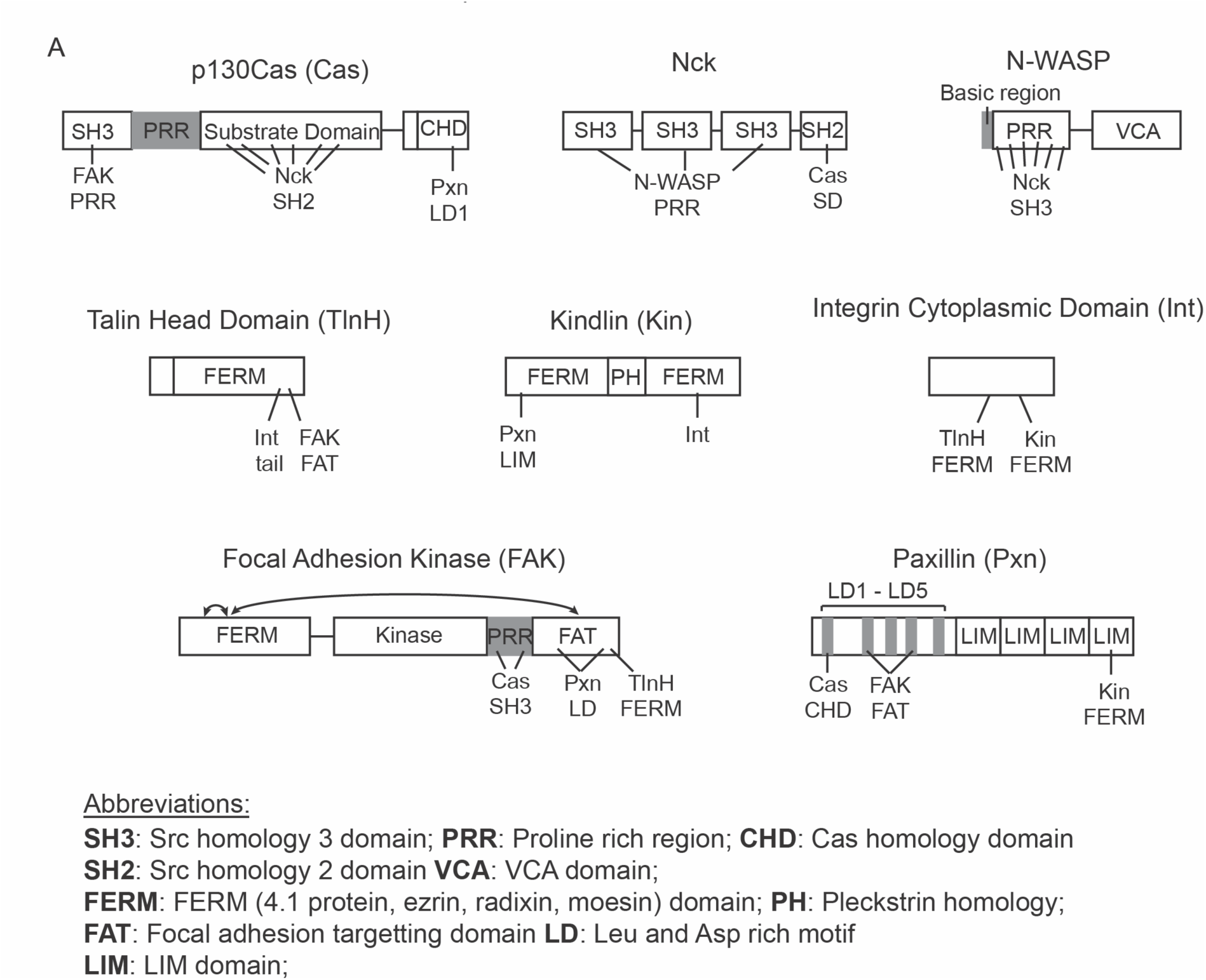
Domain organization of proteins used in this study. Relevant protein interactions are indicated with lines, number of lines corresponds to number of identified binding sites.

**Figure 1 – figure supplement 2.**
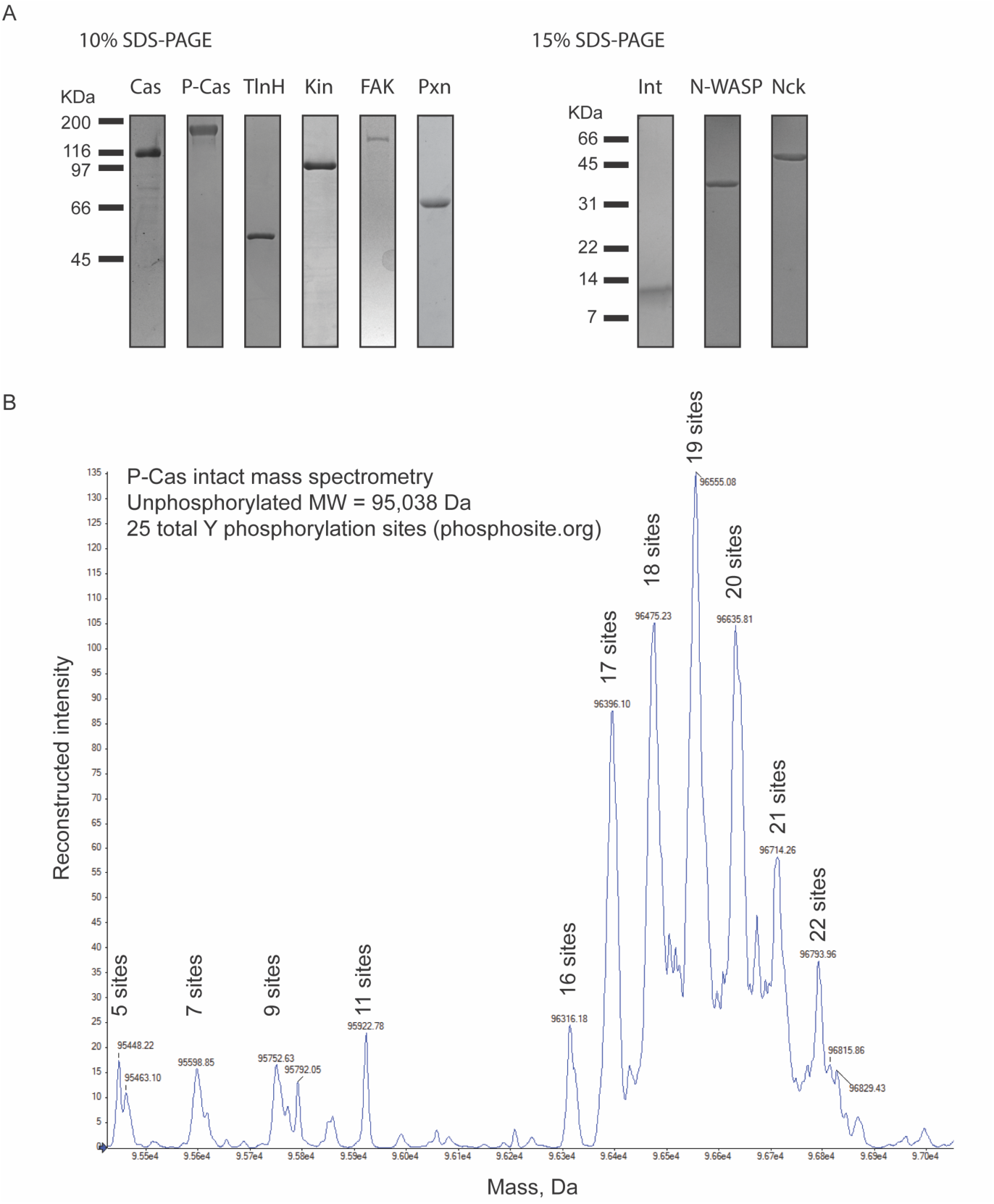
Purification of recombinant integrin adhesion complex proteins **(A)** 0.5 µg of purified protein was run on SDS-PAGE gels and stained with Coomassie Blue. Proteins include unphosphorylated p130Cas (Cas), phosphorylated p130Cas (P-Cas), Talin Head Domain (TlnH), Kindlin (Kin), FAK, Paxillin (Pxn), the cytsoplasmic tail of β1 Integrin (Int), Nck, and N-WASP. Original raw files of gels in Figure1-figure supplement2-source data. **(B)** Phosphorylated p130Cas (P-Cas) was analyzed with mass spectrometry to characterize the degree of phosphorylation. Number of phosphorylation sites of each peak is indicated (assuming 79.98 Da per phosphorylation).

**Figure 1 – figure supplement 3.**
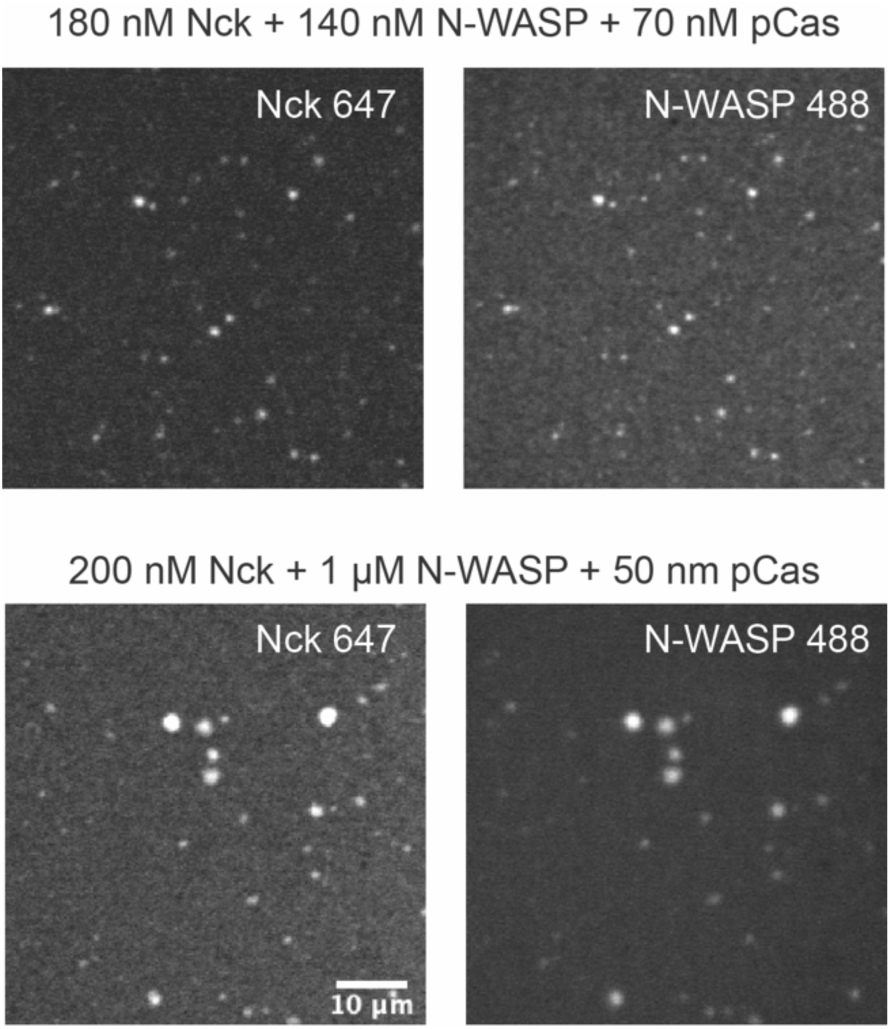
Droplets form with physiological protein concentrations Spinning disk fluorescence microscopy images of droplets. TOP: 180 nM Nck (15% Alexa647), 140 nM N-WASP (15% Alexa488) and 70 nM phosphorylated p130Cas (pCas). BOTTOM: 200 nM Nck (15% Alexa647), 1 µM N-WASP (15% Alexa488) and 50 nM pCas. Scale bar = 10 µm.

**Figure 1 – figure supplement 4.**
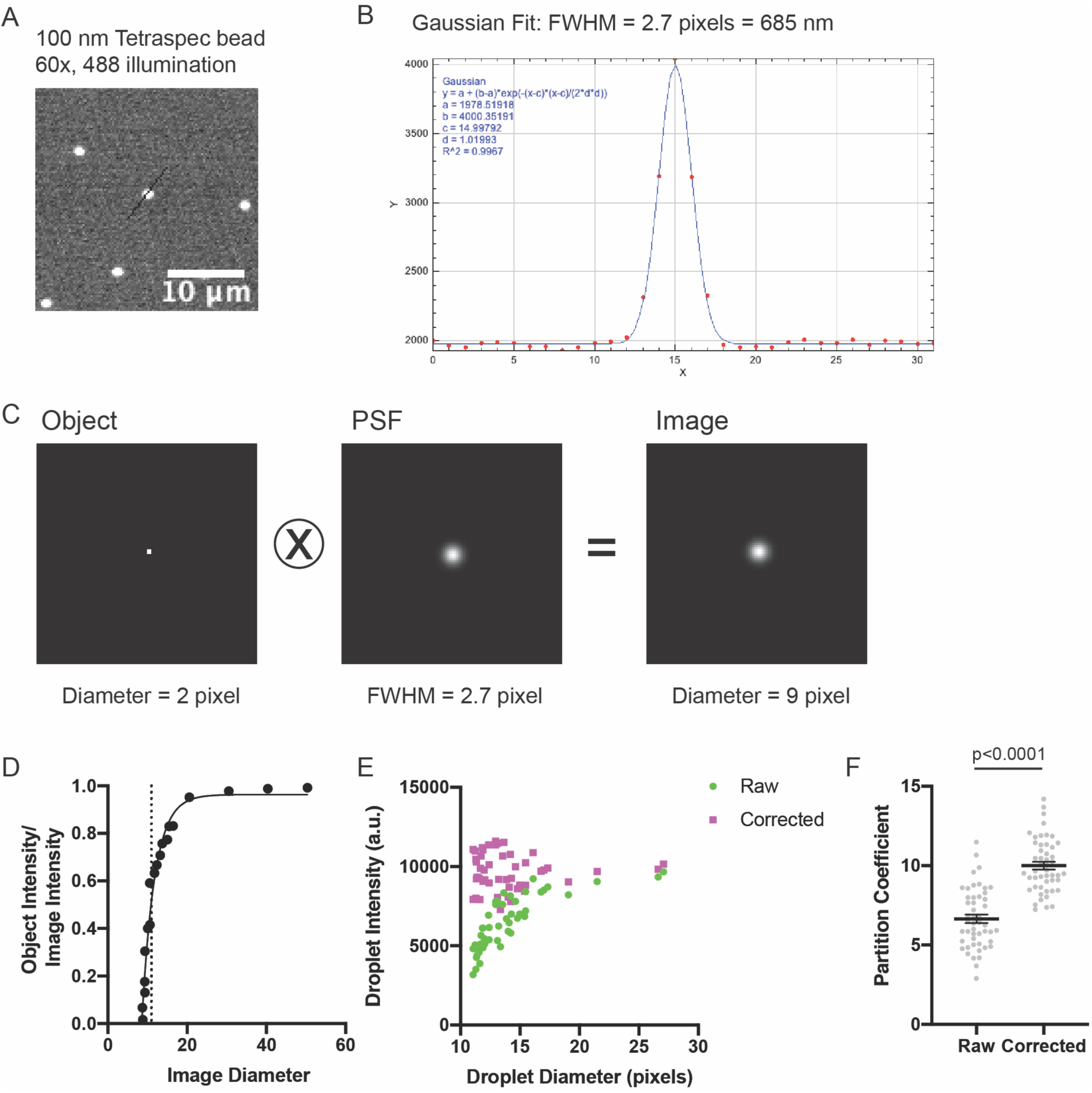
Measuring the point spread function (PSF) **(A)** Spinning disk fluorescence microscopy image of 100 nm Tetraspec bead. **(B)** Linescan of intensity with overlaid Gaussian fit (black line in (A)). **(C)** 2 pixel diameter circle (Object), calculated PSF (PSF), and resulting convolved image (Image). **(D)** Graph of apparent image diameter versus (Object Intensity/Image Intensity). Dotted line shows diameter cutoff below which we cannot accurately measure object intensities. Small objects (< 20 pixel diameter) have diluted apparent intensities. **(E-F)** Comparison of droplet intensity measurements with and without correction. Droplets were formed with 1 µM each of Nck (15% Alexa488), N-WASP, and pCas.

**Figure 1 – figure supplement 5.**
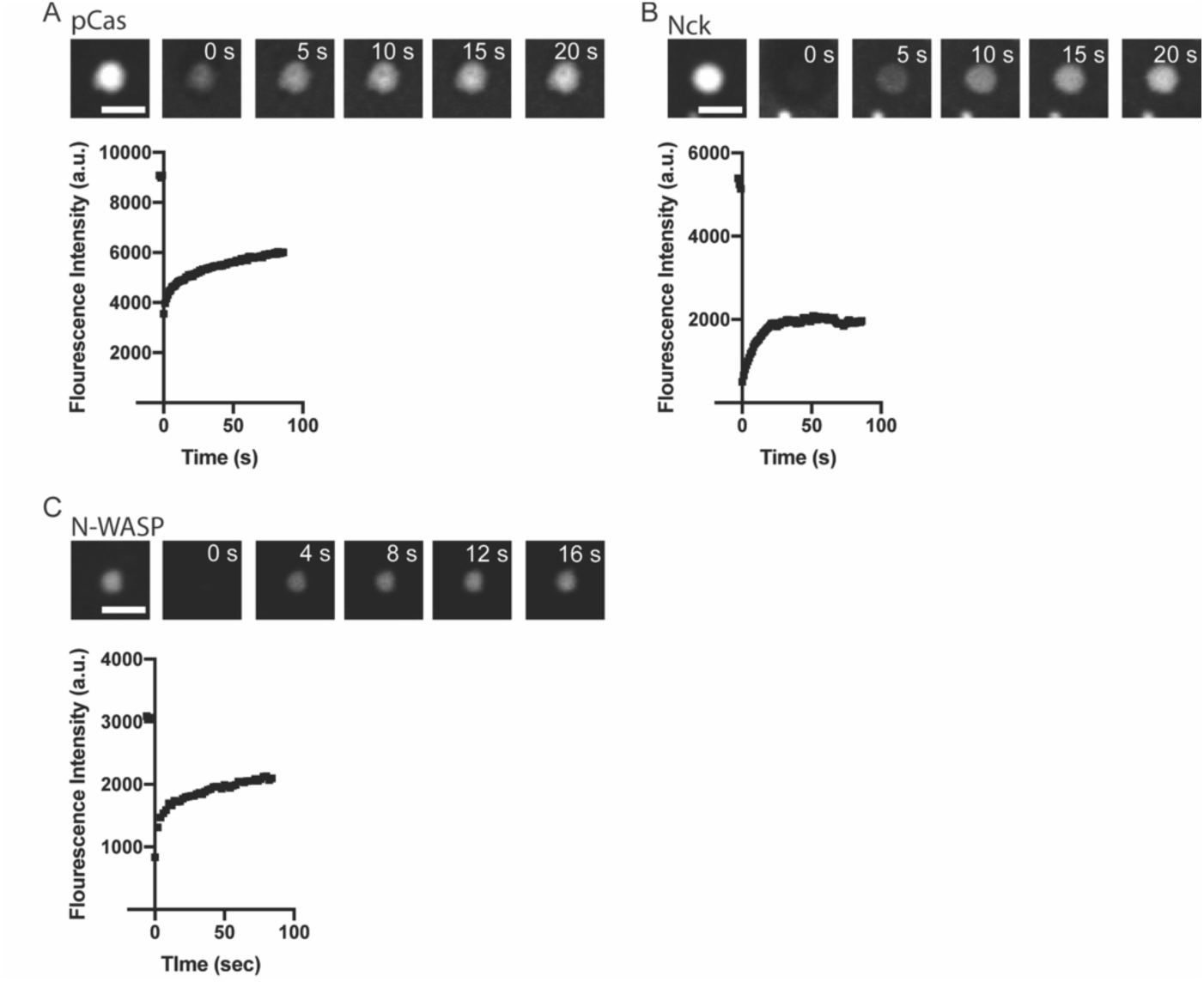
Representative fluorescence recovery after photobleaching (FRAP) data **(A)** TOP: Representative spinning disk fluorescence images of pCas fluorescence recovery after photobleaching (FRAP) experiment. Droplets were formed with 1 µM pCas (5% Alexa-568), 1 µM nck and 1 µM N-WASP, and a single droplet is bleached at t = 0 s. BOTTOM: Quantification of fluorescence intensity of the bleached droplet over time. **(B)** TOP: Representative spinning disk fluorescence images of nck FRAP experiment. Droplets were formed with 1 µM pCas, 1 µM nck (15% Alexa-488), and 1 µM N-WASP, and a single droplet is bleached at t = 0 s. BOTTOM: Quantification of fluorescence intensity of the bleached droplet over time. **(C)** TOP: Representative spinning disk fluorescence images of N-WASP FRAP experiment. Droplets were formed with 1 µM pCas, 1 µM nck, and 1 µM N-WASP (15% Alexa-647), and a single droplet is bleached at t = 0 s. BOTTOM: Quantification of fluorescence intensity of the bleached droplet over time. All scalebars = 5 micron.

**Figure 2 – figure supplement 1.**
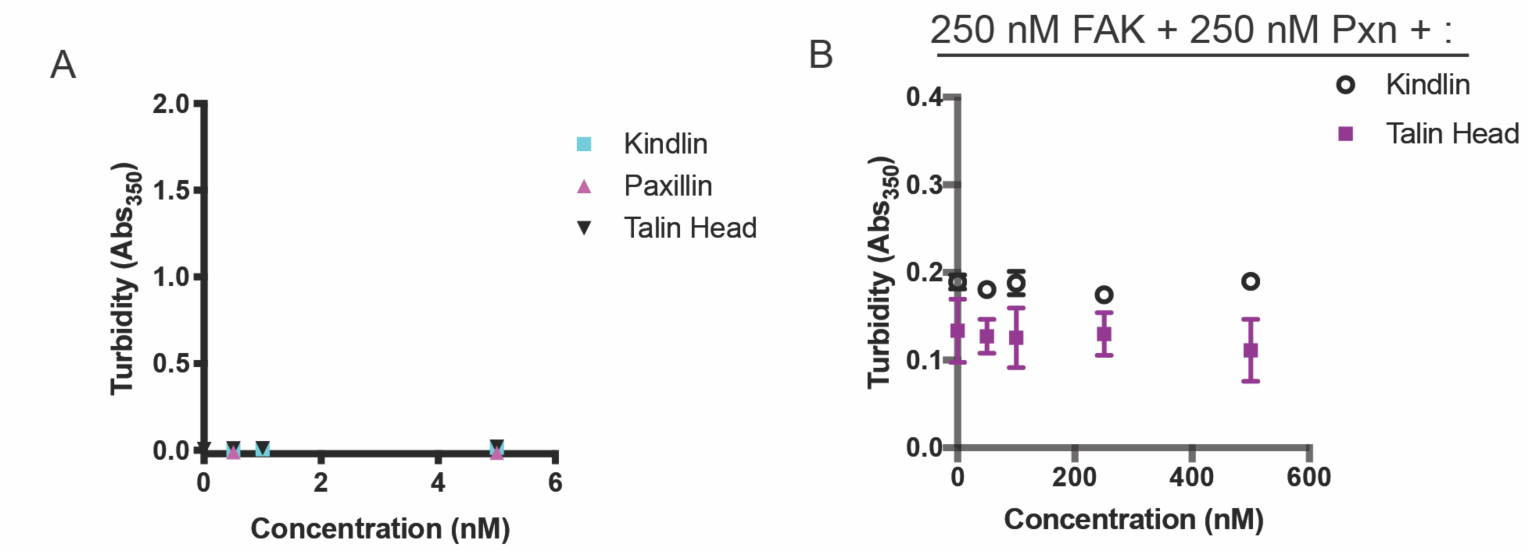
Solution turbidity measurements **(A)** Increasing concentrations of Kindlin, Talin Head or Paxillin were added to solution. **(B)** 250 nM FAK and 250 nM Paxillin were combined with increasing concentrations of Talin Head or Kindlin. In A-B, each point represents the mean ± SEM of three independent measurements.

**Figure 2 – figure supplement 2.**
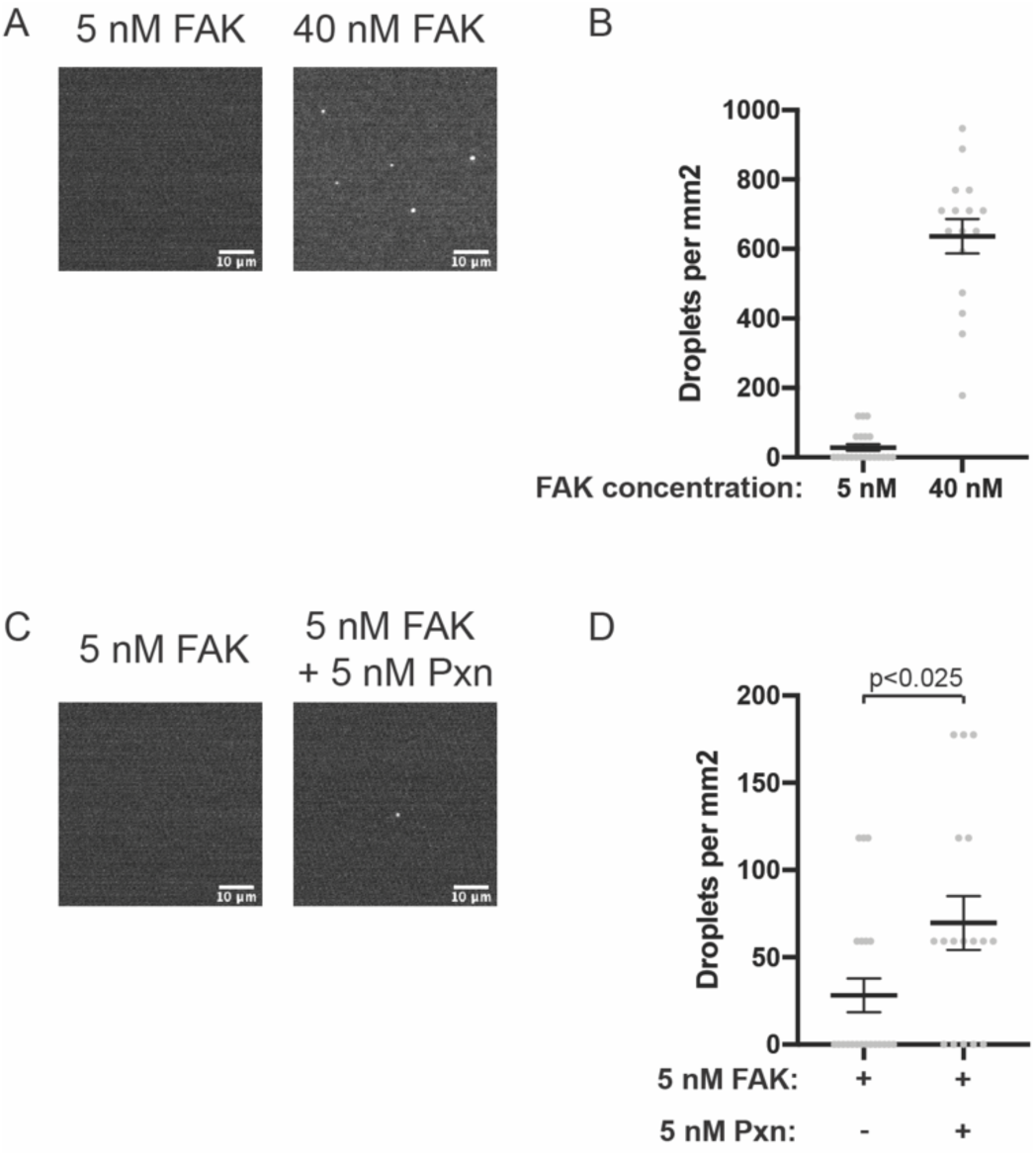
Droplets form with physiological protein concentrations **(A)** Spinning disk fluorescence microscopy images of 5 nM or 40 nM FAK (1% Alexa647). **(B)** Quantification of number of droplets per mm^2^. **(C)** Spinning disk fluorescence microscopy images of 5 nM FAK (1% Alexa647) ± 5 nM paxillin. **(D)** Quantification of number of droplets per mm^2^. All images are displayed with matched contrast settings. Scale bar = 10 µm. In (B) and (D), each grey point represents an individual measurement of a single, randomly selected field of view, and the mean is written and indicated by black line. In (D), significance tested with Student’s t-test.

**Figure 2 – figure supplement 3.**
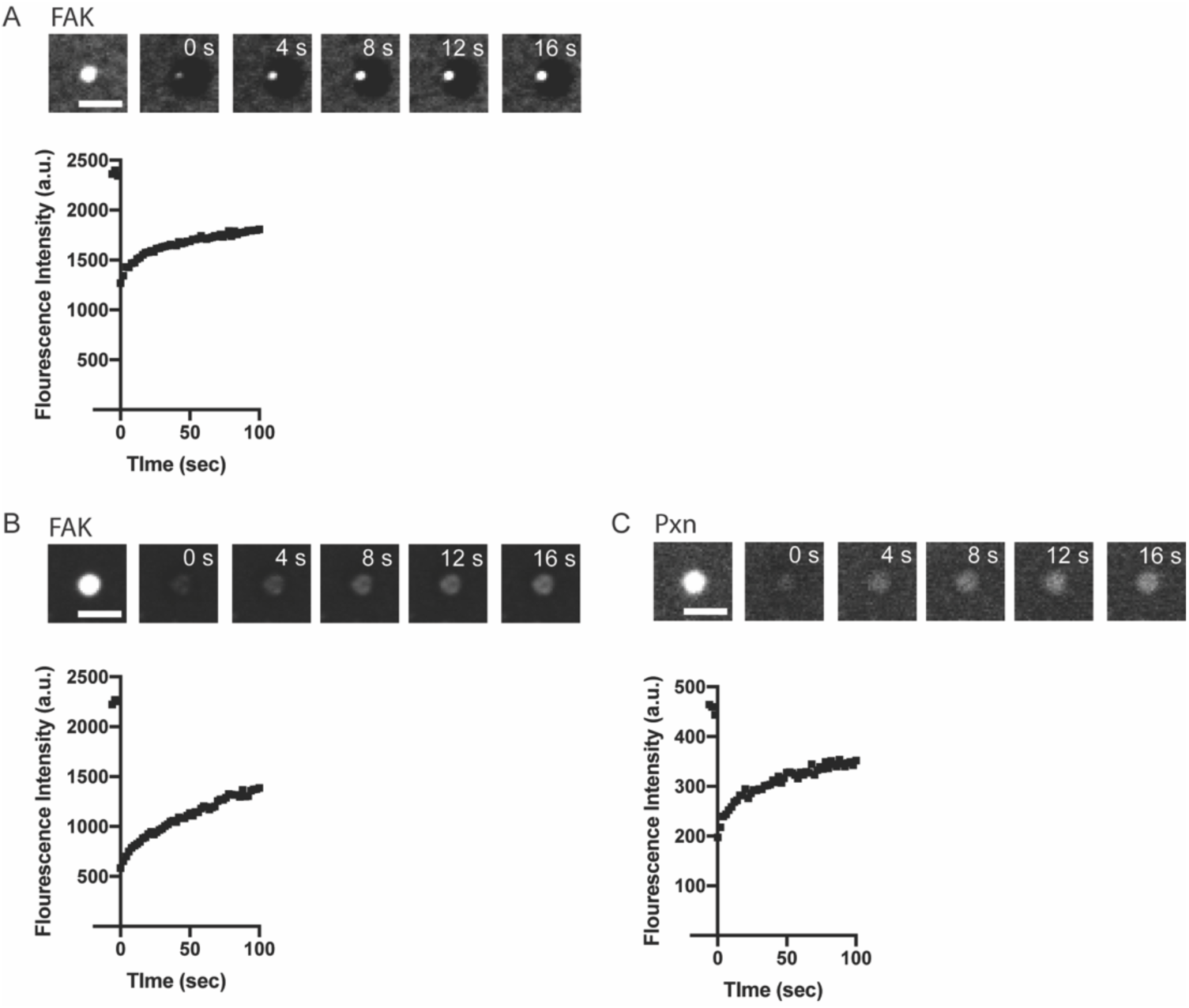
Representative fluorescence recovery after photobleaching (FRAP) data **(A)** TOP: Representative spinning disk fluorescence images of FAK fluorescence recovery after photobleaching (FRAP) experiment. Droplets were formed with 1 µM FAK (1% Alexa-647), and a single droplet is bleached at t = 0 s. BOTTOM: Quantification of fluorescence intensity of the bleached droplet over time. **(B)** TOP: Representative spinning disk fluorescence images of FAK FRAP experiment. Droplets were formed with 1 µM FAK (1% Alexa-647) and 1 µM paxillin, and a single droplet is bleached at t = 0 s. BOTTOM: Quantification of fluorescence intensity of the bleached droplet over time. **(C)** TOP: Representative spinning disk fluorescence images of paxillin FRAP experiment. Droplets were formed with 1 µM FAK and 1 µM paxillin (5% Alexa-561), and a single droplet is bleached at t = 0 s. BOTTOM: Quantification of fluorescence intensity of the bleached droplet over time. All scalebars = 5 micron.

**Figure 3 – figure supplement 1.**
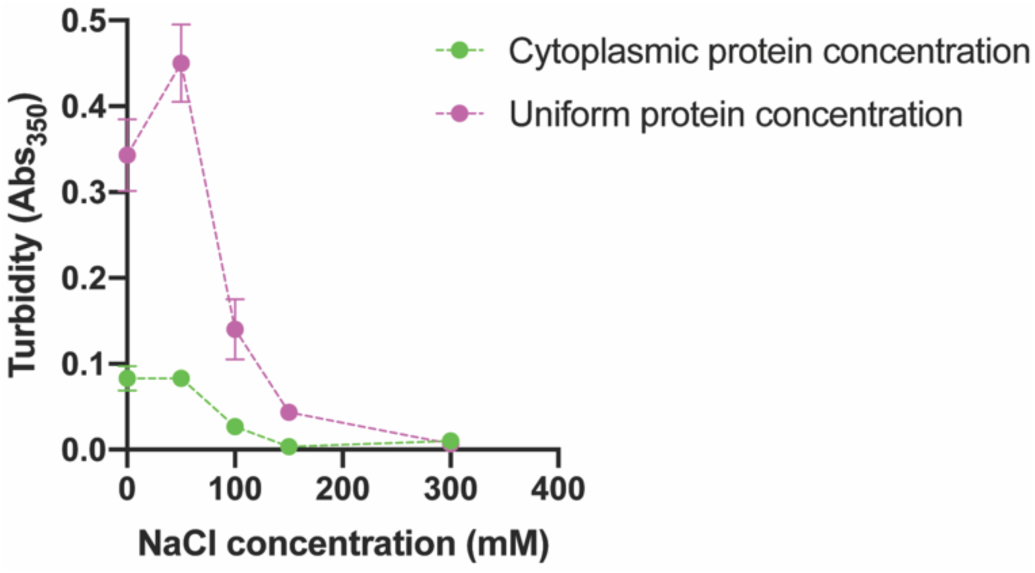
NaCl reduces solution turbidity Solution was prepared with cytoplasmic protein concentration (green; 70 nM pCas, 180 nM nck, 140 nM N-WASP, 60 nM paxillin and 40 nM FAK) or uniform protein concentration (magenta; 250 nM each of pCas, nck, N-WASP, FAK and paxillin) in buffer containing 50 mM Hepes pH 7.3, 0.1% BSA and increasing concentrations (0-300 mM) NaCl. Each point represents the mean ± SEM of three independent measurements

**Figure 3 – figure supplement 2.**
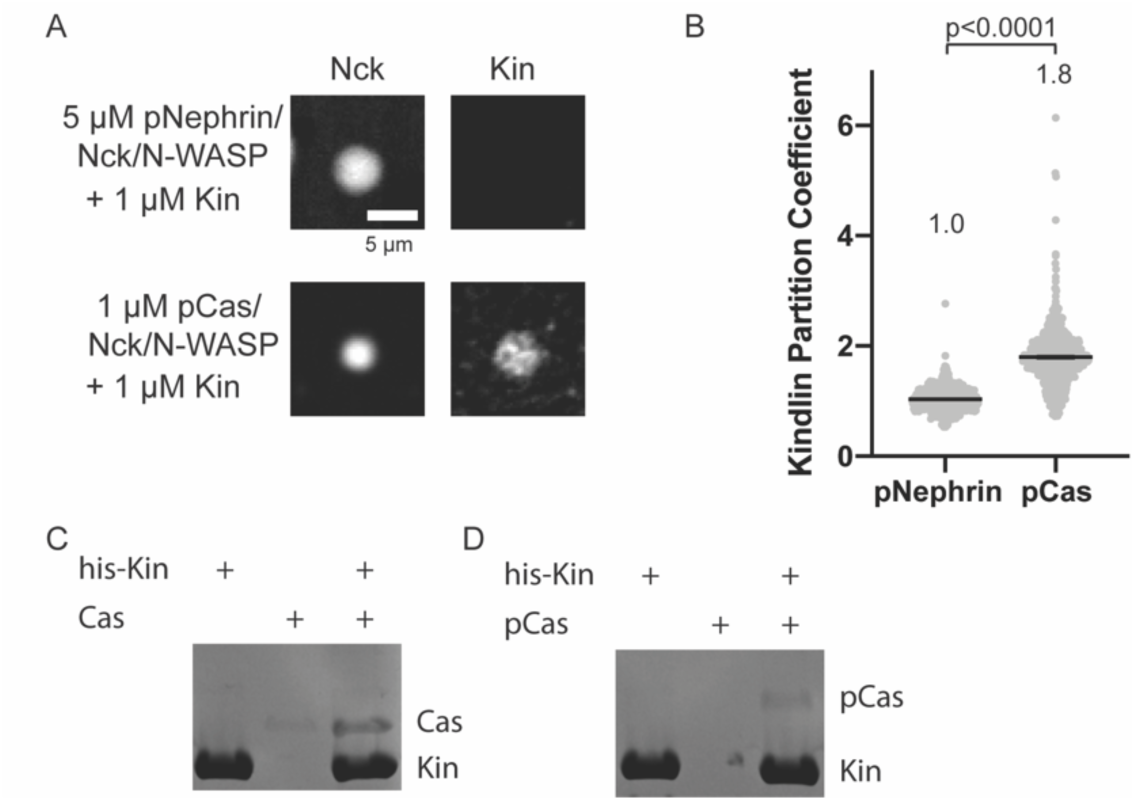
Kindlin interacts with p130Cas **(A)** Spinning disk fluorescence microscopy images of droplets. TOP: 5 µM each of phosphorylated Nephrin, Nck (15% Alexa647), and N-WASP and 1 µM Kindlin (10% Alexa546). BOTTOM: 1 µM each of phosphorylated Cas, Nck (15% Alexa647), N-WASP, and Kindlin (10% Alexa546). Kindlin images displayed with identical contrast settings. Scalebar = 5 µm. **(B)** Quantification of Kindlin partitioning in droplets. Each grey point represents an individual measurement, and the mean is written and indicated by black line. Each condition contains at least 500 measurements. Significance tested with Student’s t-test. **(C-D)** His pull-down samples analyzed on SDS-PAGE gel with Coomassie blue staining. Original raw files of gels in Figure3-figure supplement2-source data.

**Figure 4 – figure supplement 1.**
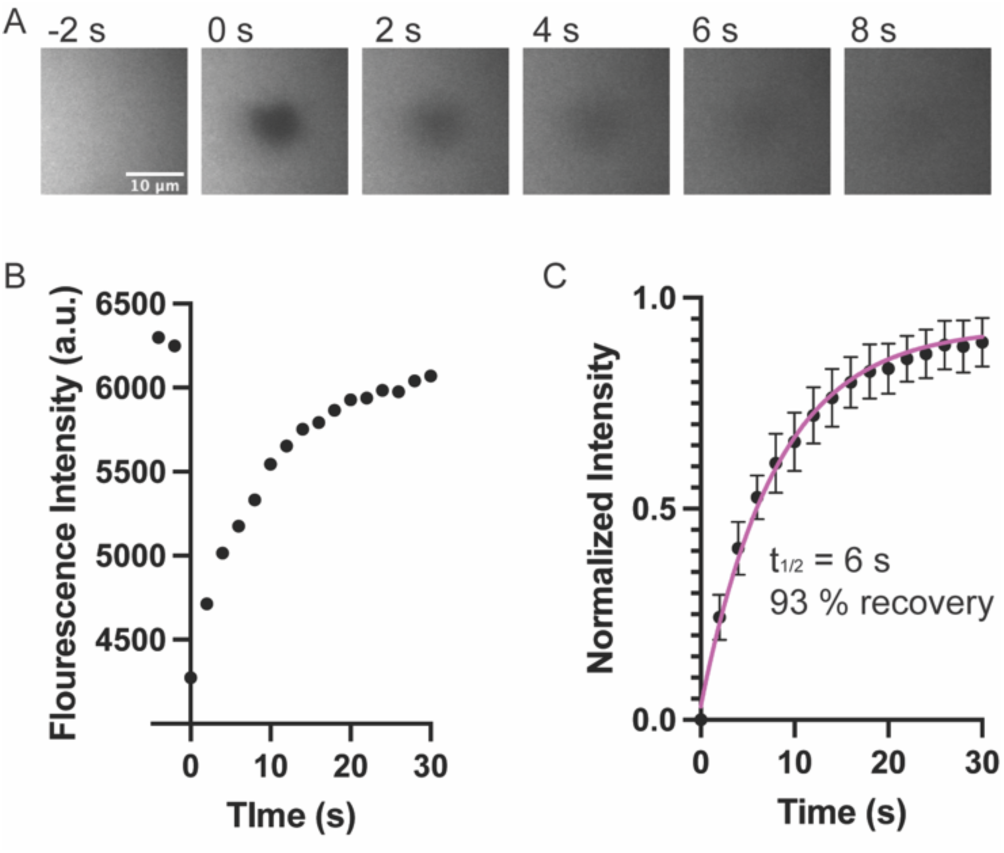
Analysis of phospholipid bilayer fluidity (A) Representative images of fluorescence phospholipid bilayers imaged with Total Internal Reflection Fluorescence (TIRF) microscopy during photobleaching analysis. Bilayers containing 97% POPC. 2% Ni-NTA, 1% PC-NBD, and 0.1% PEG-PE were imaged with 488 nm TIRF illumination to directly visualize PC-NBD lipids. A 5-micron region was bleached at t=0 s. Scalebar = 10 micron. (B) Quantification of NDB fluorescence intensity of the bleached region from A. (C) Fluorescence Recovery After Photobleaching (FRAP) measurements of bilayer. Each point represents the mean ± SEM of at 7 independent measurements. The t_1/2_ was calculated from a single exponential fit; fit overlayed on graph (magenta line).

**Figure 6 – figure supplement 1.**
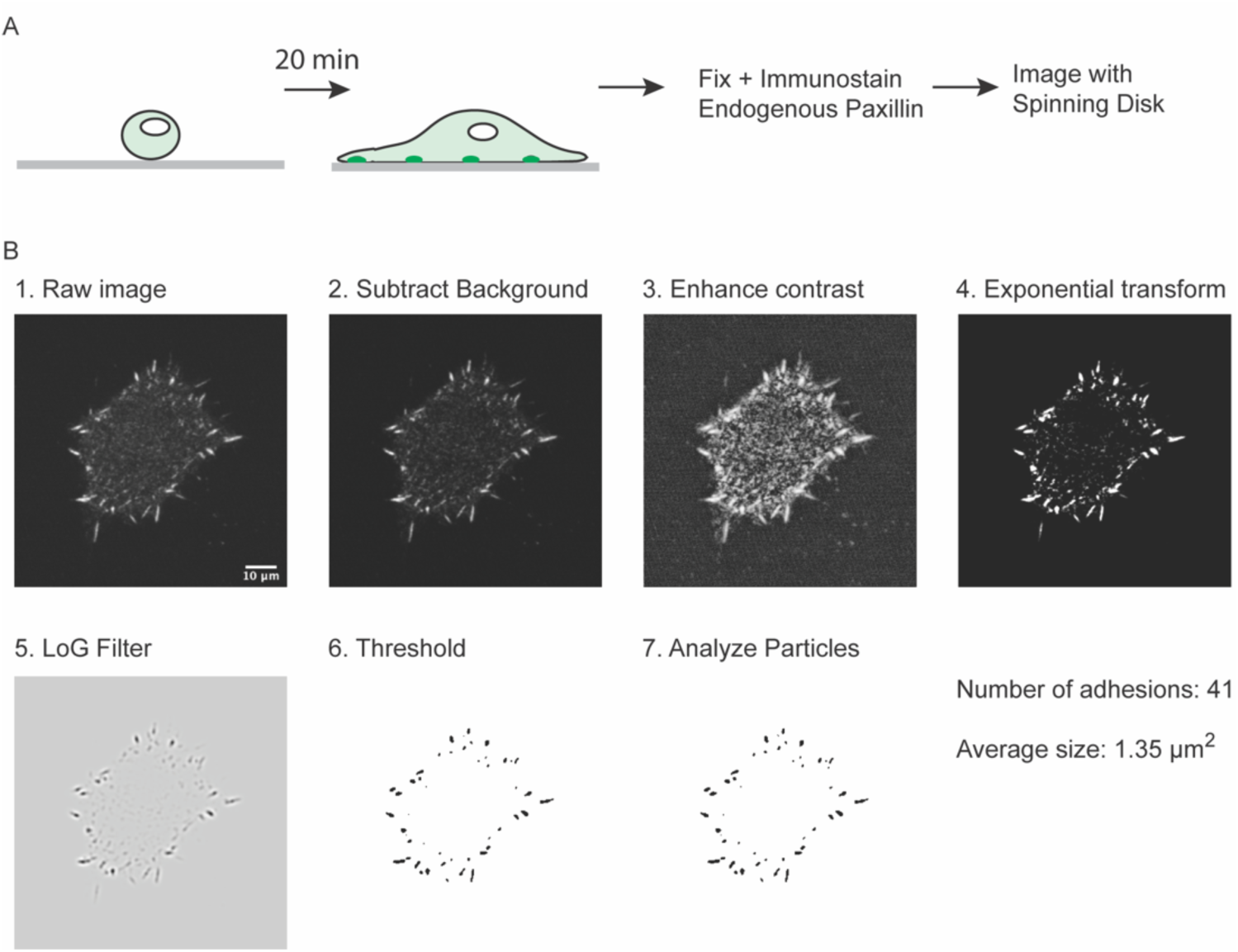
Measuring number of integrin adhesions during cell spreading **(A)** Experimental workflow **(B)** Example of image analysis workflow. Scalebar = 10 µm.

**Figure 6 – figure supplement 2.**
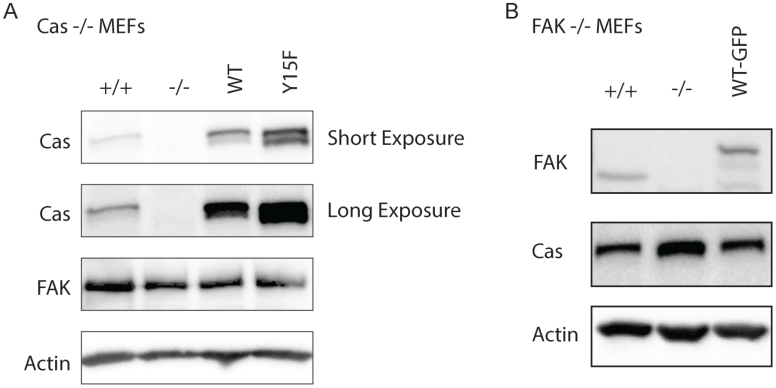
Westernblot analysis of MEF cell lines Cell lysates were blotted with antibodies to p130Cas (Cas), FAK, and actin. **(A)** Cas MEF cell lines were assessed (Cas +/+, Cas -/-, Cas-/- + WT Cas, Cas-/- + Y15F Cas). **(B)** FAK MEF cell lines were assessed (FAK+/+, FAK-/-, FAK-/- +WT-FAK-GFP). Original raw files of blots in Figure6-figure supplement2-source data.

**Figure 6 – figure supplement 3.**
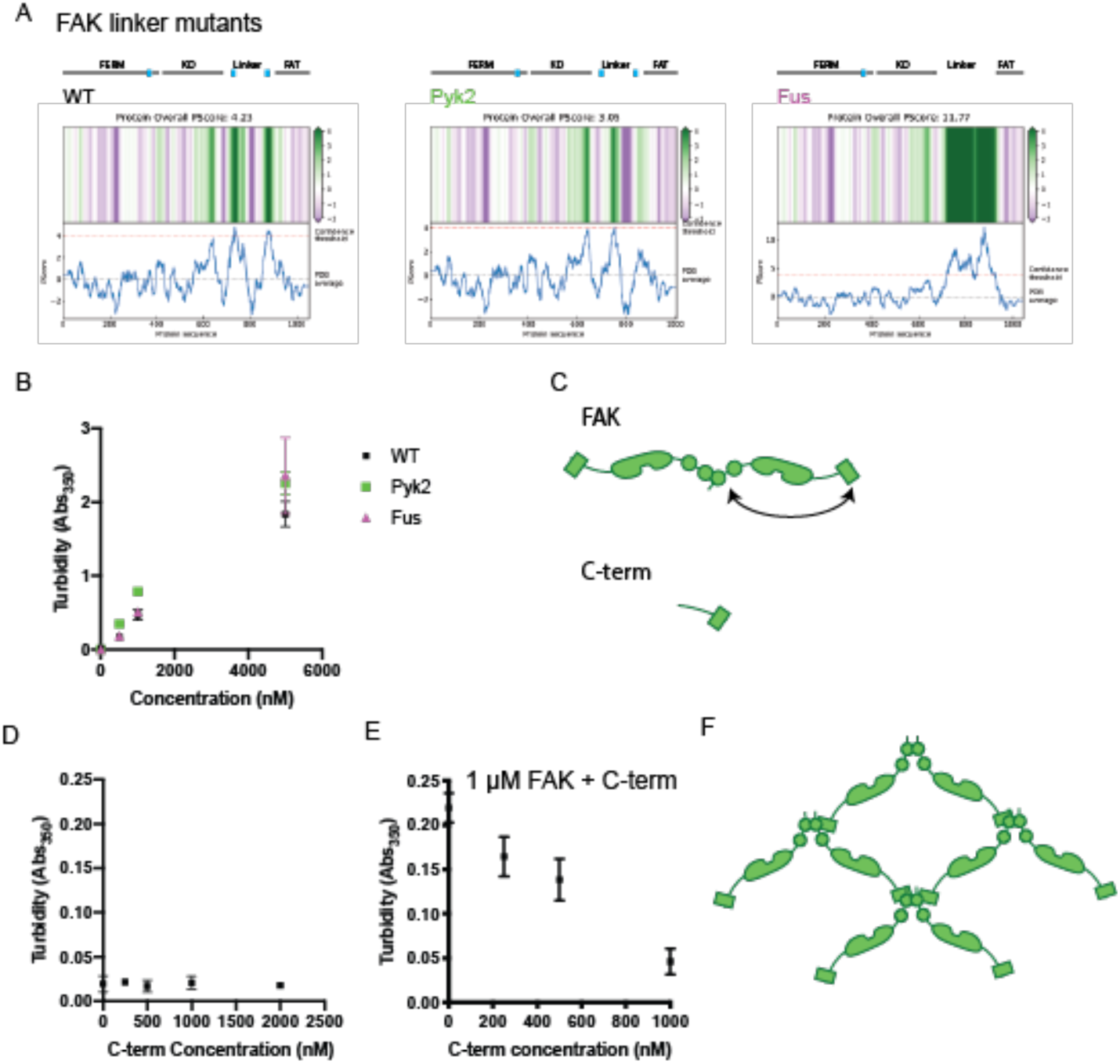
*In vitro* analysis of FAK phase separation **(A)** Computational analysis of FAK linker mutant phase separation propensity. **(B)** Solution turbidity measurements with increasing concentrations of FAK linker mutants. WT FAK data is duplicated from Fig. 2B. **(C)** Cartoon schematic of FAK C-terminal truncation (”C-term”). Full-length FAK with FERM domain and FAT domain annotated is shown for comparison. **(D)** Solution turbidity measurements with increasing concentrations of FAK C-terminal truncation. **(E)** Solution turbidity measurements with 1 µM FAK combined with increasing concentrations of FAK C-terminal truncation. In (B), (D), and (E) each point represents mean ± SEM. **(F)** Illustration demonstrating how FERM:FERM + FERM:FAT interactions could promote higher-order oligomerization.

**Figure 6 – figure supplement 4.**
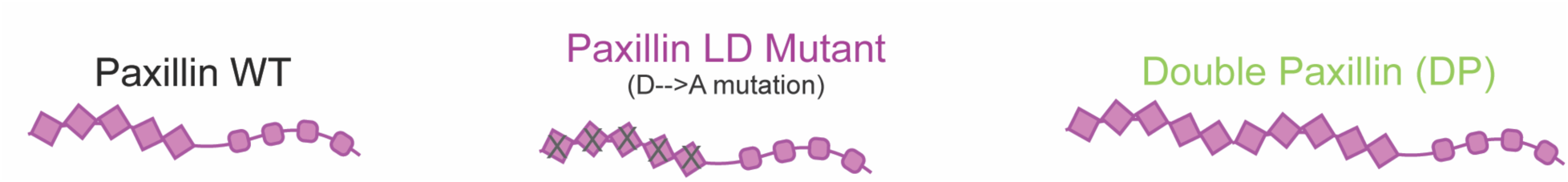
Paxillin valence variants **(A)** Cartoon schematic of engineered paxillin valence variants. Paxillin LD mutant has reduced FAK binding, while double paxillin (DP) has increased number of FAK binding sites.

**Figure 7 – figure supplement 1.**
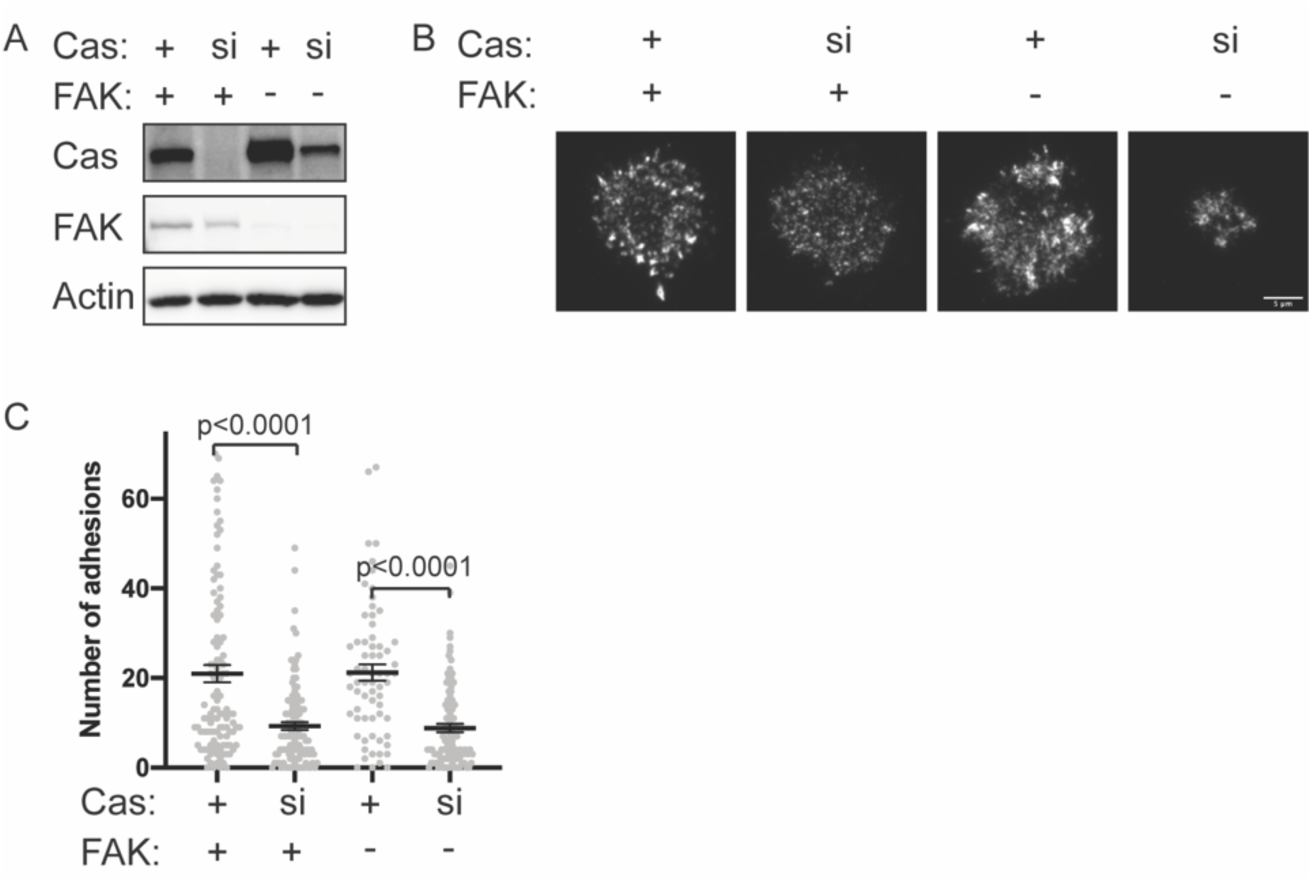
Number of adhesions in FAK -/- cells after 5 minutes spreading. **(A)** Western blot analysis of Cas knockdown. FAK^+/+^ or FAK ^-/-^ MEFs were treated with nontargeting (+) or Cas (si) siRNA for 48 hours and lysates were blotted with Cas, FAK or actin antibodies. Original raw files of blots in Figure7-figure supplement 1-source data. **(B)** Total Internal Reflection Fluorescence (TIRF) microscopy images of MEFs fixed after 5 minutes of spreading with immunostaining for endogenous paxillin. Scalebar = 5 micron. **(C)** Quantification of number of adhesions. Each grey point represents a measurement from one cell, and the mean ± SEM mean is indicated by black lines. Data from at least 45 cells from 2 or more independent experiments. Significance tested by one-way ANOVA followed by a Tukey multiple comparison test.

**Figure 7 – figure supplement 2.**
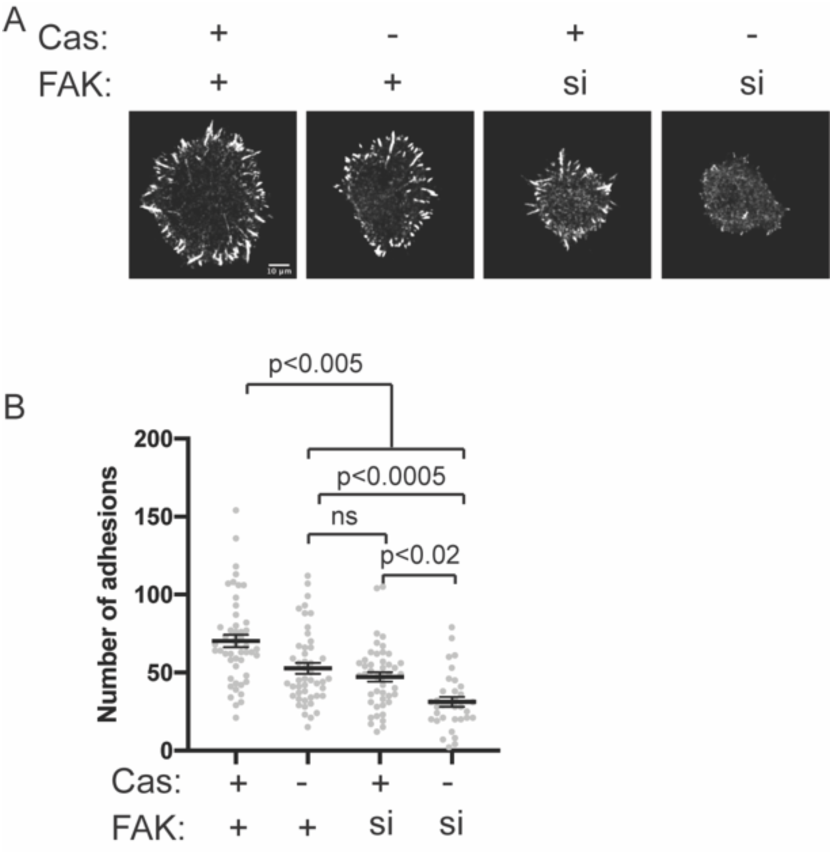
Number of adhesions in Cas -/- cells after 20 minutes spreading **(A)** Spinning disk confocal microscopy images of MEFs fixed after 20 minutes of spreading with immunostaining for endogenous paxillin. Scalebar = 10 micron. **(B)** Quantification of number of adhesions. Each grey point represents a measurement from one cell, and the mean ± SEM mean is indicated by black lines. Data from at least 30 cells from 2 or more independent experiments. Significance tested by one-way ANOVA followed by a Tukey multiple comparison test.

**Table S1.**
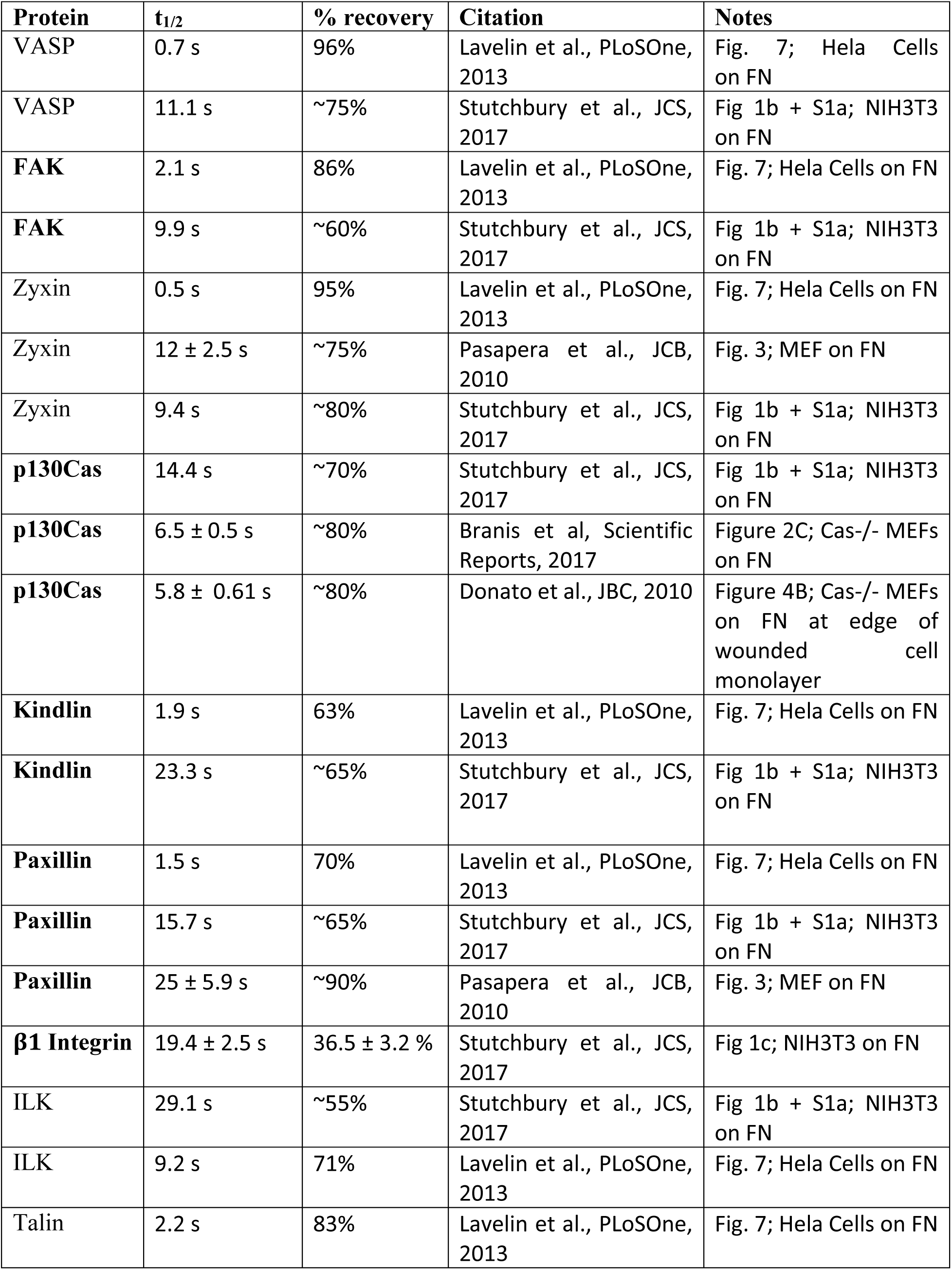
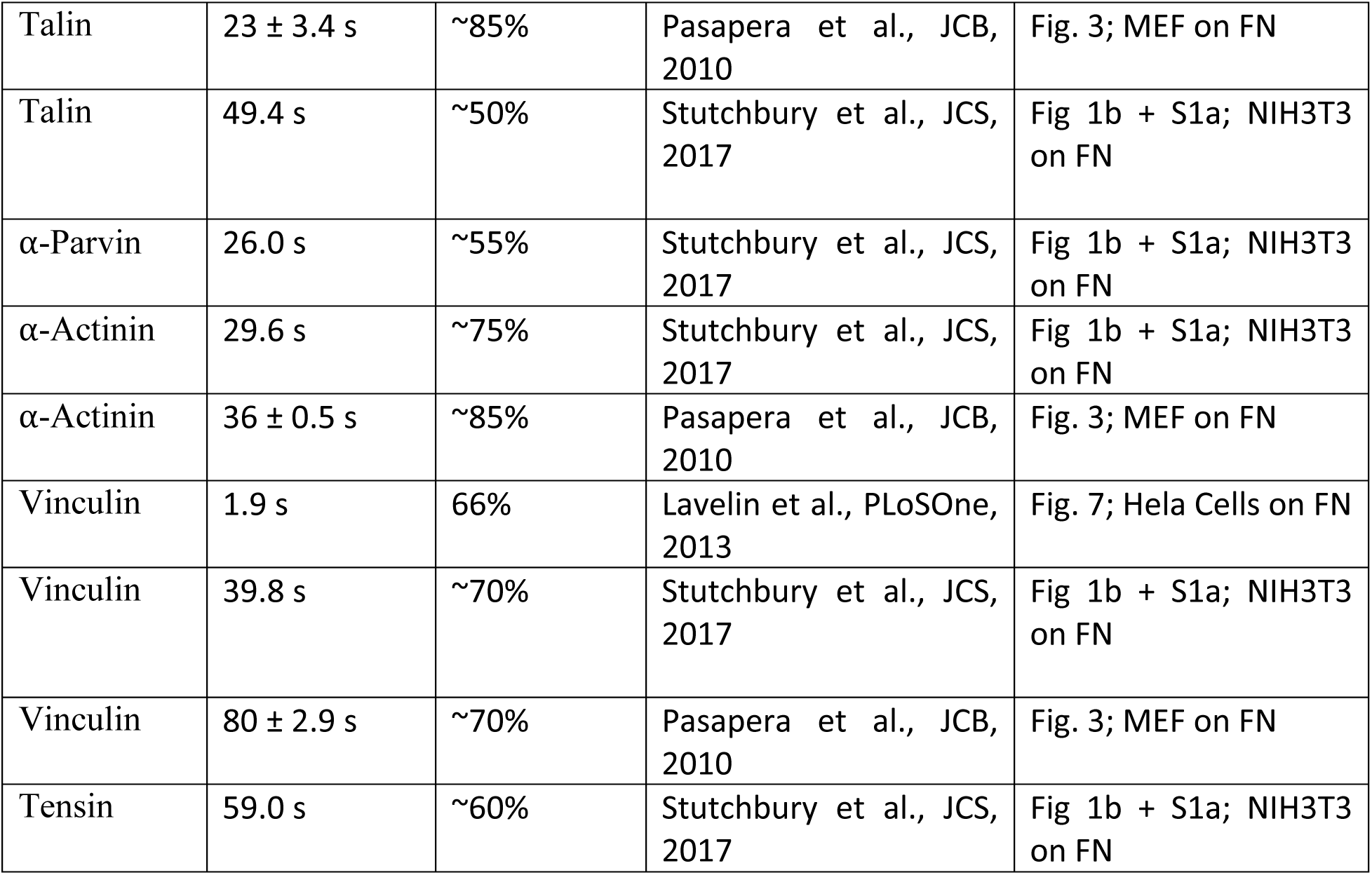
Curation of published Fluorescence Recovery After Photobleaching (FRAP) measurements of integrin adhesion components. Proteins used in this study are in **bold**. ∼ indicates the percent recovery was estimated from graph.

| **Protein** | **t<sub>1/2</sub>** | **% recovery** | **Citation** | **Notes** |
| --- | --- | --- | --- | --- |
| VASP | 0.7 s | 96% | Lavelin et al., PLoSOne, 2013 | Fig. 7; Hela Cells on FN |
| VASP | 11.1 s | ~75% | Stutchbury et al., JCS, 2017 | Fig 1b + S1a; NIH3T3 on FN |
| **FAK** | 2.1 s | 86% | Lavelin et al., PLoSOne, 2013 | Fig. 7; Hela Cells on FN |
| **FAK** | 9.9 s | ~60% | Stutchbury et al., JCS, 2017 | Fig 1b + S1a; NIH3T3 on FN |
| Zyxin | 0.5 s | 95% | Lavelin et al., PLoSOne, 2013 | Fig. 7; Hela Cells on FN |
| Zyxin | 12 ± 2.5 s | ~75% | Pasapera et al., JCB, 2010 | Fig. 3; MEF on FN |
| Zyxin | 9.4 s | ~80% | Stutchbury et al., JCS, 2017 | Fig 1b + S1a; NIH3T3 on FN |
| **p130Cas** | 14.4 s | ~70% | Stutchbury et al., JCS, 2017 | Fig 1b + S1a; NIH3T3 on FN |
| **p130Cas** | 6.5 ± 0.5 s | ~80% | Branis et al, Scientific Reports, 2017 | Figure 2C; Cas-/- MEFs on FN |
| **p130Cas** | 5.8 ± 0.61 s | ~80% | Donato et al., JBC, 2010 | Figure 4B; Cas-/- MEFs on FN at edge of wounded cell monolayer |
| **Kindlin** | 1.9 s | 63% | Lavelin et al., PLoSOne, 2013 | Fig. 7; Hela Cells on FN |
| **Kindlin** | 23.3 s | ~65% | Stutchbury et al., JCS, 2017 | Fig 1b + S1a; NIH3T3 on FN |
| **Paxillin** | 1.5 s | 70% | Lavelin et al., PLoSOne, 2013 | Fig. 7; Hela Cells on FN |
| **Paxillin** | 15.7 s | ~65% | Stutchbury et al., JCS, 2017 | Fig 1b + S1a; NIH3T3 on FN |
| **Paxillin** | 25 ± 5.9 s | ~90% | Pasapera et al., JCB, 2010 | Fig. 3; MEF on FN |
| **β1 Integrin** | 19.4 ± 2.5 s | 36.5 ± 3.2 % | Stutchbury et al., JCS, 2017 | Fig 1c; NIH3T3 on FN |
| ILK | 29.1 s | ~55% | Stutchbury et al., JCS, 2017 | Fig 1b + S1a; NIH3T3 on FN |
| ILK | 9.2 s | 71% | Lavelin et al., PLoSOne, 2013 | Fig. 7; Hela Cells on FN |
| Talin | 2.2 s | 83% | Lavelin et al., PLoSOne, 2013 | Fig. 7; Hela Cells on FN |
| Talin | 23 ± 3.4 s | ~85% | Pasapera et al., JCB, 2010 | Fig. 3; MEF on FN |
| Talin | 49.4 s | ~50% | Stutchbury et al., JCS, 2017 | Fig 1b + S1a; NIH3T3 on FN |
| α-Parvin | 26.0 s | ~55% | Stutchbury et al., JCS, 2017 | Fig 1b + S1a; NIH3T3 on FN |
| α-Actinin | 29.6 s | ~75% | Stutchbury et al., JCS, 2017 | Fig 1b + S1a; NIH3T3 on FN |
| α-Actinin | 36 ± 0.5 s | ~85% | Pasapera et al., JCB, 2010 | Fig. 3; MEF on FN |
| Vinculin | 1.9 s | 66% | Lavelin et al., PLoSOne, 2013 | Fig. 7; Hela Cells on FN |
| Vinculin | 39.8 s | ~70% | Stutchbury et al., JCS, 2017 | Fig 1b + S1a; NIH3T3 on FN |
| Vinculin | 80 ± 2.9 s | ~70% | Pasapera et al., JCB, 2010 | Fig. 3; MEF on FN |
| Tensin | 59.0 s | ~60% | Stutchbury et al., JCS, 2017 | Fig 1b + S1a; NIH3T3 on FN |

**Table S2.**
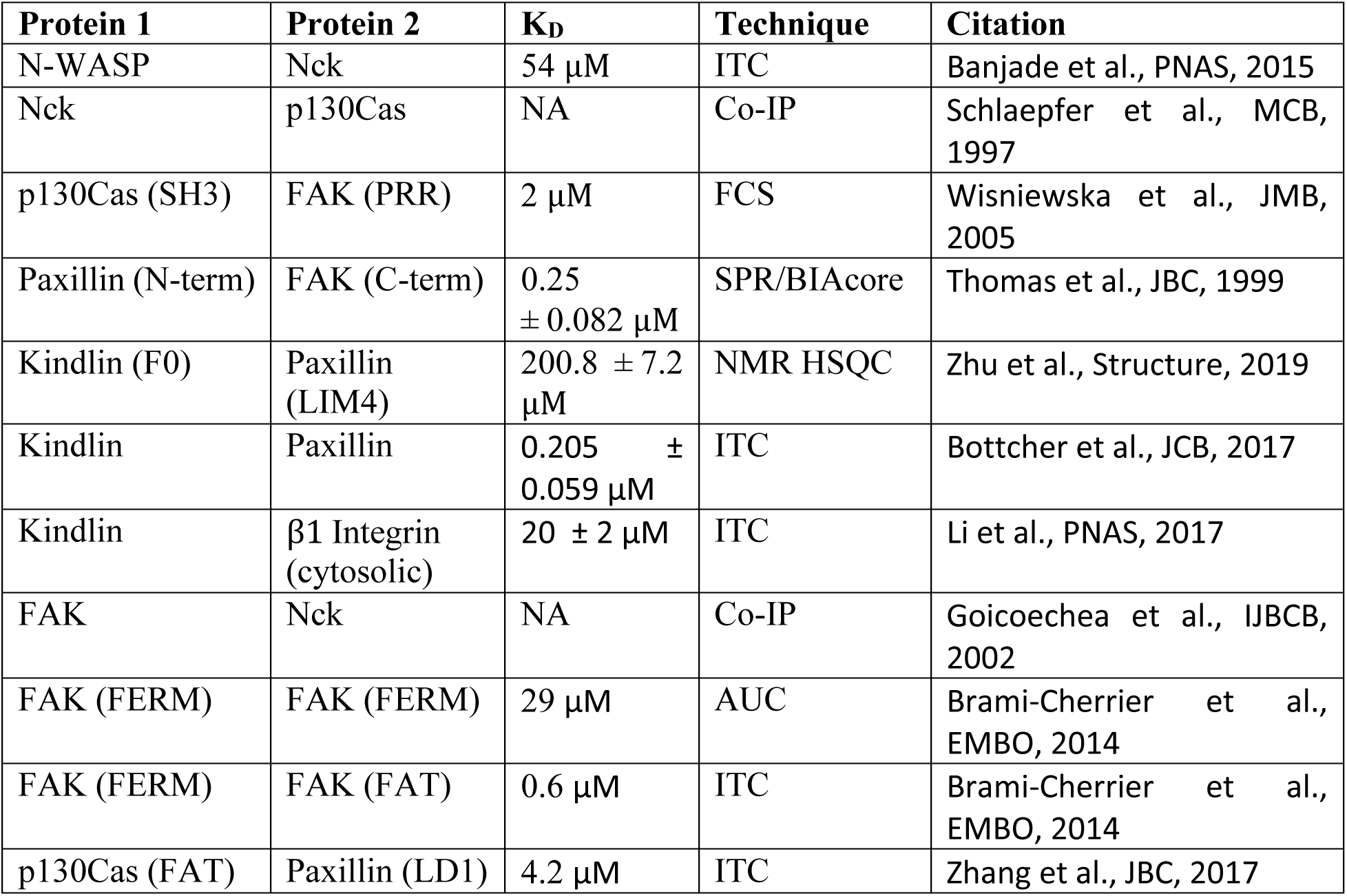
Curation of published interactions between proteins used in this study Unless the specific domain is indicated in parentheses, studies used full-length protein. Technique abbreviations: ITC: Isothermal Titration Calorimetry; FCS: Co-IP: Co-immunoprecipitation; Fluorescence correlation spectroscopy; SPR: Surface Plasmon Resonance; NMR HSQC: Nuclear Magnetic Resonance Heteronuclear Single Quantum Coherence; AUC: Analytical ultracentrifugation.

**Table S3.** Curation of published cellular concentrations of proteins used in this study.

| **Protein** | **Concentration** | **Cell Type** | **Citation** |
| --- | --- | --- | --- |
| WASP | 9000 nM | human peripheral neutrophil | Higgs and Pollard, JCB, 2000 |
| WASP | 2000-8000 nM | T-cell (primed primary CD4+ T cells from 5C.C7 TCR transgenic mice) | Roybal et al., Sci Signaling, 2016 |
| N-WASP | 138 nM | HeLa | Hein et al, Cell, 2015 |
| Nck1 | 180 nM | HeLa | Hein et al, Cell, 2015 |
| p130Cas | 72 nM | HeLa | Hein et al, Cell, 2015 |
| FAK | 40 nM | HeLa | Hein et al, Cell, 2015 |
| FAK | 5-6 nM | NIH3T3 | Brami-Cherrier, EMBO, 2014 |
| Paxillin | 61 nM | HeLa | Hein et al, Cell, 2015 |
| Kindlin | 63 nM | HeLa | Hein et al, Cell, 2015 |
| Integrin β1 | 188 nM | HeLa | Hein et al, Cell, 2015 |

**Table S4.** Floursecent Recovery After Photobleaching (FRAP) curve exponential fit of data. Biexponential and single exponential fits were statistically compared with an extra sum-of-squares F Test to determine the best fit. Values of best fit are shown in table. nd = not determined. For several molecules, the percent recovery could not be accurately fit from these data, as the recovery did not sufficiently plateau within the 100 second experiment.

| **Molecule (bi-exponential)** | **Fast $t_{1/2}$** | **Slow $t_{1/2}$** | **Percent Fast** | **Recovery** |
| --- | --- | --- | --- | --- |
| pCas | 2 s | 39 s | 35% | nd |
| N-WASP | 1 s | 37 s | 30% | 99% |
| FAK (alone) | 6 s | 56 s | 36% | 62% |
| FAK (+Paxillin) | 10 s | 98 s | 21% | nd |
| Paxillin | 7 s | 78 s | 34% | nd |
| **Molecule (single exponential)** | **$t_{1/2}$** | **Recovery** |  |  |
| Nck | 8 s | 39% |  |  |

**Table S5.**
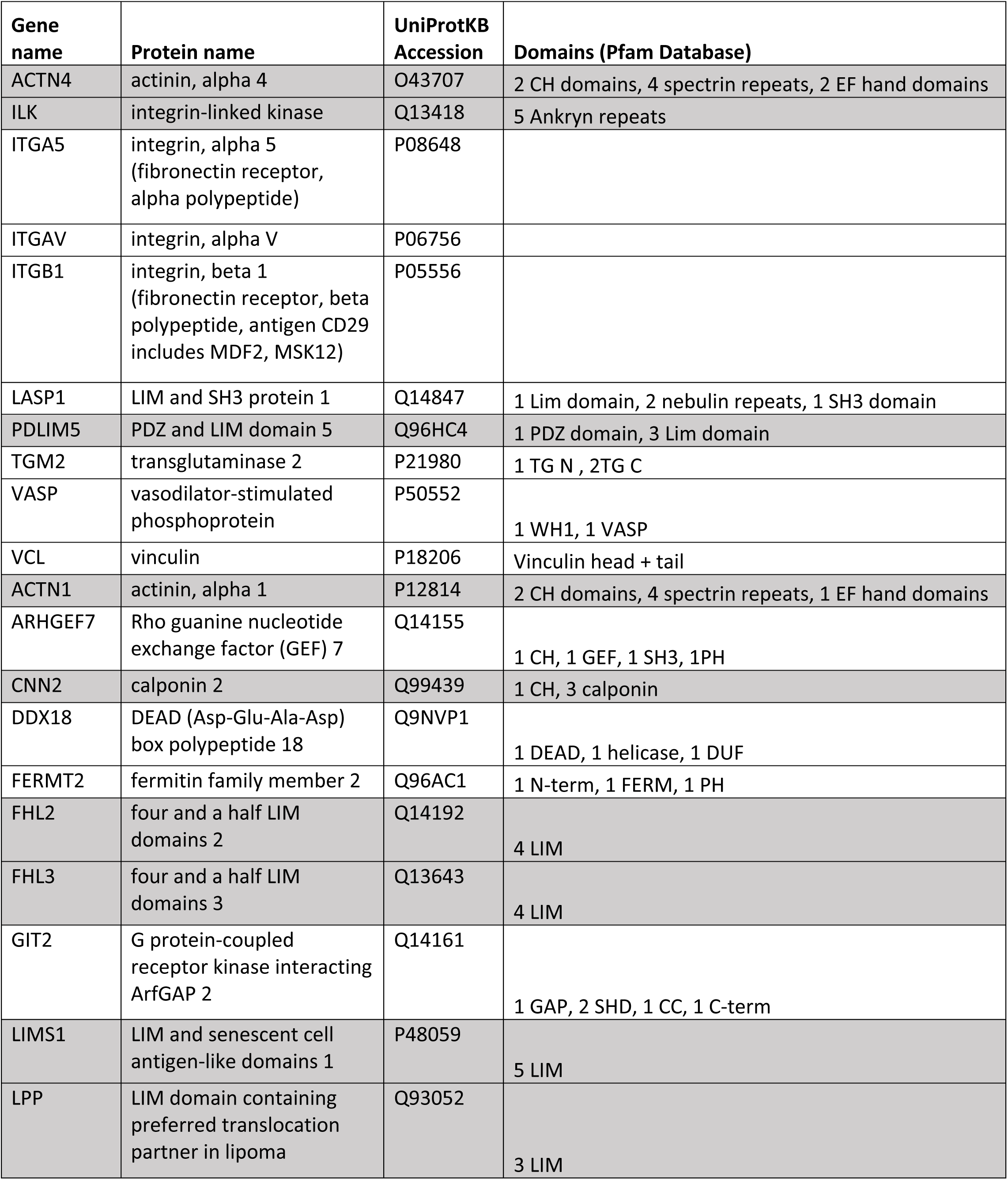
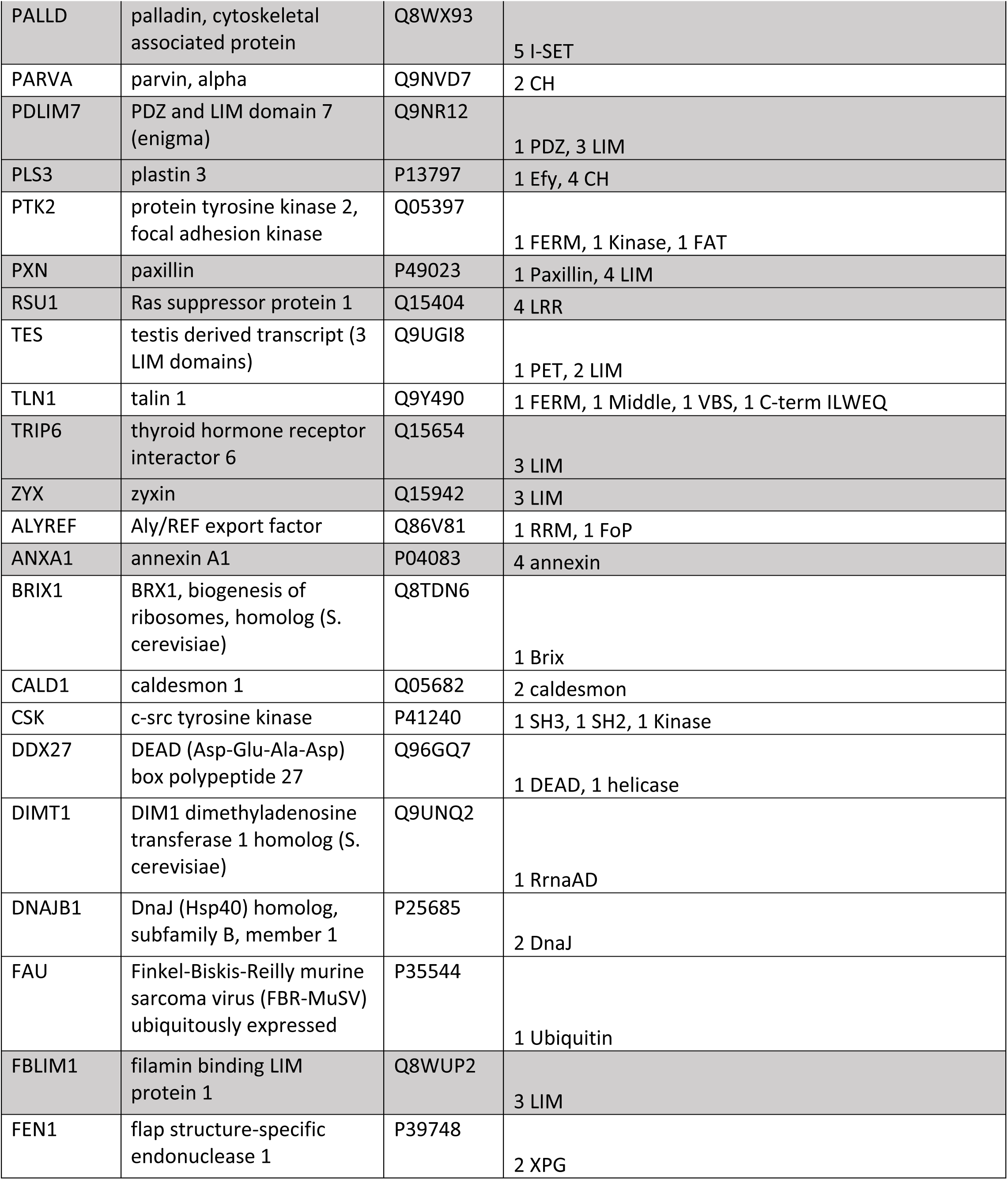
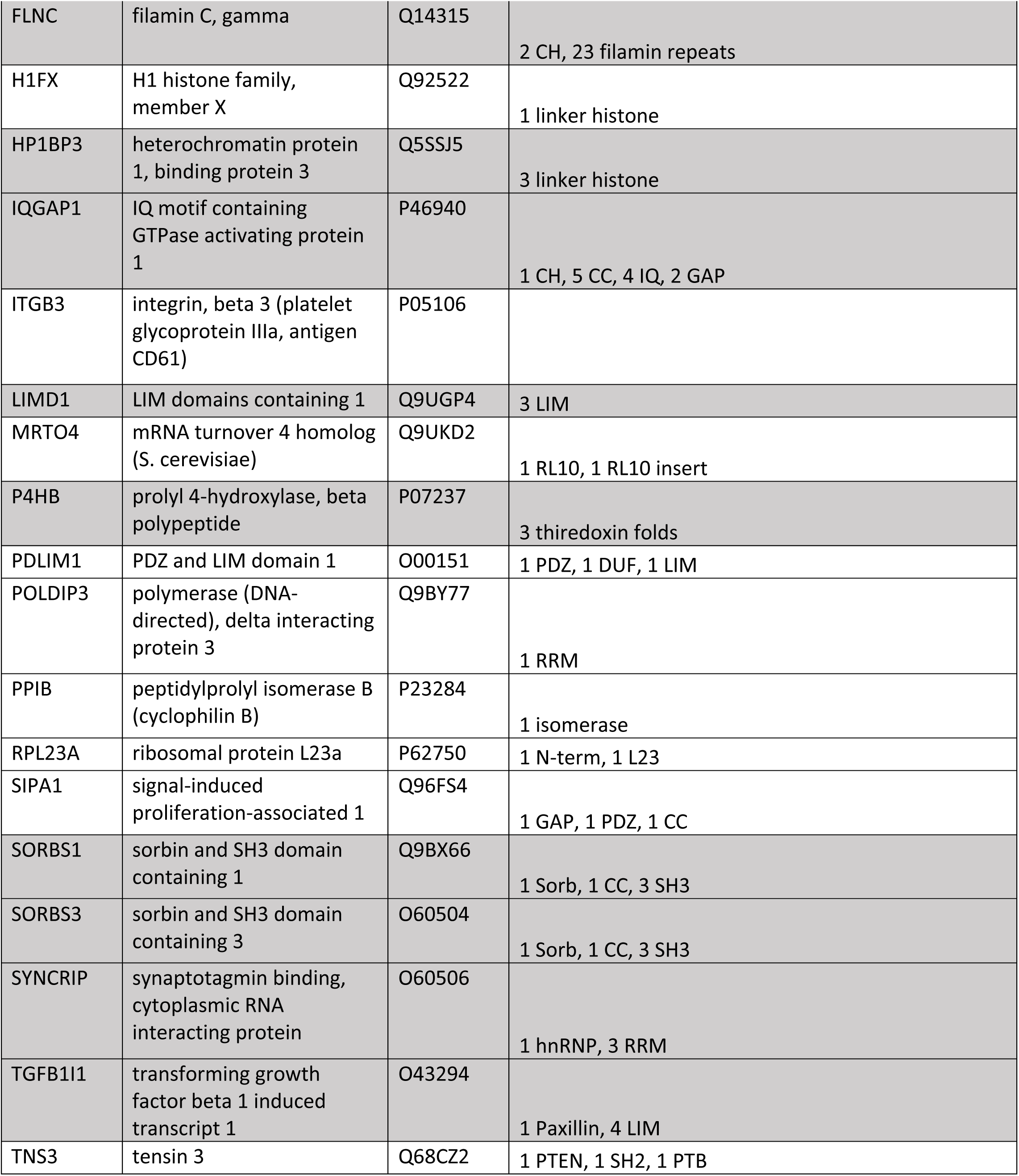
Protein domains of consensus adhesome components (Adhesome components identified in Horton et al., 2015; domains identified using http://pfam.xfam.org). Proteins with multivalent domains shaded grey).

**Table S6.**
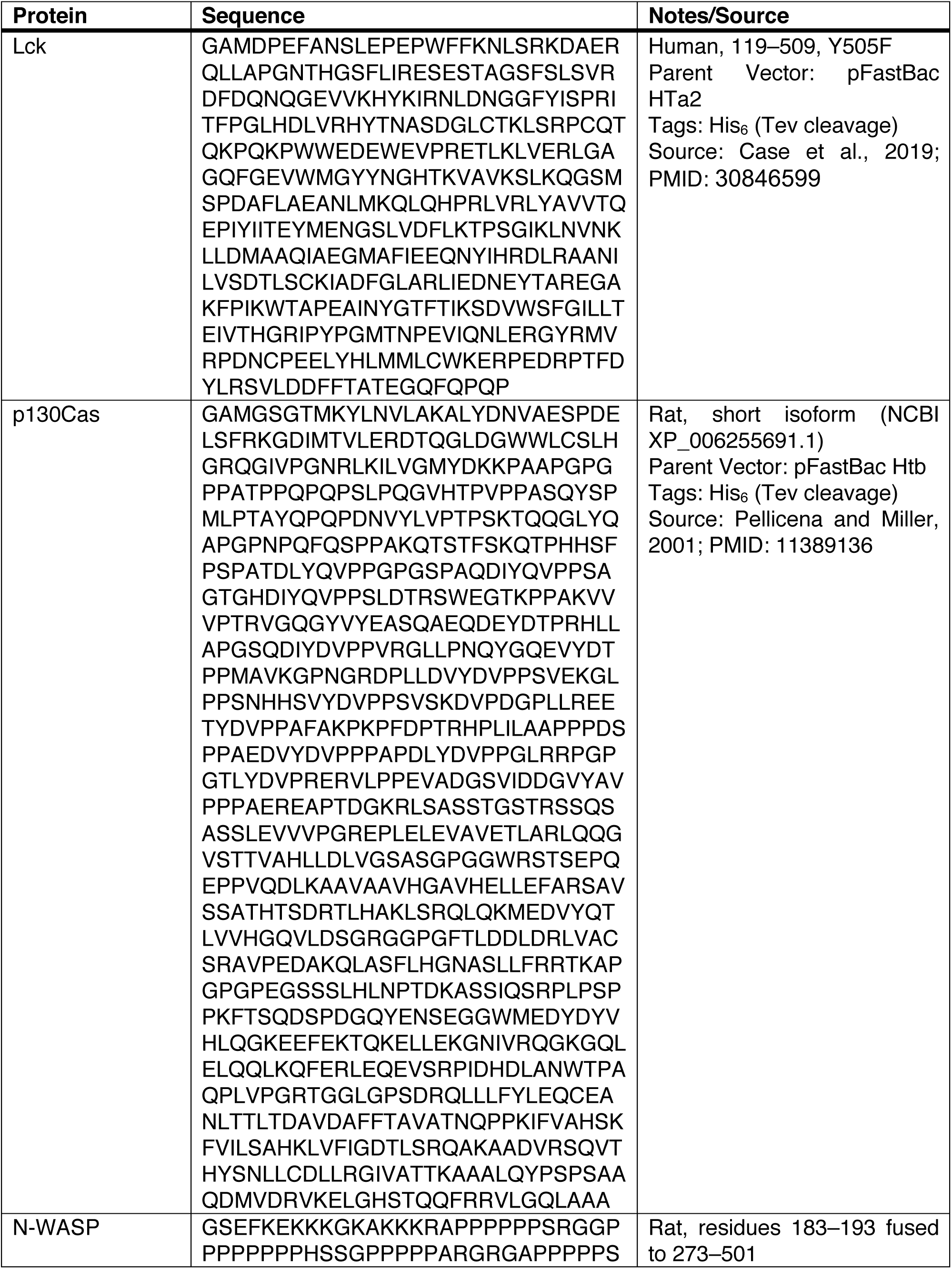
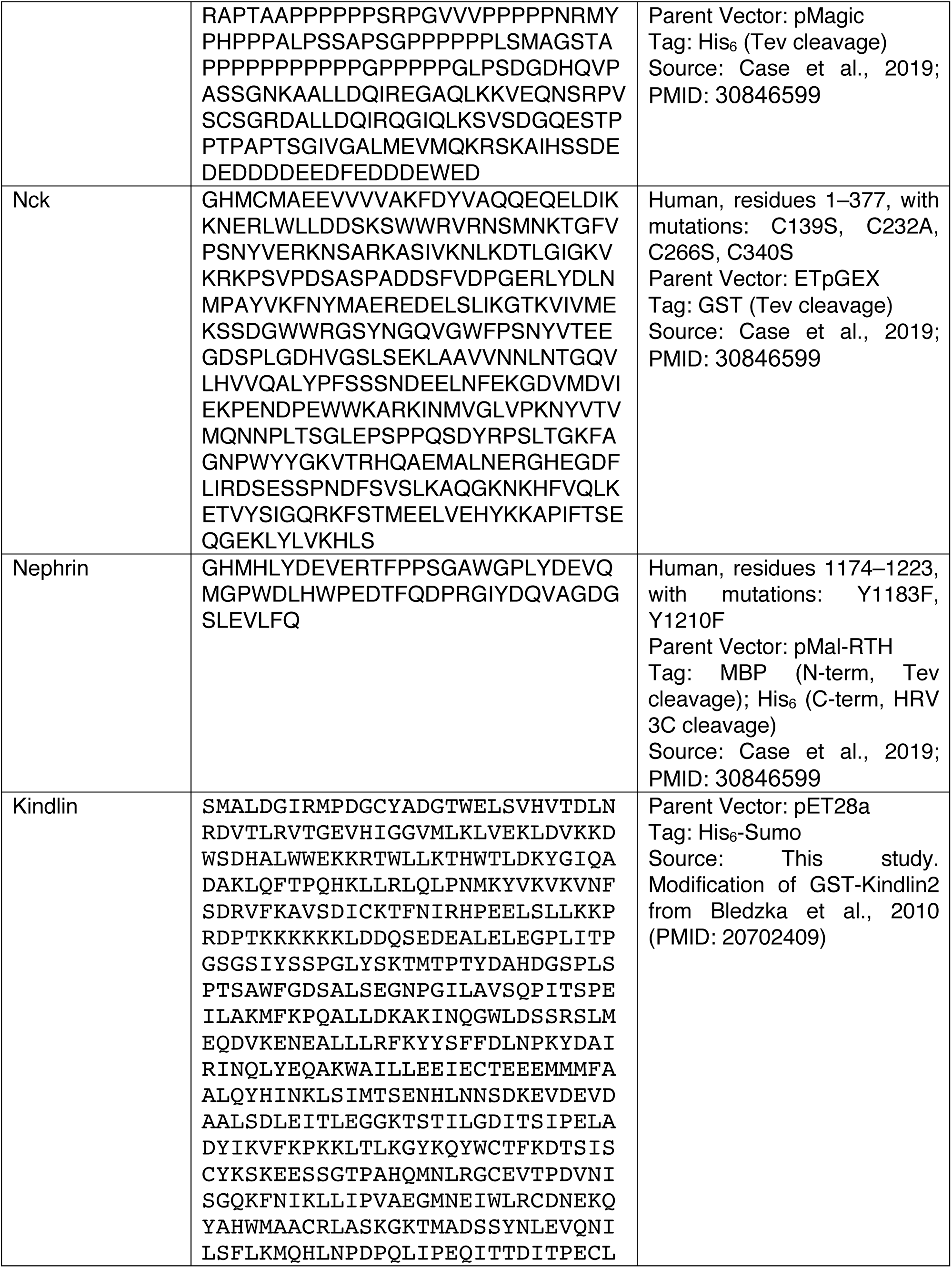
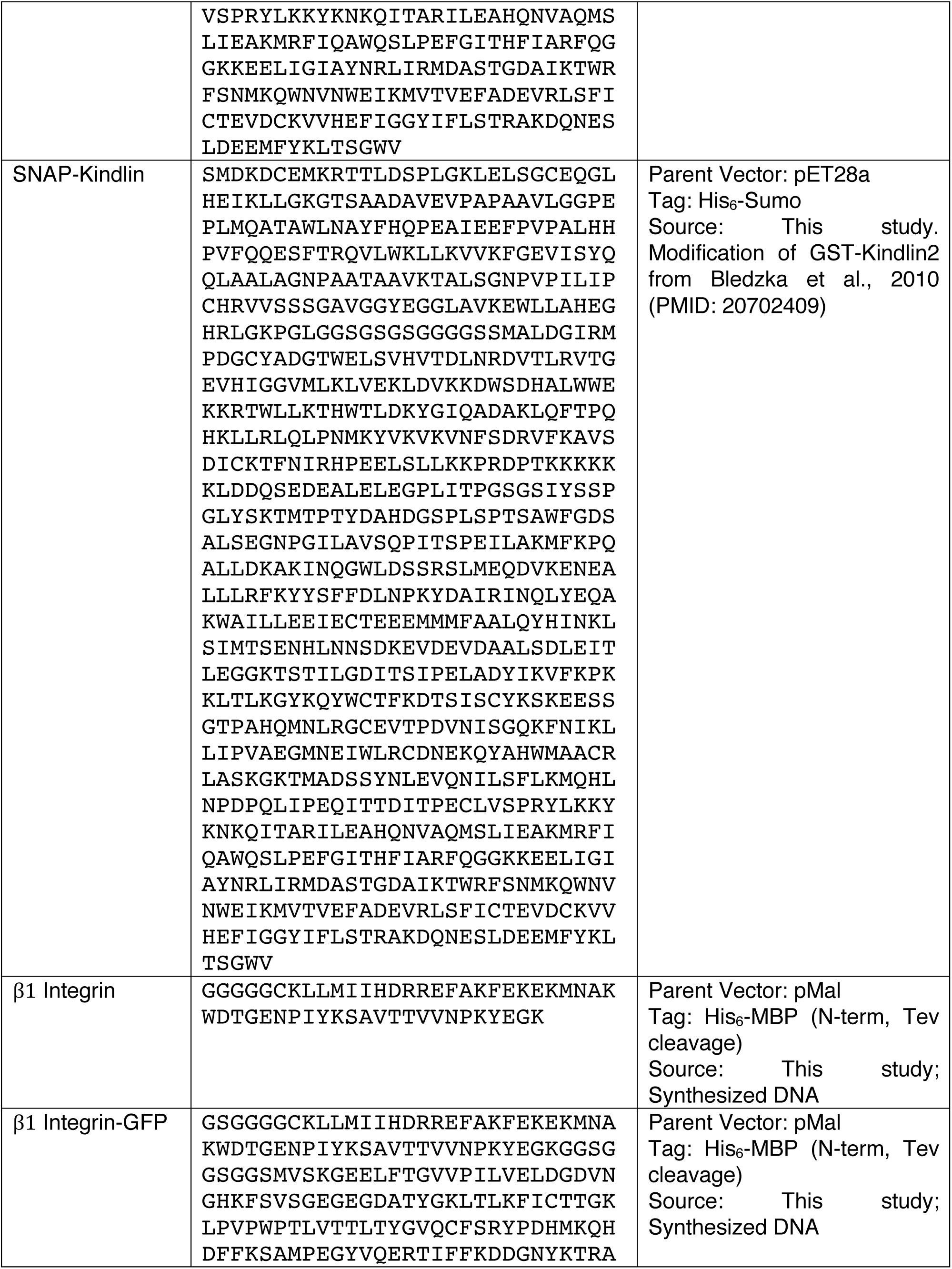
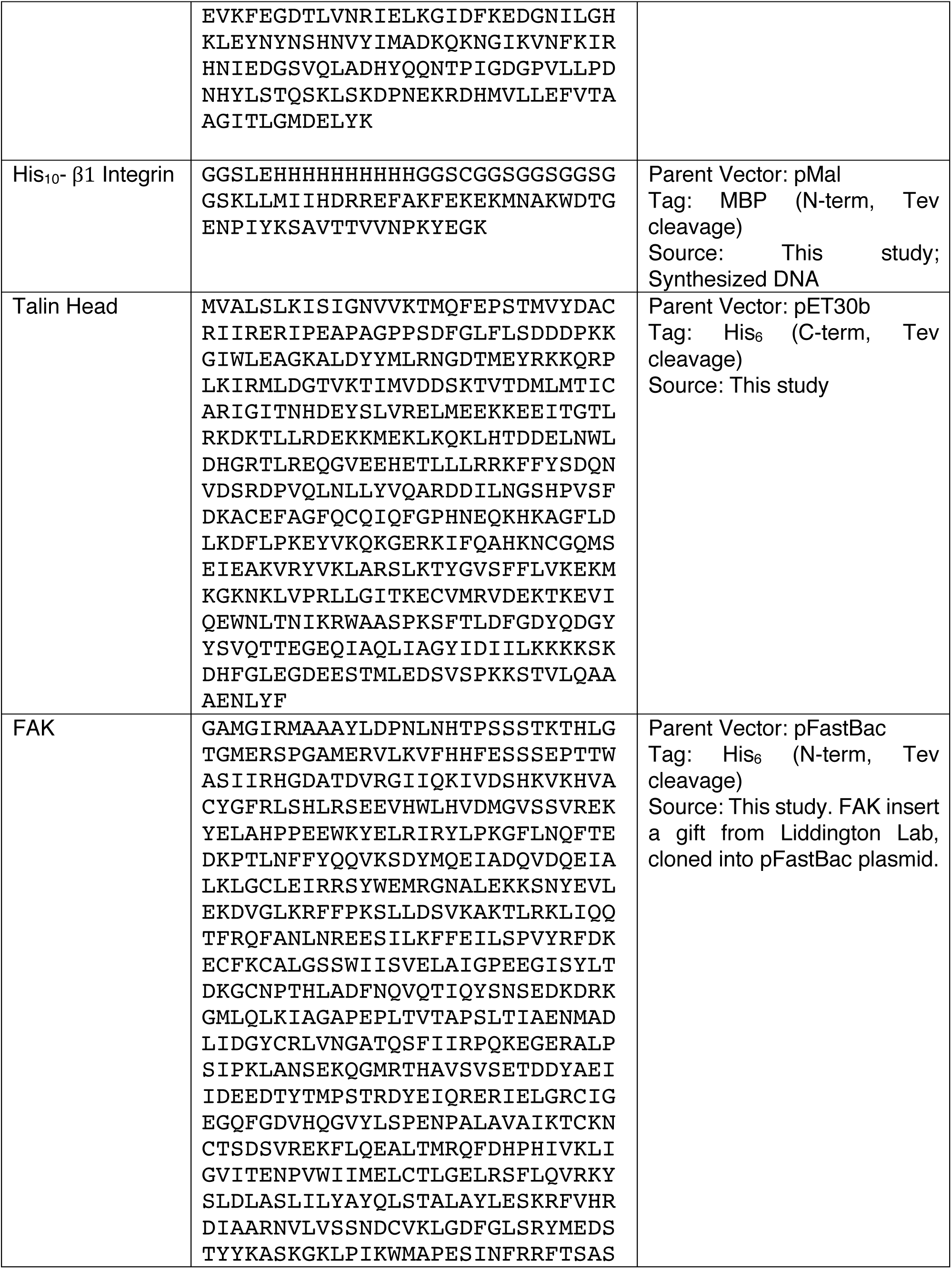
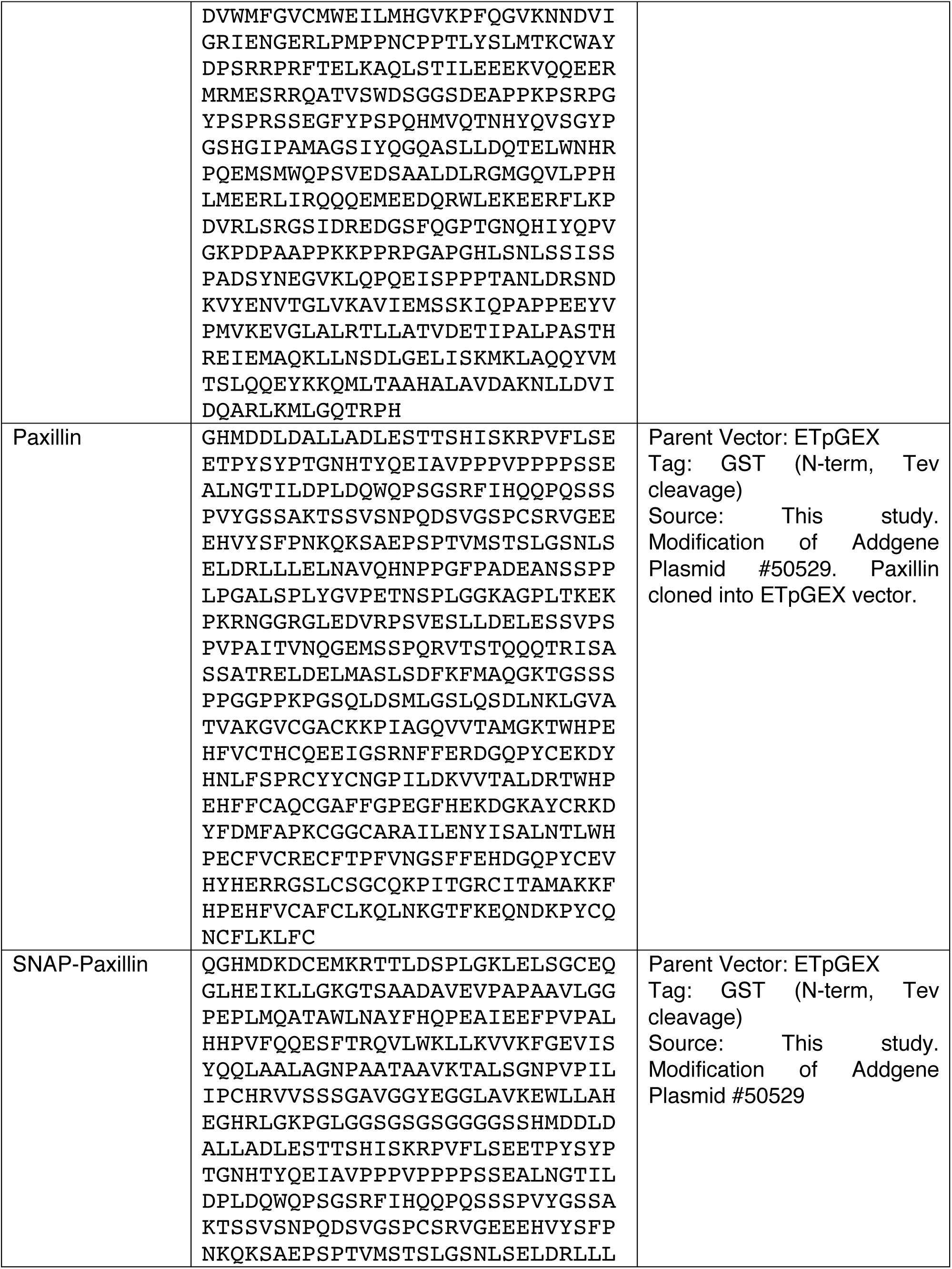
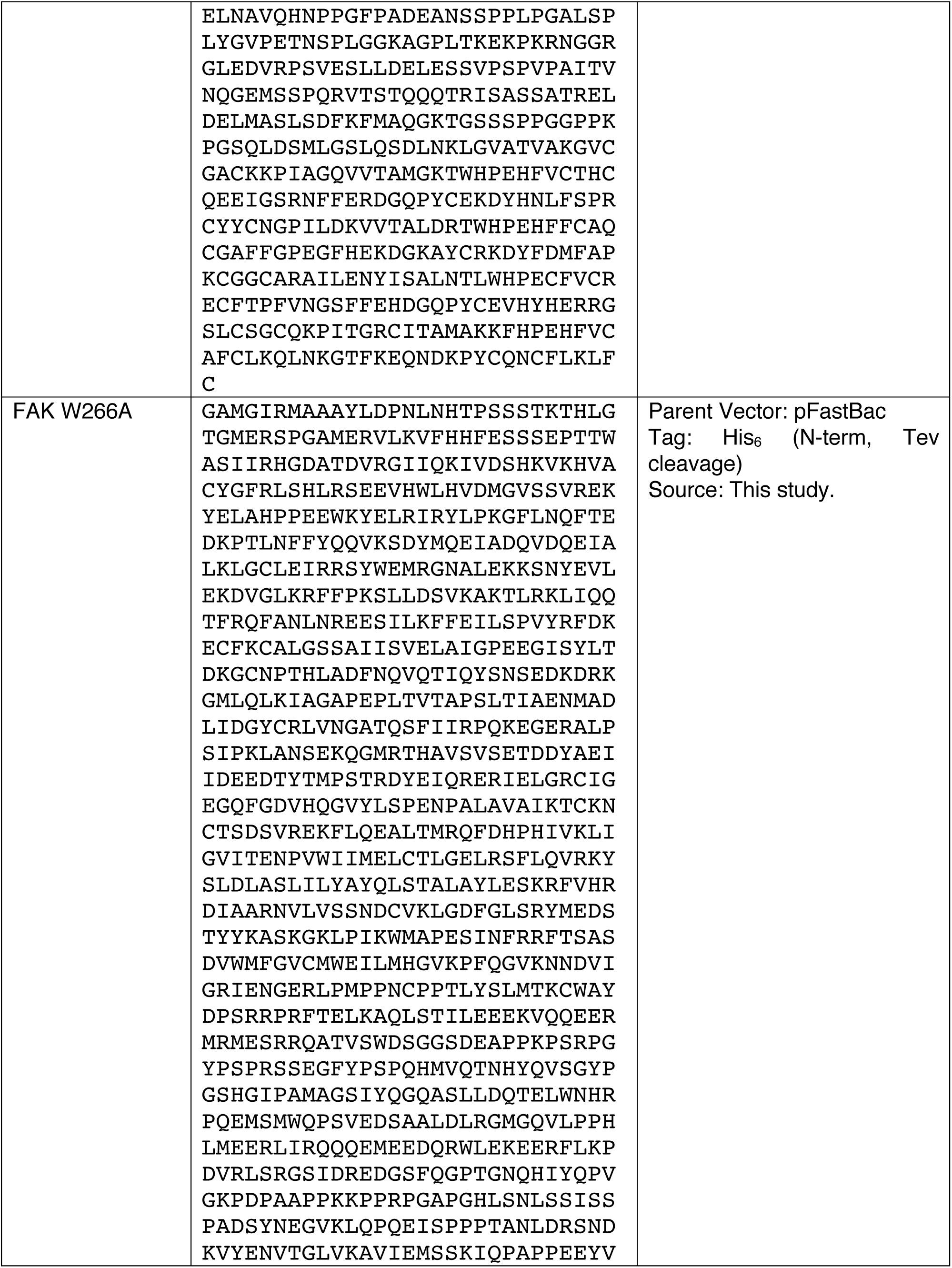
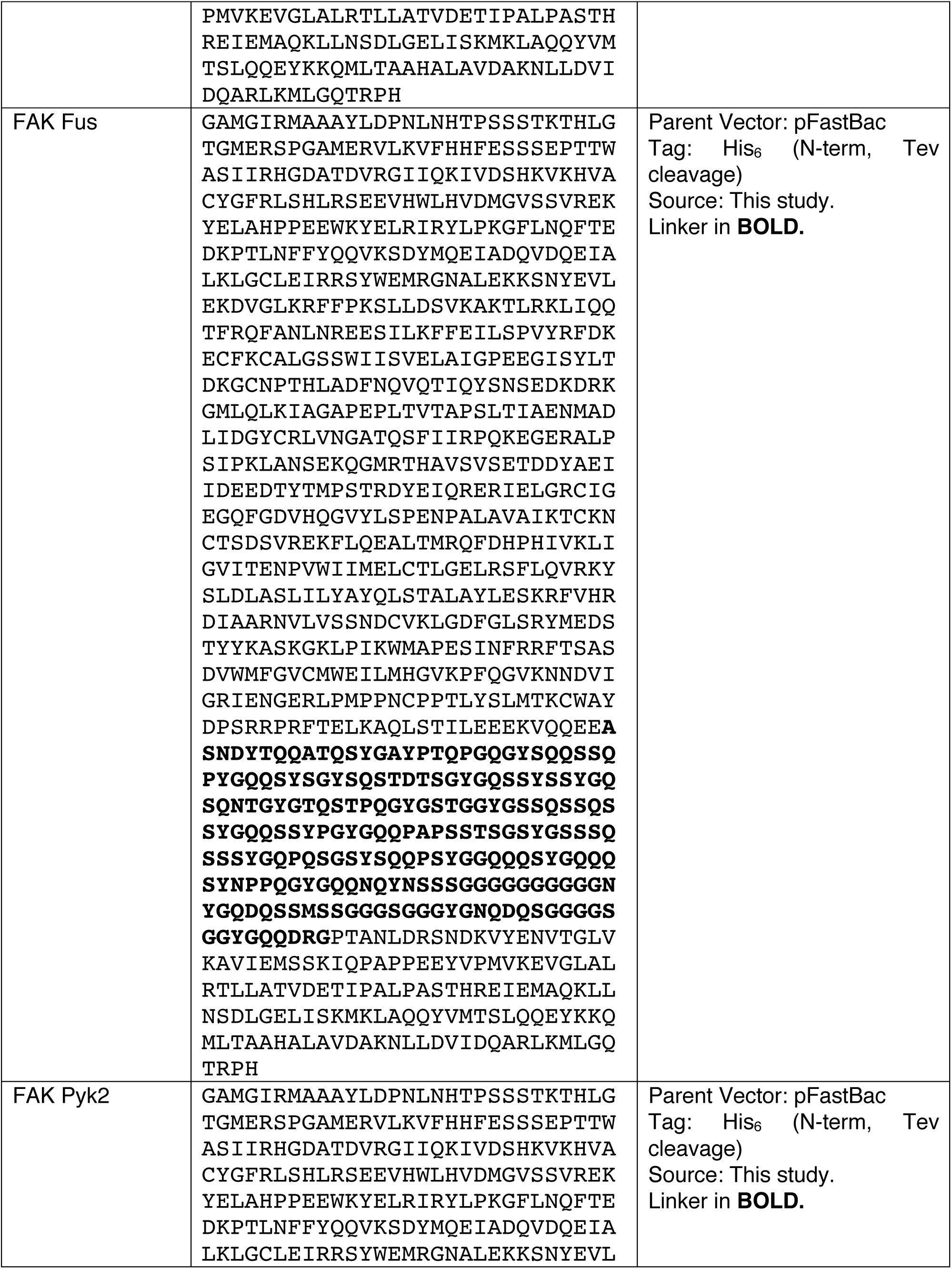
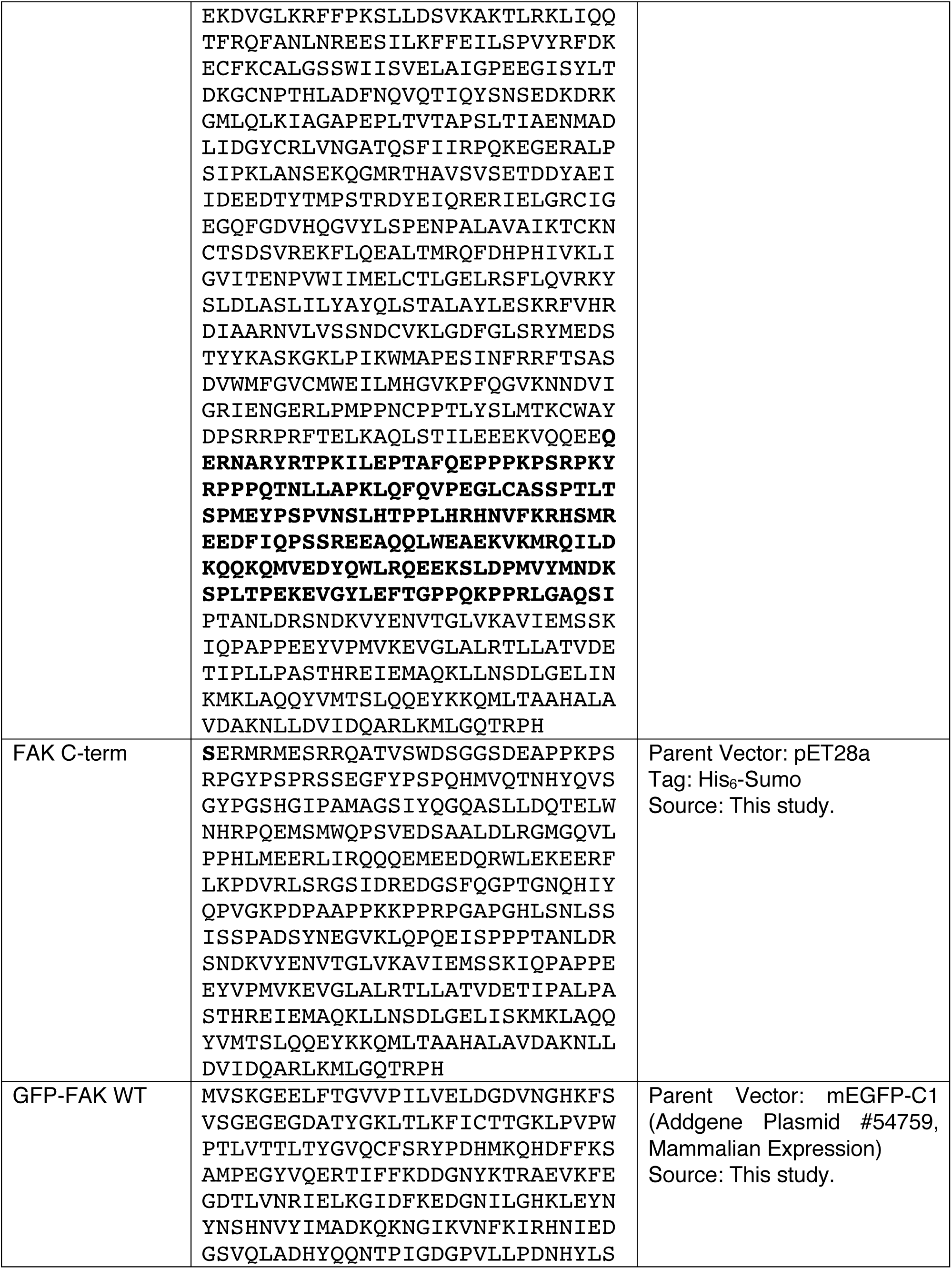
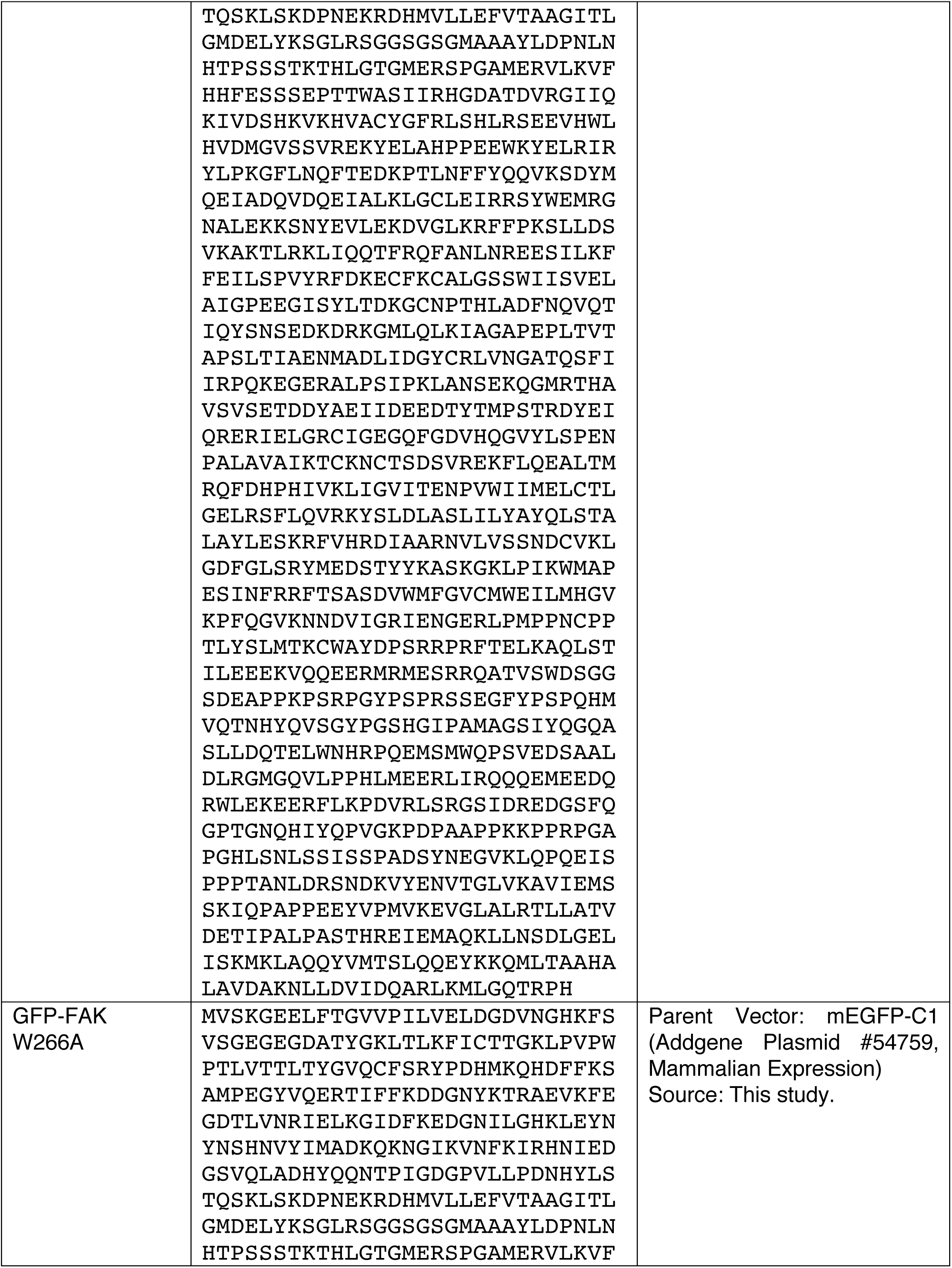
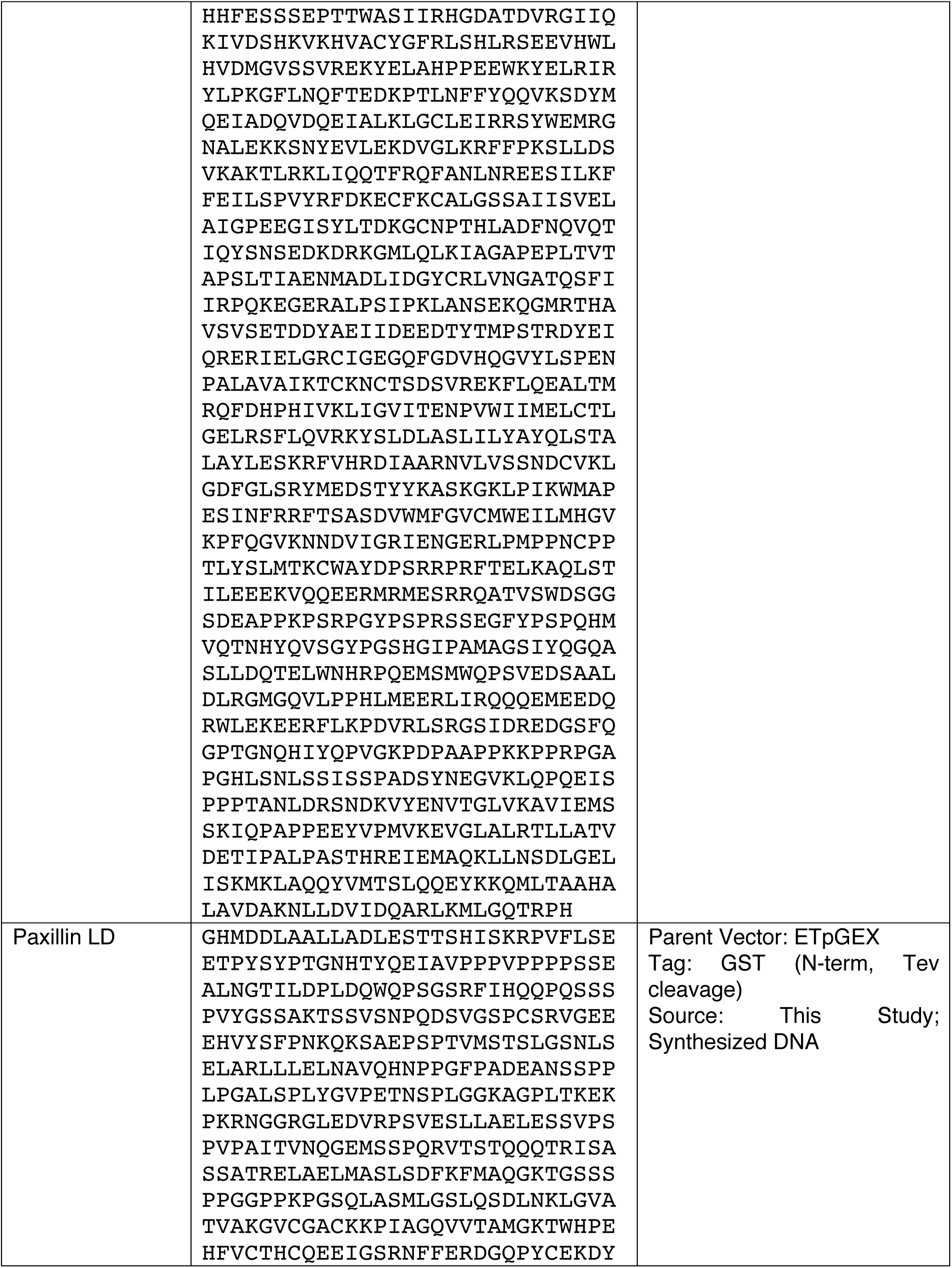
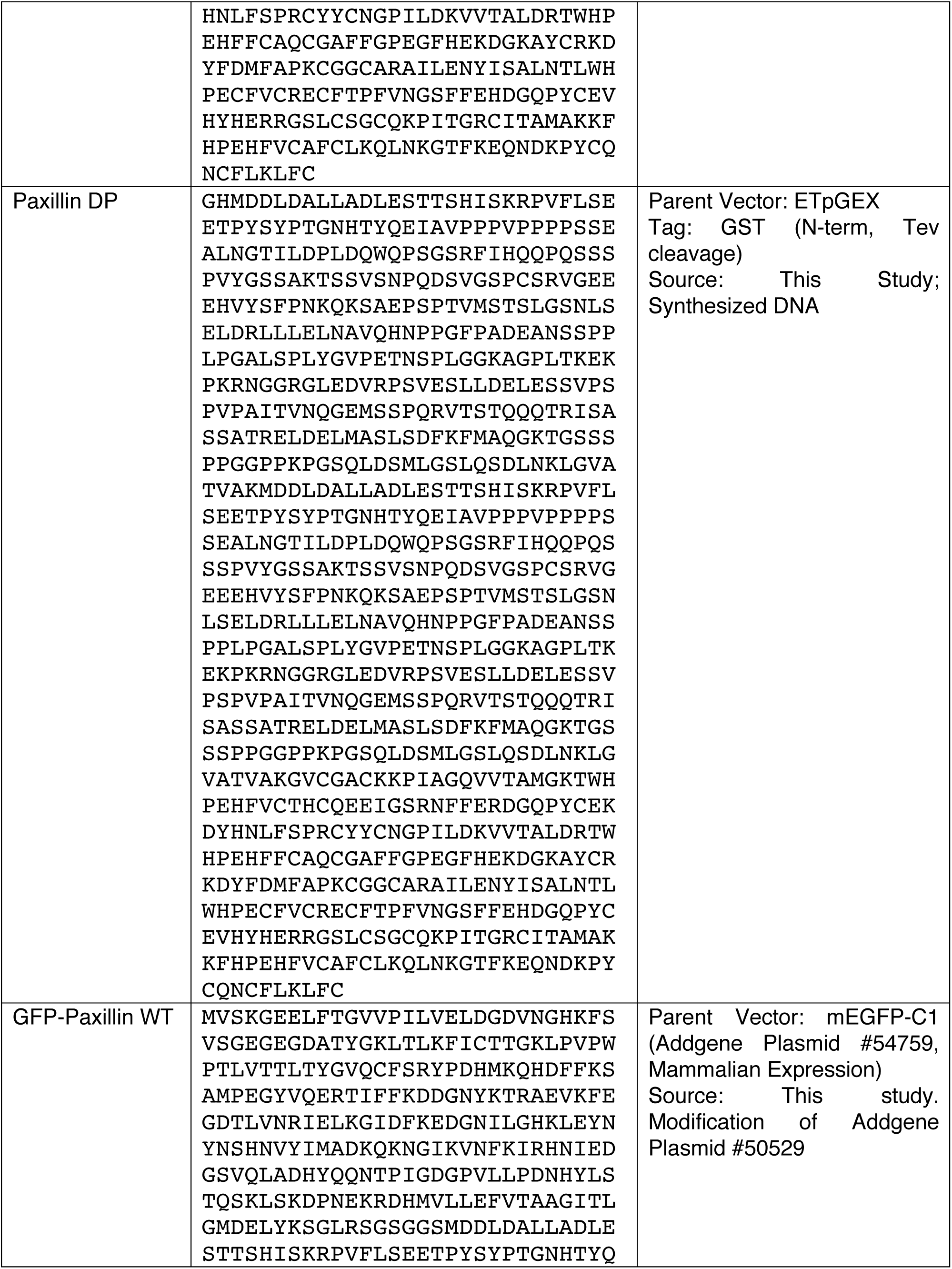
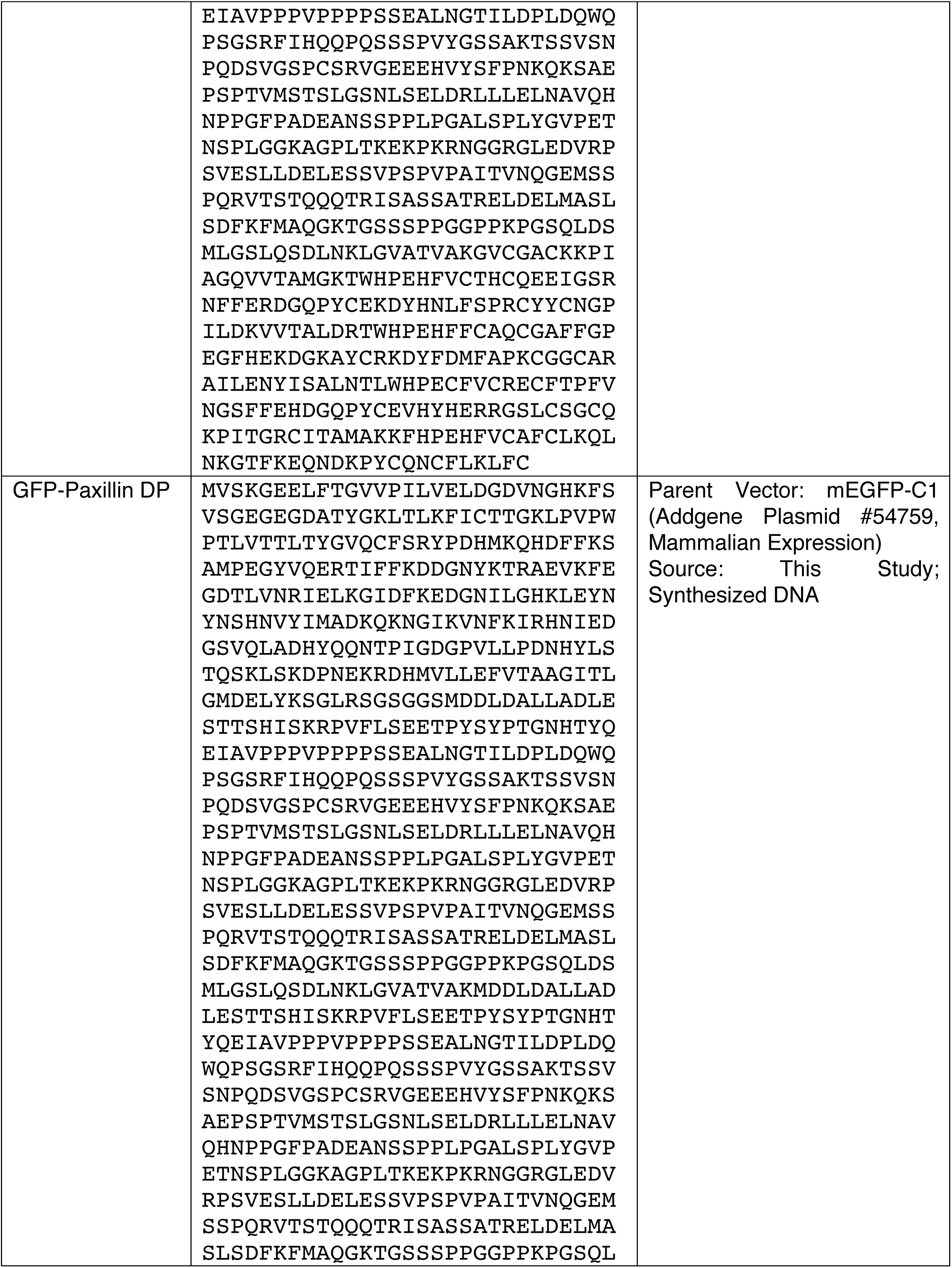
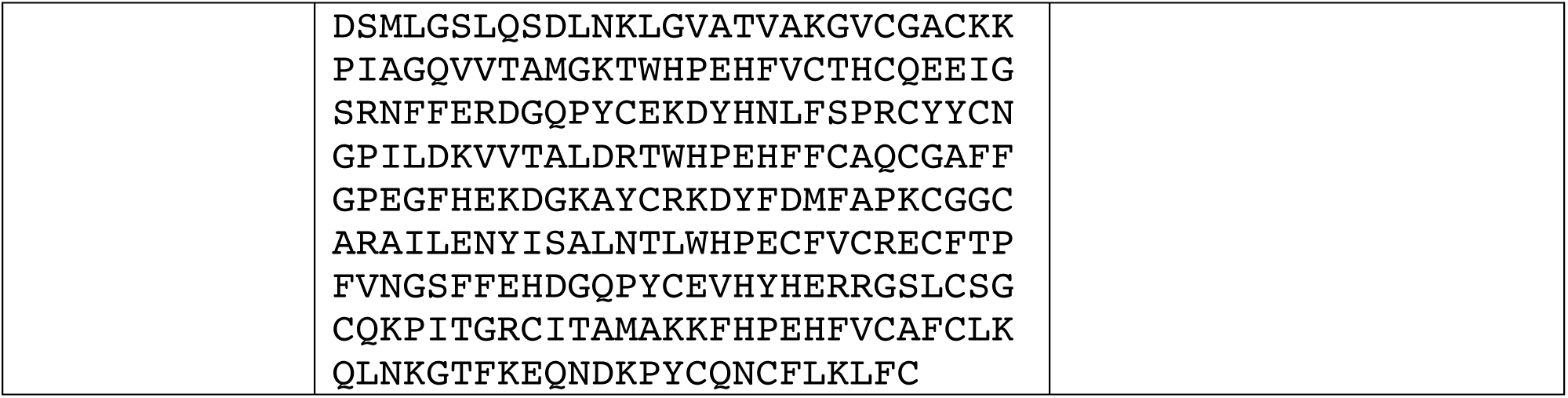
Recombinant DNA used in this study DNA used for expression in bacteria, SF9 cells (pFastBac vector), or mammalian cells (mEGFP-C1 vector). For bacteria and SF9 cell plasmids, the protein sequence of final protease-cleaved and purified protein is shown. For mammalian expression plasmids, the expressed protein sequence is shown.

**Table S7.**
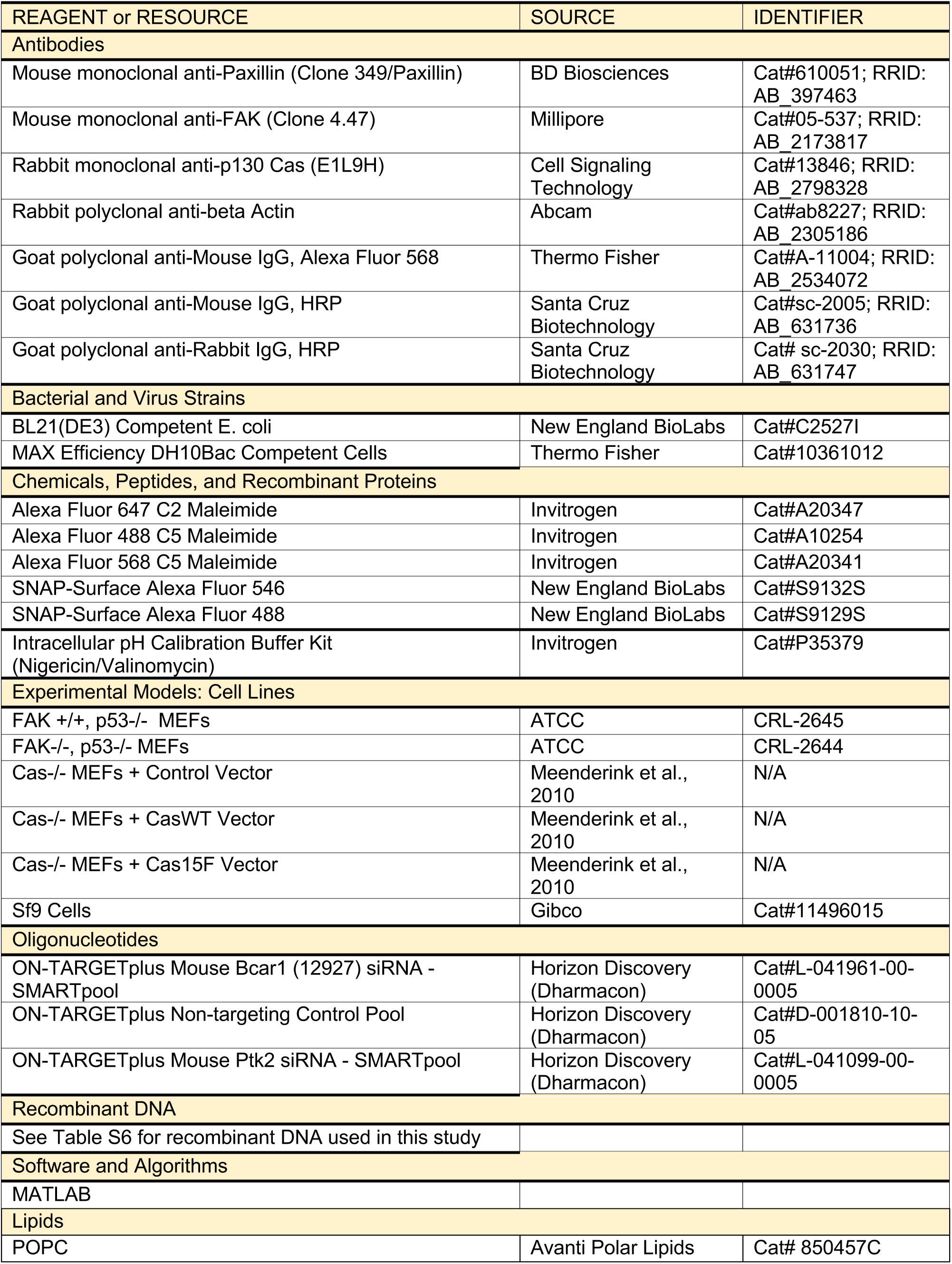
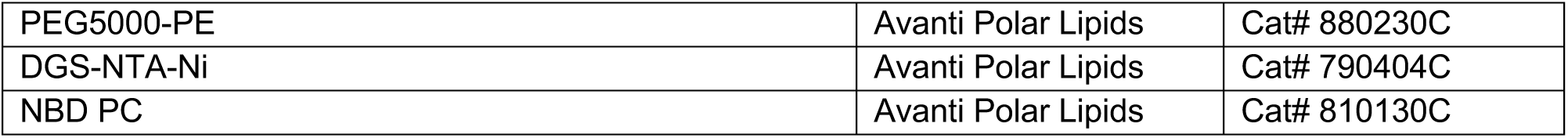
Key Resource Table

## Source Data

**Figure1-figure supplement2-sourcedata.** Original SDS-PAGE gels for Figure 1-Figure Supplement 2a. Folder contains 7 unprocessed images of gels and a file with all 7 uncropped, labeled gels with the lanes included in the figure outlined by black rectangles.

**Figure3-figure supplement2 - source data.** Original SDS-PAGE gel for Figure 3 – figure supplement 2c-d. Folder contains 1 unprocessed image of gel and a file with the uncropped gel with the lanes included in the figure outlined by black rectangles.

**Figure6-figure suppelment2-sourcedata.** Original images of western blots from Figure 6 – figure supplement 2a-b. A) Folder contains 4 unprocessed images of blots and a file with all the uncropped, labeled blots with the lanes included in the figure outlined by black rectangles. B) Folder contains 3 unprocessed images of blots and a file with all the uncropped, labeled blots with the lanes included in the figure outlined by black rectangles.

**Figure7-figure suppelment1-sourcedata.** Original images of western blots from Figure 7 – figure supplement 1a. Folder contains 3 unprocessed images of blots and a file with all the uncropped, labeled blots with the lanes included in the figure outlined by black rectangles.

**Figure7-sourcedata.** Original images of western blots from Figure 7a. Folder contains 3 unprocessed images of blots and a file with all the uncropped, labeled blots with the lanes included in the figure outlined by black rectangles.

## Materials and Methods

### Protein expression, purification, and modification

Information on different recombinant protein constructs is provided in Table S6.

#### Lck purification

His6-Lck was expressed from baculovirus in Spodoptera frugiperta (Sf9) cells. Cells were collected by centrifugation and lysed by douncing on ice in 50 mM Tris-HCl (pH 7.5), 100 mM NaCl, 5 mM βME, 0.01% NP-40, 1 mM PMSF, 1 μg/ml antipain, 1 μg/ml benzamidine, and 1 μg/ml leupeptin. Centrifugation-cleared lysate was applied to Ni-NTA agarose beads (Qiagen), washed with 20 mM Tris-HCl (pH 7.5), 1M NaCl, 20 mM imidazole (pH 7.5), 5 mM βME, and 10% glycerol, and then eluted with 20 mM Tris-HCl (pH 7.5), 100 mM NaCl, 200 mM imidazole (pH 7.5), 5 mM βME, and 10% glycerol. The elute was applied to a Source 15 Q anion exchange column and eluted with a gradient of 100 à 300 mM NaCl in 25 mM HEPES (pH 7.5) and 2 mM βME. Collected fractions were concentrated (Amicon 10 K, Millipore) and applied to an SD75 column in 25 mM HEPES (pH 7.5), 150 mM NaCl, and 1 mM βME.

#### p130Cas purification

His_6_-p130Cas was expressed from baculovirus in Spodoptera frugiperta (Sf9) cells. Cells were collected by centrifugation and lysed by douncing on ice in 20 mM Tris-HCl (pH 8.0), 20 mM Imidazole (pH 8.0), 500 mM NaCl, 10% glycerol and 5 mM βME + cOmplete, EDTA-free Protease Inhibitor tablet (Roche). Centrifugation-cleared lysate was applied to Ni-NTA agarose beads (Qiagen), washed with 20 mM Tris-HCl (pH 8.0), 20 mM Imidazole (pH 8.0), 500 mM NaCl, 10% glycerol and 5 mM βME, and then eluted with 20 mM Tris-HCl (pH 8.0), 400 mM Imidazole (pH 8.0), 500 mM NaCl, 10% glycerol and 5 mM βME. The his tag was removed using TEV protease treatment for 16 hrs at 4°C. Cleaved protein was applied to a Source 15 Q anion exchange column and eluted with a gradient of 100 à 300 mM NaCl in 20 mM Immidazole (pH 7.0), 1 mM DTT and 10% glycerol. Collected fractions were concentrated (Amicon 50 K, Millipore) and applied to an SD200 column in 25mM Hepes pH7.5, 150mM NaCl and 10% glycerol. Cas was concentrated using Amicon Ultra Centrifugal Filter units (Millipore) to >400 µM, mixed with 100 mM HEPES (pH 7.5), 100 mM NaCl, 15 mM ATP, 20 mM MgCl_2_, 2 mM DTT, and 150 nM active Lck, and incubated for 16 hrs at 30**°**C. Phosphorylated Cas was resolved on a Mono Q anion exchange column using a shallow 100 mM à 350 mM NaCl gradient in 40mM Imidazole (pH 7.0), 1mM DTT and 10% glycerol. Fully phosphorylated Cas was additionally purified by size exclusion chromatography to remove any protein aggregates using a Superdex 200 prepgrade column (GE Healthcare) in 25 mM HEPES (pH 7.5), 150 mM NaCl, 1 mM βME, and 10% glycerol. Cas phosphorylation was confirmed by size shift in SDS-PAGE gel and quantified with mass spectrometry.

#### Nephrin purification

BL21(DE3) cells expressing MBP-His_8_-Neprhin were collected by centrifugation and lysed by cell disruption (Emulsiflex-C5, Avestin) in 20 mM imidazole (pH 8.0), 150 mM NaCl, 5 mM βME, 0.1% NP-40, 10% glycerol, 1 mM PMSF, 1 μg/ml antipain, 1 μg/ml benzamidine, 1 μg/ml leupeptin, and 1 μg/ml pepstatin. Centrifugation-cleared lysate was applied to Ni-NTA agarose (Qiagen), washed with 50 mM imidazole (pH 8.0), 150 mM NaCl, 5 mM βME, 0.01% NP-40, and 10% glycerol, and eluted with 500 mM imidazole (pH 8.0), 150 mM NaCl, 5 mM βME, 0.01% NP-40, and 10% glycerol. The MBP tag was removed using TEV protease treatment for 16 hrs at 4°C (for non his-tagged nephrin, the his6-tag was also removed with precision protease treatment). Cleaved protein was applied to a Source 15 Q anion exchange column and eluted with a gradient of 150 mMà350 mM NaCl in 20 mM Immidazole (pH 8.0) and 2 mM DTT followed by size exclusion chromatography using a Superdex 200 prepgrade column (GE Healthcare) in 25 mM HEPES (pH 7.5), 150 mM NaCl, and 2 mM DTT. Nephrin was concentrated using Amicon Ultra Centrifugal Filter units (Millipore) to >400 µM, mixed with 100 mM HEPES (pH 7.5), 100 mM NaCl, 15 mM ATP, 20 mM MgCl_2_, 2 mM DTT, and 150 nM active Lck, and incubated for 16 hrs at 30**°**C. Phosphorylated Nephrin (pNephrin) was resolved on a Mono Q anion exchange column using a shallow 100 mM à 350 mM NaCl gradient in 20 mM imidazole (pH 8.0) and 2 mM DTT to separate differentially phosphorylated species of Neprhin. Fully phosphorylated pNephrin was additionally purified by size exclusion chromatography to remove any protein aggregates using a Superdex 200 prepgrade column (GE Healthcare) in 25 mM HEPES (pH 7.5), 150 mM NaCl, 5 mM βME, and 10% glycerol. Complete Nephrin phosphorylation was confirmed by mass spectrometry.

#### Nck purification

BL21(DE3) cells expressing GST-Nck were collected by centrifugation and lysed by sonication in 25 mM Tris-HCl (pH 8.0), 200 mM NaCl, 2 mM EDTA (pH 8.0), 1 mM DTT, 1 mM PMSF, 1 μg/ml antipain, 1 μg/ml benzamidine, 1 μg/ml leupeptin, and 1 μg/ml pepstatin. Centrifugation-cleared lysate was applied to Glutathione Sepharose 4B (GE Healthcare) and washed with 20 mM Tris-HCl (pH 8.0), 200 mM NaCl, and 1 mM DTT. GST was cleaved from protein by TEV protease treatment for 16 hrs at 4°C. Cleaved protein was applied to a Source 15 Q anion exchange column and eluted with a gradient of 5 à 250 mM NaCl in 20 mM imidazole (pH 7.0) and 1 mM DTT. Eluted protein was pooled and applied to a Source 15 S cation exchange column and eluted with a gradient of 0 à 500 mM NaCl in 20 mM imidazole (pH 7.0) and 1 mM DTT. Eluted protein was concentrated using Amicon Ultra 10k concentrators and further purified by size exclusion chromatography to remove any protein aggregates using a Superdex 75 prepgrade column (GE Healthcare) in 25 mM HEPES (pH 7.5), 150 mM NaCl, and 1 mM βME.

#### N-WASP purification

BL21(DE3) cells expressing His_6_-N-WASP were collected by centrifugation and lysed by cell disruption (Emulsiflex-C5, Avestin) in 20 mM imidazole (pH 7.0), 300 mM KCl, 5 mM βME, 0.01% NP-40, 1 mM PMSF, 1 μg/ml antipain, 1 μg/ml benzamidine, 1 μg/ml leupeptin, and 1 μg/ml pepstatin. The cleared lysate was applied to Ni-NTA agarose (Qiagen), washed with 50 mM imidazole (pH 7.0), 300 mM KCl, 5 mM βME, and eluted with 300 mM imidazole (pH 7.0), 100 mM KCl, and 5 mM βME. The elute was further purified over a Source 15 Q column using a gradient of 250 → 450 mM NaCl in 20 mM imidazole (pH 7.0), and 1 mM DTT. The His_6_-tag was removed by TEV protease at 4°C for 16 hr (for His-N-WASP, no TEV treatment occurred). Cleaved N-WASP (or uncleaved for His-N-WASP) was then applied to a Source 15 S column using a gradient of 110 → 410 mM NaCl in 20 mM imidazole (pH 7.0), 1 mM DTT. Fractions containing N-WASP were concentrated using an Amicon Ultra 10k concentrator (Millipore) and further purified by size exclusion chromatography to remove any protein aggregates using a Superdex 200 prepgrade column (GE Healthcare) in 25 mM HEPES (pH 7.5), 150 mM KCl, 1 mM βME, and 10% glycerol.

#### Kindlin Purification

BL21(DE3) cells expressing His_6_-sumo-kindlin were collected by centrifugation and lysed by cell disruption (Emulsiflex-C5, Avestin) in 20 mM Tris-HCl (pH 8.0), 20 mM Imidazole (pH 8.0), 150 mM NaCl, 0.01% NP-40, 5 mM βME, 1 μg/ml antipain, 1 μg/ml benzamidine, 1 μg/ml leupeptin, and 1 μg/ml pepstatin. The cleared lysate was applied to Ni-NTA agarose (Qiagen) and washed with 20 mM Tris-HCl (pH 8.0), 20 mM Imidazole (pH 8.0), 150 mM NaCl, 0.01% NP-40, 5 mM βME. Kindlin was eluted with 20 mM Tris-HCl (pH 8.0), 300 mM Imidazole (pH 8.0), 150 mM NaCl, 0.01%NP-40, 5 mM βME. The elute was further purified over a Source 15 Q column using a gradient of 0 → 300 mM NaCl in 20 mM Tris-HCl (pH 8.0), 1 mM DTT. The Collected fractions were pooled and the His_6_-sumo tag was removed by Ulp1 sumo protease for 2 hr at room temperature. Protein was concentrated using an Amicon Ultra 10k concentrators and further purified by size exclusion chromatography to remove any protein aggregates using a Superdex 200 column (GE Healthcare) in 25 mM HEPES (pH 7.5), 300 mM NaCl, 10% glycerol, and 1 mM DTT.

#### Talin Head Purification

BL21(DE3) cells expressing talin head-His_6_ were collected by centrifugation and lysed by cell disruption (Emulsiflex-C5, Avestin) in 20 mM Tris-HCl (pH 8.0), 5 mM Imidazole (pH 8.0), 500 mM NaCl, 1% TritonX, 5 mM βME, 1 μg/ml antipain, 1 μg/ml benzamidine, 1 μg/ml leupeptin, and 1 μg/ml pepstatin. Centrifugation-cleared lysate was applied to Ni-NTA agarose (Qiagen), washed with 20 mM Tris-HCl (pH 8.0), 30 mM Imidazole (pH 8.0), 500 mM NaCl, 5 mM βME, 1 μg/ml benzamidine, and eluted with 20 mM Tris-HCl (pH 8.0), 100 mM Imidazole (pH 8.0), 500 mM NaCl, 5 mM βME, 1 μg/ml benzamidine. The his6-tag was removed using TEV protease treatment for 16 hrs at 4°C. Cleaved protein was applied to a Source 15 S cation exchange column and eluted with a gradient of 150 mMà500 mM NaCl in 20 mM Immidazole (pH 7.0), 10% glycerol, and 2 mM DTT followed by size exclusion chromatography to remove any protein aggregates using a Superdex 75 column (GE Healthcare) in 25 mM HEPES (pH 7.5), 150 mM NaCl, and 2 mM DTT.

#### Integrin Purification

BL21(DE3) cells expressing integrin (MBP-his_10_-Integrin, MBP-his_6_-Integrin, or MBP-his_6_- Integrin-GFP) were collected by centrifugation and lysed by cell disruption (Emulsiflex-C5, Avestin) in 20 mM Tris-HCl (pH 8.0), 20 mM Imidazole (pH 8.0), 150 mM NaCl, 0.01% NP-40, 10% glycerol, 5 mM βME, 1 μg/ml antipain, 1 μg/ml benzamidine, 1 μg/ml leupeptin, and 1 μg/ml pepstatin. Centrifugation-cleared lysate was applied to Ni-NTA agarose (Qiagen), washed with 20 mM Tris-HCl (pH 8.0), 20 mM Imidazole (pH 8.0), 150 mM NaCl, 10% glycerol, 5 mM βME, and eluted with 20 mM Tris-HCl (pH 8.0), 300 mM Imidazole (pH 8.0), 150 mM NaCl, 10% glycerol, 5 mM βME. Eluate was applied to a Source 15Q anion exchange column and eluted with a gradient of 0 -300 mM NaCl in 20 mM Tris-HCl (pH 8.0), 10% glycerol and 1 mM DTT. The MBP-tag or MBP-his_6_-tag was removed using TEV protease treatment for 16 hrs at 4°C. Cleaved protein was applied to a Source 15 S cation exchange column and eluted with a gradient of 150 mMà500 mM NaCl in 20 mM Bis-Tris (pH 6.0), 10% glycerol, and 1 mM DTT followed by size exclusion chromatography to remove any protein aggregates using a Superdex 75 column (GE Healthcare) in 25mM HEPES (pH 7.5), 1M NaCl, 10% glycerol, 1 mM βME.

#### FAK purification

His_6_-FAK was expressed from baculovirus in Spodoptera frugiperta (Sf9) cells. Cells were collected by centrifugation and lysed by douncing on ice in 25 mM HEPES (pH 7.5), 20 mM Imidazole (pH 7.5), 500 mM NaCl, 10% glycerol and 5 mM βME + cOmplete(TM), EDTA-free Protease Inhibitor tablet. Centrifugation-cleared lysate was applied to Ni-NTA agarose beads (Qiagen), washed with 25 mM HEPES (pH 7.5), 20 mM Imidazole (pH 7.5), 500 mM NaCl, 10% glycerol, 5 mM βME, 1 μg/ml benzamidine, and eluted with 25 mM HEPES (pH 7.5), 400 mM Imidazole (pH 7.5), 1 M NaCl, 10% glycerol, 5 mM βME, 1 μg/ml benzamidine. The his6-tag was removed using TEV protease treatment for 16 hrs at 4°C. Cleaved protein was further purified with size exclusion chromatography to remove any protein aggregates applied to a Superdex 200 column (GE Healthcare) in 25 mM HEPES (pH 7.5), 300 mM NaCl, and 1 mM DTT.

#### Paxillin purification

BL21(DE3) cells expressing GST-Paxillin were collected by centrifugation and lysed by cell disruption (Emulsiflex-C5, Avestin) in 20 mM Tris-HCl (pH 8.0), 300 mM NaCl, 0.01% NP-40, 10% glycerol, 1mM DTT, 1 μg/ml antipain, 1 μg/ml benzamidine, 1 μg/ml leupeptin, and 1 μg/ml pepstatin. Centrifugation-cleared lysate was applied to Glutathione Sepharose 4B (GE Healthcare) and washed with 20 mM Tris-HCl (pH 8.0), 300 mM NaCl, 0.01% NP-40, 10% glycerol, 1mM DTT, 1 μg/ml benzamidine. GST was cleaved from protein by TEV protease treatment for 16 hrs at 4°C. Cleaved protein was applied to a Source 15 Q anion exchange column and eluted with a gradient of 5 à 500 mM NaCl in 20 mM imidazole (pH 8.0), 10% glycerol and 1 mM DTT. Eluted protein was concentrated using Amicon Ultra 10k concentrators and further purified by size exclusion chromatography to remove any protein aggregates using a Superdex 200 column (GE Healthcare) in 25 mM HEPES (pH 7.5), 150 mM NaCl, and 1 mM βME.

#### FAK c-term purification

BL21(DE3) cells expressing His_6_-sumo-FAK c-term were collected by centrifugation and lysed by cell disruption (Emulsiflex-C5, Avestin) in 20 mM Tris-HCl (pH 8.0), 20 mM Imidazole (pH 8.0), 300 mM NaCl, 0.01% NP-40, 5 mM βME, 1 μg/ml antipain, 1 μg/ml benzamidine, 1 μg/ml leupeptin, and 1 μg/ml pepstatin. Centrifugation-cleared lysate was applied to Ni-NTA agarose (Qiagen), washed 20 mM Tris-HCl (pH 8.0), 300 mM Imidazole (pH 8.0), 300 mM NaCl, 0.01% NP-40, 5 mM βME, and eluted with 20 mM Tris-HCl (pH 8.0), 20 mM Imidazole (pH 8.0), 300 mM NaCl, 5 mM βME,. Eluate was applied to a Source 15 Q anion exchange column and eluted with a gradient of 150 mMà300 mM NaCl in 20 mM Tris-HCl (pH 8.0), 10% glycerol, and 1 mM DTT. Collected fractions were pooled and the His_6_-sumo tag was removed by Ulp1 sumo protease for 2 hr at room temperature. Protein was concentrated using an Amicon Ultra 10k concentrators and further purified by size exclusion chromatography to remove any protein aggregates using a Superdex 75 column (GE Healthcare) in 25 mM HEPES (pH 7.5), 300 mM NaCl, and 1 mM DTT.

#### Fluorophore conjugation

For conjugation with Maleimide chemistry (Nck, N-WASP, Integrin, FAK) recombinant proteins to be labeled with Alexa fluorophores were concentrated using Amicon Ultra Centrifugal Filter units (Millipore) to ∼100 μM. 5 mM βME was added to reduce cysteine residues followed by buffer exchange using a HiTrap 26/10 Desalting column (GE Healthcare) in 25 mM HEPES (pH 7.5) and 150 mM NaCl. Fractions containing protein were collected and concentrated to 100 μM. 500 μM Alexa Fluor 647 C_2_ Maleimide (ThermoFisher, for N-WASP), Alexa Fluor 488 C_5_ Maleimide (ThermoFisher, for his_8_-pNephrin, N-WASP, and his_8_-N-WASP) or Alexa Fluor 568 C_5_ Maleimide (ThermoFisher, for Nck) was added, and the reaction was incubated with gentle mixing at 4 °C for 16 hr. The reaction was quenched with 1 μl 14.3 M βME followed by final buffer exchange using size exclusion chromatography (GE Healthcare) in 25 mM HEPES (pH 7.5), 150 mM NaCl, and 1 mM βME. We consistently achieve > 98% labeling efficiency.

For conjugation of SNAP-tagged proteins (paxillin, kindlin) recombinant proteins to be labeled with fluorophores were concentrated using Amicon Ultra Centrifugal Filter units (Millipore) to ∼50 μM. 1 mM DTT and 2-fold molar excess SNAP-tag Substrate (SNAP-Surface Alexa488 or SNAP-Surface Alexa546) were added. The mixture was incubated for 30 min at 37°C. Labeled proteins were purified by final buffer exchange using size exclusion chromatography (Superdex 200 column, GE Healthcare) in the appropriate buffer (i.e. final buffer in the above purification protocol). We consistently achieve > 98% labeling efficiency.

Final protein concentration and degree of labeling were calculated from the protein absorbance using the following formulas:

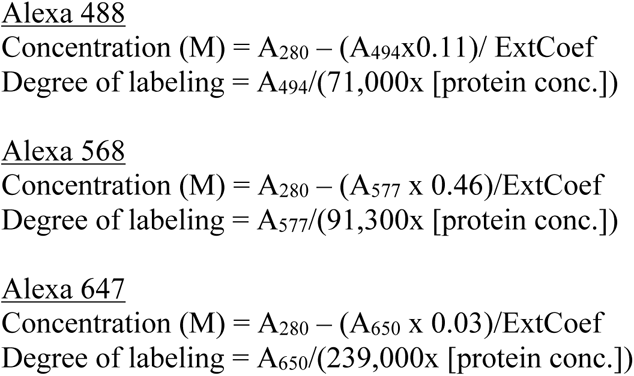

#### Mass spectrometry analysis of p130Cas phosphorylation

Protein samples were analyzed by LC/MS using a Sciex X500B Q-ToF mass spectrometer running Sciex OS v.1.6.1, coupled to an Agilent 1290 Infinity II HPLC. Samples were injected onto a POROS R1 reverse-phase column (2.1 x 30 mm, 20 µm particle size, 4000 Å pore size), desalted, and the amount of buffer B was manually increased stepwise until the protein eluted off the column. Buffer A contained 0.1% formic acid in water and buffer B contained 0.1% formic acid in acetonitrile. The mobile phase flow rate was 300 uL/min. The acquired mass spectra for the protein of interest were deconvoluted using BioPharmaView v. 3.0.1 software (Sciex) in order to obtain the molecular weight.

#### Turbidity Measurements

Unlabeled proteins were diluted in 50 mM HEPES (pH 7.3), 50 mM KCl, 1 mM TCEP (“Buffer A”), and 0.1% (1 mg/mL BSA), unless a different buffer is indicated in figure legends. After a 30-60 min incubation, solution was transferred to Quartz cuvette and Absorbance at 350 nm was measured in a Spectrophotometer (Agilent). While we use 1 mg/mL BSA (0.1%) to prevent nonspecific interactions with surfaces, we note that this is well below the BSA concentration necessary to induce crowding (typically 100 mg/mL or greater). For temperature-controlled measurements, Absorbance at 350 nm was measured using a Cary 100 UV-Visible spectrophotometer equipped with a Peltier thermal controller (Agilent Technologies, Australia). The reaction mixture was incubated in a microcentrifuge tube at the desired temperature for 30 min, and then placed into a pre-equilibrated cuvette and spectrophotometer for measurement.

#### 3D droplet assays

384-well glass-bottomed plates (Brooks) were washed with 5% Hellmanex III (Hëlma Analytics) for 3.5 hrs at 55**°**C and thoroughly rinsed with MilliQ H_2_O. Plates were then washed with 1M NaOH for 1hr at 55**°**C and thoroughly rinsed with MilliQ H_2_O. 50 µL of 20 mg/mL mPEG silane MW 5k (Creative PEGworks) in 95% EtOH was added to each well. The plate was covered in parafilm and incubated overnight at room temperature. The plate was thoroughly rinsed with MilliQ H_2_O, dried, and sealed with foil. Prior to experiment, individual wells were washed 3X with MilliQ H_2_O, blocked with Buffer A containing 0.1% BSA for 30 min at room temperature. For droplet experiments, proteins were combined at 1 µM concentration in Buffer A containing 0.1% BSA and a glucose/glucose oxidase/catalase O_2_-scavenging system. 80 µL of the mixture was added to the well and incubated for 30 min prior to imaging.

#### Small Unilamellar Vesicle Preparation

The general protocol we follow to make small unilamellar vesicles (SUVs) and supported phospholipid bilayers is described in (Su et al., 2017). Synthetic 1-palmitoyl-2-oleoyl-glycero-3-phosphocholine (POPC), 1,2-dioleoyl-*sn*-glycero-3-[(*N*-(5-amino-1-carboxypentyl)iminodiacetic acid)succinyl] (nickel salt, DGS-NTA-Ni), and 1,2-dioleoyl-*sn*-glycero-3-phosphoethanolamine-N-[methoxy(polyethyleneglycol)-5000] (ammonium salt) (PEG5000 PE) were purchased from Avanti Polar Lipids as chloroform suspension. Using glass Hamilton syringes, lipids were mixed to make a chloroform suspension containing 98% POPC, 2% DGS-NTA-Ni and 0.1% PEG5000 PE). Chloroform was evaporated with gentle stream of Argon Gas, desiccated in a vacuum overnight, and resuspended in PBS (pH 7.3) with vortexing. To promote the formation of small unilamellar vesicles (SUVs), the lipid solution was repeatedly frozen in liquid N_2_ and thawed using a 37**°**C water bath until the solution cleared (∼35 freeze-thaw cycles). SUV-containing solution was centrifuged at 33,500g for 45 min at 4**°**C to remove large vesicles. Cleared supernatant containing SUVs was collected and stored at 4**°**C covered with Argon for up to two weeks.

#### Reconstitution on Supported Phospholipid Bilayers

Briefly, 96-well glass-bottomed plates (Brooks) were washed with 5% Hellmanex III (Hëlma Analytics) for 3.5 hrs at 55**°**C. The plate was thoroughly rinsed with MilliQ H_2_O, dried, and sealed with foil. Prior to experiment, individual wells were washed with 6M NaOH for 30 min at 50**°**C two times, and thoroughly rinsed with MilliQ H_2_O followed by equilibration with 50 mM HEPES (pH 7.3), 50 mM KCl, and 1 mM TCEP (“Buffer A”). 12 μL SUVs were added to cleaned wells covered by Buffer A and incubated for 40 min hr at 40**°**C to allow SUVs to collapse on glass and fuse to form the bilayer. Bilayers were washed three times with Buffer A to remove excess SUVs, and then blocked with Buffer A containing 0.1% BSA for 30 min at room temperature. His-tagged proteins (10 nM his10-Integrin) were mixed in Buffer A containing 0.1% BSA, added to phospholipid bilayers, and incubated for 2 hrs. Bilayers were then washed with Buffer A containing 0.1% BSA to remove unbound His-tagged proteins. Additional proteins were added to the well if required for the experiment. Microscopy experiments were performed in the presence of a glucose/glucose oxidase/catalase O_2_-scavenging system to reduce photodamage and photobleaching. Unexpectedly, we found that FAK interacts with Nickel and competes with his-tagged proteins for Ni-lipid binding (data not shown). To reduce this effect, we lowered FAK concentration to 200 nM for experiments on supported phospholipid bilayers. Prior to any experiment, the fluidity of bilayers was indirectly assessed by imaging integrin Alexa-488. We photobleached a 5-micron region, and experiments were only performed if FRAP t_1/2_ < 10 s. To confirm that our methods consistently give fluid, uniform bilayers, we also assessed bilayer fluidity directly in bilayers containing 1% PC-NBD, a fluorescent lipid (Figure 4, supplement 1).

#### Microscopy

TIRF images were captured using a TIRF/iLAS2 TIRF/FRAP module (Biovision) mounted on a Leica DMI6000 microscope base equipped with a plan apo 100 X 1.49 NA TIRF objective and a 405/488/561/647nm Laser Quad Band Set filter cube for TIRF applications (Chroma). Illumination was provided by an integrated laser engine equipped with multiple laser lines (405nm-100mw/445nm-75mw/488nm-150mw/514nm-40mw/561nm-150mw/637nm-140mw/730nm-40mw, Spectral). Confocal images were captured using a Yokogawa spinning disk and a 405/488/561/647nm Laser Quad Band Set filter cube (Chroma) with a plan apo 63X 1.40 NA objective. Images were acquired using a Hamamatsu ImagEMX2 EM-CCD camera.

In Figure 7, TIRF images were captured using a Leica TIRF-module mounted on a Leica DMi8 equipped with a plan apo 100X 1.47 NA TIRF objective and a Quad Band set filter cube for TIRF applications. Illumination was provided by an integrated laser system equipped with multiple laser lines (405nm-50mw/488nm-150mw/561nm-120mw/638nm-150mW). Images were acquired using LASX software a Hamamatsu Flash 4.0 V3 CMOS camera.

#### Quantification of Microscope PSF

Measurements were performed in 384-well glass-bottomed plates (Brooks) prepared identically to the 3D droplet assays. 5 μL of 0.1 μm diameter TetraSpeck beads (Fisher) were diluted in 120 μL ethanol (final density of ∼ 7.5 x 10^9^ particles/mL). 100 μL was added to a well and incubated for 10 min. Ethanol was removed and the well was rinsed 3X with Buffer A containing 0.1% BSA. 200 μL of Buffer A containing 0.1% BSA was added and images were acquired in all channels with identical acquisition settings to 3D droplet assays. Using spinning disk fluorescence microscopy, Z-stacks of beads were acquired. The PSF was measured for each channel in Image J. For each bead, the Z-plane with the highest intensity was identified (Fig S4a). A linescan through the bead was plotted and the full width half max (FWHM) was determined from a Gaussian fit (Fig S4). For each channel, 20 beads were measured and the mean FWHM was determined. A single plane PSF with the calculated FWHM was generated using the Gaussian PSF 3D ImageJ plugin. Images with circles of known diameter (1 – 50 pixels) were generated. The images were convolved with the PSF. The convolved images were analyzes using MATLAB (Mathworks). A mask was generated by a global image threshold using Otsu’s method. The diameter and mean intensity inside each masked region were measured. The original circle intensity (255) was divided by the measured intensity of the convolved image and plotted against the measured diameter of the convolved image (Fig S4d). The data were fit with a single exponential association. This equation was used to correct the intensity of small droplets.

#### Measuring Droplet Partition Coefficient

Quantitative image analysis was performed using Matlab (Mathworks). Background images were collected and subtracted from all images before processing. To correct for uneven illumination and detector sensitivity, pixel intensities across a solution containing dye were normalized to the maximum intensity of the image to obtain pixel-by-pixel correction factors (in a 0 to 1 range). Experimental images were then corrected by dividing by these factors. A mask of droplets was generated from the 647-channel (Either Nck-Alexa647 or FAK-Alexa647) by a global image threshold using Otsu’s method. The diameter of each masked droplet was measured. For each individual masked droplet, the mean fluorescence intensity inside the masked region was calculated for each channel (Alexa488, Alexa546, Alexa647, although the number of channels in each experiment varied). The mask was dilated and inverted to calculate the mean intensity in the bulk solution (outside of droplets). Droplet intensity was plotted against droplet diameter. If intensity increased with increasing diameter, the intensity was corrected using the PSF correction as in Fig. S4. Droplets with a diameter less than 12 pixels were discarded, since the intensity cannot be accurately measured or corrected. Partition coefficient was calculated for each droplet by dividing droplet mean intensity by the bulk mean intensity.

#### Droplet FRAP measurements

FRAP analysis of droplets was performed using spinning disk confocal microscopy as described above. Droplets with a diameter of ∼4-5 microns were used for FRAP measurements. A 19-pixel (4.8 micron) diameter circular region of interest was drawn around the droplet, and the entire droplet was bleached with 405 laser illumination. Images were acquired at 1s interval for 90 sec. Quantification of droplet intensity was performed using ImageJ. The mean intensity inside the ROI was measured, and an identical sized region in the bulk solution was used to correct for photobleaching during imaging. Background and photobleaching corrected intensities were used for normalization.

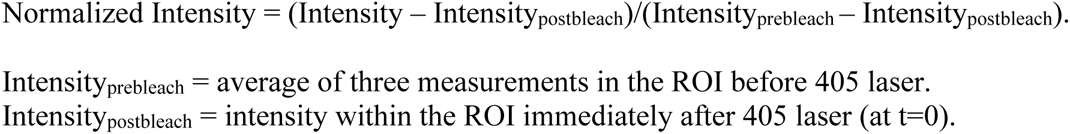

Measurements were repeated at least 8 times, and the curves were averaged and fit using Prism software. To determine the best fit, a single exponential and double exponential fit were statistically compared with an extra sum-of-squares F Test. The values of the best fit are displayed in Table S4. For several molecules, the percent recovery could not be accurately fit from these data, as the recovery did not sufficiently plateau within the 90 second experiment. These are listed as nd.

#### His-kindlin pull-down

His_6_-sumo-kindlin was purified as described above, skipping the Ulp1 cleavage step to keep the his_6_ tag attached. For the pull-down assay, we made a wash Buffer (20 mM Tris pH 7.5, 150 mM NaCl, 10 mM Immidazole) and elution buffer (20 mM Tris pH 7.5, 150 mM NaCl, 300 mM Immidazole). We added 50 µL of resuspended Ni-NTA Resin to Pierce spin columns (Thermo Scientific). Resin was washed 5x with 400 µL of wash buffer, removing buffer with centrifugation at 10,000x g after each wash. A 10 µM solution of his-kindlin was prepared in wash buffer. 100 µL of 10 µM Kin or 100 µL of wash buffer control was added to resin and incubated with gentle shaking for 1 hour at 4°C. After 1 hr, flowthrough was collected by centrifugation at 10,000x g. Resin was washed 5x with 400 uL of wash buffer, removing buffer with centrifugation at 10,000x g after each wash. A 10 µM solution of Cas or pCas was prepared in wash buffer. 100 µL of 10 µM Cas, 10 µM pCas or wash buffer control was added to resin and incubated with gentle shaking for 2 hours at 4°C. The flowthrough was collected by centrifugation at 10,000x g. Resin was washed 5x with 400 µL of wash buffer, removing buffer with centrifugation at 10,000x g after each wash. Finally, 250 µL elution buffer was added to resin and incubated for 5 min at room temperature. Eluted proteins were collected by centrifugation at 10,000x g. The final elution was mixed with equal volume 2X SDS-PAGE loading buffer and boiled for 10 min to denature proteins. Samples were loaded onto a 10% bis-Acrylamide gel for SDS-PAGE (200V for 40 min). Gels were stained with Coomassie Blue and imaged. The following conditions were tested: 1) 10 µM his-kindlin + buffer control; 2) Buffer control + 10 µM Cas; 3) 10 µM his-kindlin + 10 µM Cas; 4) Buffer control + 10 µM pCas; 5) his-kindlin + 10 µM pCas;

#### Cell Culture and Transfection

Mouse Embryonic Fibroblasts (MEFs) were grown in DMEM supplemented with 10% FBS, 2 mM GlutaMAX (Gibco), 100 U/mL penicillin, and 100 µg/mL Streptomycin. FAK +/+ and FAK-/- MEFs were obtained from ATCC. Cas-/- MEFs with Control, CasWT, or CasY15F vectors stably expressed were obtained from Steve Hanks (Vanderbilt) and Larisa Ryzhova (Maine Medical Center). For expression of GFP-tagged Paxillin or FAK variants, cells were transiently transfected with 0.5 μg DNA of EGFP-tagged proteins using lipofectamine 2000 and incubated for 24 hours prior to experiment. For siRNA experiments, cells were transfected using lipofectamine RNAiMAX with pooled oligos (Nontargeting, BCAR1 or PTK2, Dharmacon) and incubated for 48 hours prior to experiment. Details of plasmids used for expression in cells found in Table S6.

#### Quantification of total adhesion area

To determine the transient effect of solvent perturbations on adhesion size and/or number, we quantified total adhesion area after a 10-minute incubation in specified buffers or media. Cells were plated on glass coverslips coated with 10 μg/mL fibronectin and incubated overnight. The experimental buffer or media was added, and cells were incubated for 10 minutes. For temperature experiments, the media was pre-incubated at 4°C, 22°C or 37°C prior to adding to cells. After a 10 min incubation, cells were fixed with 3% paraformaldehyde in Cytoskeleton Buffer (CB: 10 mM MES pH 6.1, 150 mM NaCl, 5 mM EGTA, 5 mM MgCl_2_, 5 mM glucose) for 20 min at room temperature. Cells were then permeabilized for 8 min with CB + 0.5% triton-X, quenched for 10 min with CB + 0.1 M glycine, and rinsed 2 x 5 min in TBS-T. Coverslips were blocked with 2% BSA in TBS-T for 1 hour. Coverslips were incubated with primary antibody (Ms anti Paxillin, 1:100) in TBS-T with 2% BSA overnight at 4°C. Coverslips were rinsed 3 x 5 min in TBS-T and incubated with secondary antibody (anti Ms Alexa568, 1:250) in in TBS-T with 2% BSA for 45 min at room temperature. Coverslips were rinsed 3 x 5 min with TBS-T and mounted on slides with Vectashield (Fischer Scientific, H1000NB). Cells were imaged with spinning disk fluorescence microscopy. To quantify total adhesion area, paxillin images were analyzed in ImageJ. Images were thresholded with a lower threshold level of 5500 and an upper threshold level of 65535. The total thresholded area was then calculated.

#### Adhesion Formation Assay

Cells were trypsinized and plated on glassbottom Matek dishes (5-minute timepoint; #1.5 glass) or glass coverslips (20-minute timepoint; #1.5 glass) coated with 10 μg/mL fibronectin. For a 5-minute time point, unbound cells were gently washed away with fresh media after 1 min. After another 4 min (total spreading time of 5 min), cells were fixed. For a 20-minute time point, unbound cells were gently washed away with fresh media after 10 min. After another 10 min (total spreading time of 20 min), cells were fixed. For all experiments, cells were fixed with 3% paraformaldehyde in Cytoskeleton Buffer (CB: 10 mM MES pH 6.1, 150 mM NaCl, 5 mM EGTA, 5 mM MgCl_2_, 5 mM glucose) for 20 min at room temperature. Cells were then permeabilized for 8 min with CB + 0.5% triton-X, quenched for 10 min with CB + 0.1 M glycine, and rinsed 2 x 5 min in TBS-T. Coverslips were blocked with 2% BSA in TBS-T for 1 hour. Coverslips were incubated with primary antibody (Ms anti Paxillin, 1:100) in TBS-T with 2% BSA overnight at 4°C. Coverslips were rinsed 3 x 5 min in TBS-T and incubated with secondary antibody (anti Ms Alexa568, 1:250) in in TBS-T with 2% BSA for 45 min at room temperature. Coverslips were rinsed 3 x 5 min with TBS-T. For the 5-minute timepoint, PBS was added to the dish and cells were imaged with TIRF microscopy using 561 illumination. For the 20-minute timepoint coverslips were mounted on slides with Vectashield (Fischer Scientific, H1000NB) and imaged with spinning disk fluorescence microscopy. Images in the 488 channel (GFP-tagged proteins) and 561 channel (endogenous paxillin) were acquired).

#### Counting Adhesions

To reduce experimental noise due to differences in expression levels, GFP-transfected cells were only analyzed if the mean GFP intensity within the cell fell within a defined range (intensity between 1000 – 5,000 a.u. following background subtraction, at least 50% of imaged cells were retained with these cutoffs). Adhesions were segmented and counted with ImageJ macros based on a previously published ImageJ workflow (Horzum et al., 2014). First macro: run(“Subtract Background…”, “rolling=50 sliding”); run(“Enhance Local Contrast (CLAHE)”, “blocksize=19 histogram=256 maximum=6 mask=*None* fast_(less_accurate)”); run(“Exp”); run(“Enhance Contrast”, “saturated=0.35”); Then manually run LoG 3D plugin with sigma = 2. Final macro: setAutoThreshold(“Default dark”); setOption(“BlackBackground”, false); run(“Convert to Mask”); run(“Invert”); run(“Analyze Particles…”, “size=5-1000 circularity=0.00-1.00 summarize”). The final mask was compared to the original image to visually confirm the results. We also validated the final results of automated analysis by manually segmenting adhesions (data not shown).

#### Measuring Adhesion Partition Coefficient

Cells expressing GFP-FAK variants or GFP-Paxillin variants were further analyzed to quantify adhesion partitioning of GFP-tagged protein. The same set of images were analyzed for partitioning and counting adhesions. In ImageJ, the GFP image was manually segmented by thresholding using Otsu’s method for dark background. The mean intensity within the threshold was measured. Then the thresholding was manually adjusted to segment the cytoplasm (i.e. exclude the adhesions) and the mean intensity was measured. The background was subtracted from intensity measurements, and the intensity within adhesions was divided by the intensity within the cytoplasm.

#### Western Blot Analysis

Cells were plated on 10 cm round tissue culture dish and grown until ∼75% confluent. Media was removed and plate was gently washed with 1mL cold PBS. 1 mL of cold PBS was added to the dish and cells were scraped for 30 sec. Cells were pelleted with centrifugation at 100g for 4min at 4C. PBS was aspirated and cells were resuspended and lysed in 300 μL cold RIPA buffer (10 mM Tris-HCl (pH 7.6), 1 mM EDTA, 0.1% SDS, 0.1% Na-Deoxycholate, 1% TritonX-100) + cOmplete, EDTA-free Protease Inhibitor tablet (Roche). Lysates were centrifuged for 10min at 10,000 rpm and the supernatant was collected. 100 μL of supernatant was combined with 100 μL of 2X SDS-PAGE loading buffer and boiled for 10min to denature proteins. Samples were loaded onto a 10% bis-Acrylamide gel for SDS-PAGE (200V for 40 min). Samples were transferred onto Immobilon-P PVDF Membrane (90V for 90 min). Membrane was blocked with 5% BSA in TBS-T for 1 hr. Membrane was incubated with primary antibody in TBS-T + 2% BSA overnight at 4°C (Ms anti FAK: 1:1000; Rb anti Cas: 1:1000; Rb anti actin: 1:5000). Membrane was rinsed 3 x 5 min with TBS-T and incubated with secondary antibody 45 min at room temperature (HRP tagged antibodies, 1:10,000). Membrane rinsed 3 x 5 min with TBS-T. Membrane was treated with ECL (Immobilon Western chemiluminescence HRP substrate) for 1 min and visualized with the ChemiDoc MP imaging system (BioRad). Exposure was optimized for each blot.

